# Signaling architectures that transmit unidirectional information despite retroactivity

**DOI:** 10.1101/111971

**Authors:** Rushina Shah, Domitilla Del Vecchio

## Abstract

A signaling pathway transmits information from an upstream system to downstream systems, ideally in a unidirectional fashion. A key obstacle to unidirectional transmission is retroactivity, the additional reaction flux that affects a system once its species interact with those of downstream systems. This raises the fundamental question of whether signaling pathways have developed specialized architectures that overcome retroactivity and transmit unidirectional signals. Here, we propose a general procedure based on mathematical analysis that provides an answer to this question. Using this procedure, we analyze the ability of a variety of signaling architectures to transmit one-way (from upstream to downstream) signals, as key biological parameters are tuned. We find that single stage phosphorylation and phosphotransfer systems that transmit signals from a kinase show a stringent design trade-off that hampers their ability to overcome retroactivity. Interestingly, cascades of these architectures, which are highly represented in nature, can overcome this trade-off and thus enable unidirectional transmission. By contrast, phosphotransfer systems, and single and double phosphorylation cycles that transmit signals from a substrate are unable to mitigate retroactivity effects, even when cascaded, and hence are not well suited for unidirectional information transmission. Our results identify signaling architectures that, allowing unidirectional transmission of signals, embody modular processes that conserve their input/output behavior across multiple contexts. These findings can be used to decompose natural signal transduction networks into modules, and, at the same time, they establish a library of devices that can be used in synthetic biology to facilitate modular circuit design.

## 1 Introduction

Cellular signal transduction is typically viewed as a unidirectional transmission of information via biochemical reactions from an upstream system to multiple downstream systems through signaling pathways [1]–[7]. However, without the presence of specialized mechanisms, signal transmission via chemical reactions is not in general unidirectional. In fact, the chemical reactions that allow a signal to be transmitted from an upstream system to downstream systems also affect the upstream system due to the resulting reaction flux. This flux is called retroactivity, which is one of the chief hurdles to one-way transmission of information [8]–[13]. Signaling pathways, typically composed of phosphorylation, dephosphorylation and phosphotransfer reactions, are highly conserved evolutionarily, such as the MAPK cascade [14] and two-component signaling systems [15]. Thus, the same pathways act between different upstream and downstream systems in different scenarios and organisms, facing different effects of retroactivity in different contexts. For signal transmission to be unidirectional in these different contexts, a signaling pathway should have evolved architectures that overcome retroactivity. Specifically, these architectures should impart a small retroactivity to their upstream system (called retroactivity to the input) and should be minimally affected by the retroactivity imparted to them by their downstream systems (retroactivity to the output).

Phosphorylation-dephosphorylation cycles, phosphotransfer reactions, and cascades of these are ubiquitous in both prokaryotic and eukaryotic signaling pathways, playing a major role in cell cycle progression, survival, growth, differentiation and apoptosis [1]–[7], [16]–[19]. Numerous studies have been conducted to analyze such systems, starting with milestone works by Stadtman and Chock, [20], [21], [22] and Goldbeter et al. [23], [24], [25], which theoretically and experimentally analyzed phosphorylation cycles and cascades. These systems were further investigated by Kholdenko et al. [26], [27], [28] and Gomez-Uribe et al. [29], [30]. However, these studies considered signaling cycles in isolation, and thus did not investigate the effect of retroactivity. The effect of retroactivity on such systems was theoretically analyzed in the work by Ventura et al. [31], where retroactivity is treated as a “hidden feedback” to the upstream system. Experimental studies then confirmed the effects of retroactivity in signaling systems through *in vivo* experiments on the MAPK cascade [12], [13] and *in vitro* experiments on reconstituted covalent modification cycles [9], [11]. These studies clearly demonstrated that the effects of retroactivity on a signaling system manifest themselves in two ways. They cause a slow down of the temporal response of the signaling system’s output to its input and lead to a change of the output’s steady state.

In 2008, Del Vecchio et al. demonstrated theoretically that a single phosphorylation-dephosphorylation (PD) cycle with a slow input kinase can attenuate the effect of retroactivity to the output when the total substrate and phosphatase concentrations of the cycle are increased together [8]. Essentially, a sufficiently large phosphatase concentration along with relatively large kinetic rates of modification adjusts the cycle’s internal dynamics very quickly with respect to a relatively slower input, making any retroactivity-induced delays negligible on the time scale of the signal being transmitted [32]. A similarly large concentration of total cycle’s substrate ensures that the output’s steady state is not significantly affected by the presence of downstream sites. These theoretical findings were later verified experimentally both *in vitro* [11] and *in vivo* [33]. Although a single PD cycle can attenuate the effect of retroactivity to the output, it is unfortunately unsuitable for unidirectional signal transmission. In fact, as the substrate concentration is increased, the PD cycle applies a large retroactivity to the input, causing the input signal to slow down. This was experimentally observed in [33]. The experimental results of [34] further suggest that a cascade composed of two PD cycles and a phosphotransfer reaction could overcome both retroactivity to the input and retroactivity to the output. In [35], it was theoretically found that, for certain parameter conditions, a cascade of PD cycles could attenuate the upward (from downstream to upstream) propagation of disturbances applied downstream of the cascade. These results suggest that specific signaling architectures may be able to counteract retroactivity. However, to the best of the authors’ knowledge, no attempt has been made to systematically characterize signaling architectures with respect to their ability to overcome the effects of retroactivity and therefore enable unidirectional signal transmission.

This work presents a procedure to identify and characterize signaling architectures that can transmit unidirectional signals. This procedure is based on mathematical analysis of a reaction-rate ordinary differential equation (ODE) model for a general signaling system that operates on a fast timescale relative to its input. Such a model is valid for many signaling systems that transmit relatively slower signals, such as those from slowly varying “clock” proteins that operate on the timescale of the circadian rhythm [36], from proteins signaling nutrient availibility [37], or from proteins whose concentration is regulated by transcriptional networks, which operate on the slower timescale of gene expression [38]. Our framework provides expressions for retroactivity to the input and to the output as well as the input-output relationship of the signaling system. These expressions are given in terms of the reaction-rate parameters and protein concentrations. Based on these expressions, we present a procedure to analyze the ability of signaling systems to transmit unidirectional signals by tuning their total (modified + unmodified) protein concentrations. We focus on total protein concentrations as a design parameter because these appear to be highly variable in natural systems and through the course of evolution, where they may have been optimized to improve systems' performance [39], [40]. Protein concentration is also an easily tunable quantity in synthetic genetic circuits. We use the procedure we have presented to analyze a number of signaling architectures composed of PD cycles and phosphotransfer systems.

## 2 Methods

### 2.1 Problem Definition

In this work, we consider a general signaling system **S** connected between an upstream and downstream system, as shown in Fig. 1A. Here, *<u>X</u>* is the state-variable vector of **S**, and each component of *<u>X</u>* represents the concentration of a species of system **S**. System **S** receives an input from the upstream system in the form of a protein whose concentration is *U*, and sends an output to the downstream system in the form of a protein whose concentration is *Y*. When this output protein reacts with the species of the downstream system, whose normalized concentrations are represented by state variable *υ*, the resulting reaction flux changes the behavior of the upstream system. We represent this reaction flux as an additional input, 𝒮, to the signaling system. Similarly, when the input protein from the upstream system reacts with the species of the signaling system, the resulting reaction flux changes the behavior of the upstream system. We represent this as an input, ℛ, to the upstream system. We call ℛ the retroactivity to the input of **S** and 𝒮 the retroactivity to the output of **S**, as in [8]. For system **S** to transmit a unidirectional signal, the effects of ℛ on the upstream system and of 𝒮 on the downstream system must be small. Retroactivity to the input R changes the input from *U*_ideal_ to *U*, where *U*_ideal_ is shown in Fig. 1B. Thus, for the effect of ℛ to be small, the difference between *U* and *U*_ideal_ must be small. Retroactivity to the output 𝒮 changes the output from *Y*_is_ (where “is” stands for isolated) to *Y*, where *Y*_is_ is shown in Fig 1C, and for the effect of retroactivity to the output to be small, the difference between *Y*_is_ and *Y* must be small. An *ideal unidirectional signaling system* is therefore a system where the input *U*_ideal_ is transmitted from the upstream system to the signaling system without any change imparted by the latter, and the output *Y*_is_ of the signaling system is also transmitted to the downstream system without any change imparted to it by the downstream system. Based on this concept of ideal unidirectional signaling system, we then present the following definition of a signaling system that can transmit information unidirectionally. In order to give this definition, we assume that the proteins (besides the input species) that compose signaling system **S** are constitutively produced and therefore their total concentrations (modified and unmodified) are constant. The vector of these total protein concentrations is denoted by <u>Θ</u>.

**Figure 1:**
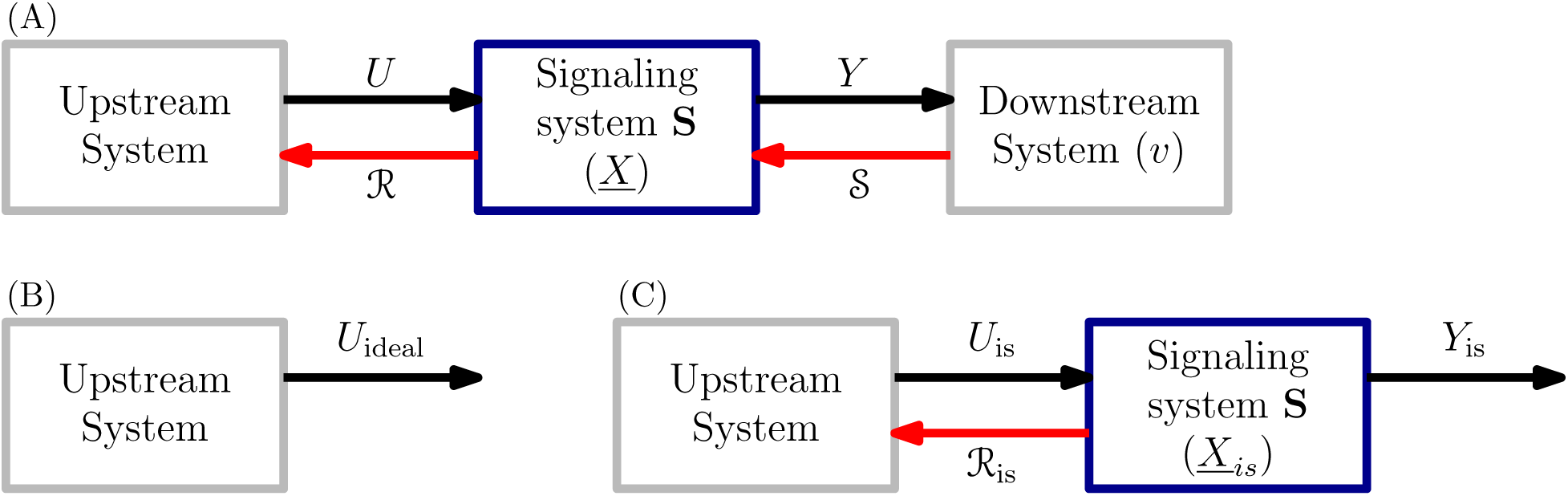
Interconnections between a signaling system S and its upstream and downstream systems, along with input, output and retroactivity signals. (A) Full system showing all interconnection signals: *U* (*t*) is the input from the upstream system to the signaling system, with state variable vector *<u>X</u>*. *Y* (*t*) is the output of the signaling system, sent to the downstream system, whose state variable is *υ*. ℛ is the retroactivity signal from the signaling system to the upstream system (retroactivity to the input of **S**), and 𝒮 is the retroactivity signal from the downstream system to the signaling system (retroactivity to the output of **S**). (B) Ideal input *U*_ideal_: output of the upstream system in the absence of the signaling system (ℛ = 0). (C) Isolated output *Y*_is_: output of the signaling system in the absence of the downstream system (𝒮 = 0). *<u>X</u>*_is_ denotes the corresponding state of **S**.

#### Definition 1

We will say that system **S** is a signaling system that can transmit unidirectional signals for all inputs *U ∈* [0, *U_b_*], if <u>Θ</u> can be chosen such that the following properties are satisfied:

(i) ℛ is small: this is mathematically characterized by requiring that |*U*_ideal_ (*t*) − *U* (*t*)| be small for all *U ∈* [0, *U_b_*].
(ii) System **S** attenuates the effect of 𝒮 on *Y*: this is mathematically characterized by requiring that |*Y*_is_(*t*) − *Y*(*t*)| be small for all *U ∈* [0, *U_b_*].
(iii) Input-output relationship: *Y*_is_(*t*) ≈ *KU_is_*(*t*)*^m^*, for some *m* ≥ 1, for some *K* > 0 and for all *U ∈* [0, *U_b_*].

Note that Def. 1 specifies that the signaling system must impart a small retroactivity to its input (i) and attenuate retroactivity to its output (ii). In particular, it specifies that these properties should be satisfied for a full range of inputs and outputs, implying that these properties must be guaranteed by the features of the signaling system and cannot be enforced by tuning the amplitudes of inputs and/or outputs.

### 2.2 Example

As an illustrative example of the effects of ℛ and 𝒮 on a signaling architecture, we consider a signaling system **S** composed of a single PD cycle [8], [11], [33]. The system is shown in Fig. 2A. It receives a slowly varying input signal *U* in the form of kinase concentration *Z* generated by an upstream system, and has as the output signal *Y* the concentration of X^*^, which in this example is a transcription factor that binds to promoter sites in the downstream system. Kinase Z phosphorylates protein X to form X^*^, which is dephosphorylated by phosphatase M back to X. The state variables *<u>X</u>* of **S** are the concentrations of the species in the cycle, that is, *X, M, X*^*^, *C*_1_, *C*_2_, where C_1_ and C_2_ are the complexes formed by X and Z during phosphorylation, and by X^*^ and M during dephosphorylation, respectively. The state variable *υ* of the downstream system is the normalized concentration of C, the complex formed by X^*^ and p (i.e., 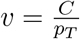 where *p_T_* is the total concentration of the downstream promoters). This configuration, where a signaling system has as downstream system(s) gene expression processes, is common in many organisms as it is often the case that a transcription factor goes through some form of covalent modification before activating or repressing gene expression [41]. However, the downstream system could be any other system, such as another covalent modification process, which interacts with the output through a binding-unbinding reaction. We denote the total amount of cycle substrate by *X_T_* = *X* + *X*^*^ + *C*_1_ + *C*_2_ + *C* and the total amount of phosphatase by *M_T_* = *M* + *C*_2_.

**Figure 2:**
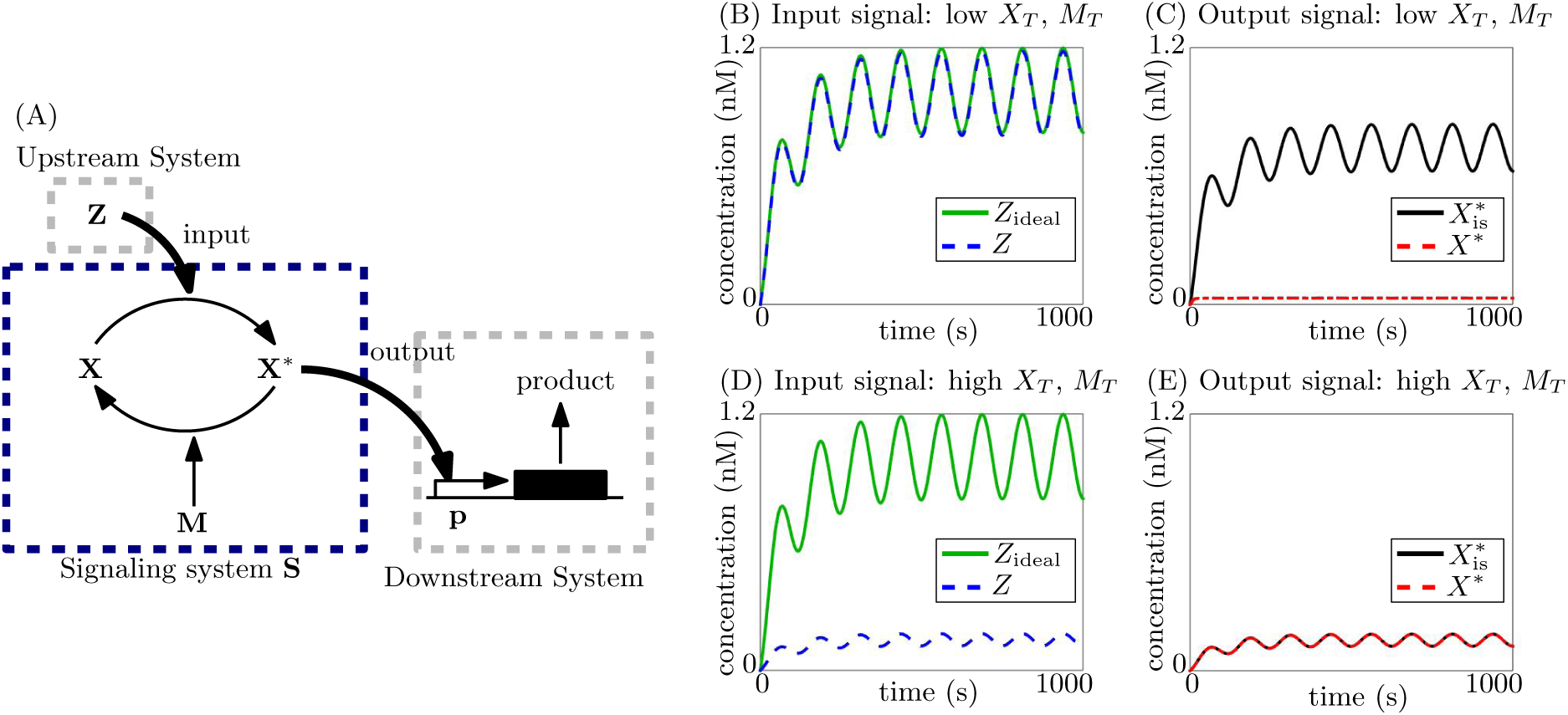
Tradeoff between small retroactivity to the input and attenuation of retroactivity to the output in a single phosphorylation cycle. (A) Single phosphorylation cycle, with input Z as the kinase: X is phosphorylated by Z to X^*^, and dephosphorylated by the phosphatase M. X^*^ is the output and acts on sites p in the downstream system, which is depicted as a gene expression system here. (B)-(E) Simulation results for ODE model shown in SI Section 5.3 eqn. (22). Simulation parameters are given in Table 1 in SI Section 5.2. Ideal system is simulated for *Z*_ideal_ with *X_T_* = *M_T_* = *p_T_* = 0. Isolated system is simulated for 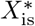 with *p_T_* = 0.

According to Def. 1, we vary the total protein concentrations of the cycle, <u>Θ</u> = [*X_T_*, *M_T_*], to investigate the ability of this system to transmit unidirectional signals. To this end, we consider two extreme cases: first, when the total substrate concentration *X_T_* is low (simulation results in Figs. 2B, 2C); second, when it is high (simulation results in Figs. 2D, 2E). For both these cases, we change *M_T_* proportionally to *X_T_.* This is because, for large Michaelis-Menten constants, we have an input-output relationship with *m* = 1 and 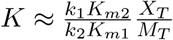 (details in SI Section 5.3, eqn. (26)) as defined in Def. 1(iii). To maintain the same *K* for fair comparison between the two cases, we vary *M_T_* proportionally with *X_T_*. Here, *K*_*m*1_ and *k*_1_ are the Michaelis-Menten constant and catalytic rate constant for the phosphorylation reaction, and *K*_*m*2_ and *k*_2_ are the Michaelis-Menten constant and catalytic rate constant for the dephosphorylation reaction. These reactions are shown in (eqns. 21) in SI Section 5.3. For the simulation results, we consider a sinusoidal input to see the dynamic response of the system to a time-varying signal. For these two cases then, we see from Fig. 2B that when *X_T_* (and *M_T_*) is low, ℛ is small, i.e., |*U*_ideal_(*t*) — *U*(*t*)| is small (satisfying requirement (i) of Def. 1). This is because kinase Z must phosphorylate very little substrate X, and thus, the reaction flux due to phosphorylation to the upstream system is small. However, as seen in Fig. 2C, for low *X_T_*, the signaling system is unable to attenuate 𝒮. The difference 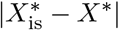 is large, and requirement (ii) of Def. 1 is not satisfied for low *X_T_*. This large retroactivity to the output is due to the reduction in the total substrate available for the cycle because of the sequestration of X^*^ by the promoter sites in the downstream system. Since *X_T_* is low, this sequestration results in a large relative change in the amount of total substrate available for the cycle, and thus interconnection to the downstream system has a large effect on the behavior of the cycle. For the case when *X_T_* (and *M*_T_) is high, the system shows exactly the opposite behavior. From Fig. 2D, we see that ℛ is high (thus not satisfying requirement (i) of Def. 1), since the kinase must phosphorylate a large amount of substrate, but 𝒮 is attenuated (satisfying requirement (ii)) since there is enough total substrate available for the cycle even once X^*^ is sequestered. Thus, this system shows a trade-off: by increasing *X_T_* (and *M_T_*) we attenuate retroactivity to the output but do so at the cost of increasing retroactivity to the input. Similarly, by decreasing *X_T_* (and *M_T_*), we make retroactivity to the input smaller, but at the cost of being unable to attenuate retroactivity to the output. Therefore, requirements (i) and (ii) cannot be independently obtained by tuning *X_T_* and *M_T_*.

We note that because the signaling reactions, i.e., phosphorylation and dephosphorylation, act on a faster timescale than the input, the signaling system operates at quasi-steady state and the output is able to quickly catch up to changes in the input. It has been demonstrated in [32], [34] that this fast timescale of operation of the signaling system attenuates the temporal effects of retroactivity to the output, which would otherwise result in the output slowing down in the presence of the downstream system. Thus, while the high substrate concentration *X_T_* is required to reduce the effect of retroactivity to the output due to permanent sequestration, timescale separation is necessary for attenuating the temporal effects of the binding-unbinding reaction flux [32].

### 2.3 Generalized model

The single phosphorylation cycle, while showing some ability to attenuate retroactivity, is not able to transmit unidirectional signals due to the trade-off seen above. We therefore study different architectures of signaling systems, composed of phosphorylation cycles and phosphotransfer systems which are ubiquitous in natural signal transduction [1]–[7], [14]–[19]. All reactions are modeled as two step reactions. Phosphorylation and dephosphorylation reactions are modeled as Michaelis-Menten reactions, and phosphotransfer reactions are modeled as reversible, two-step reactions resulting in the transfer of the phosphate group via the formation of an intermediate complex. Based on these reactions, as well as production and decay of the various species, ODE models are created for the systems using their reaction-rate equations. The following general ODE model then describes any signaling system architecture in the interconnection topology of Fig. 1A:

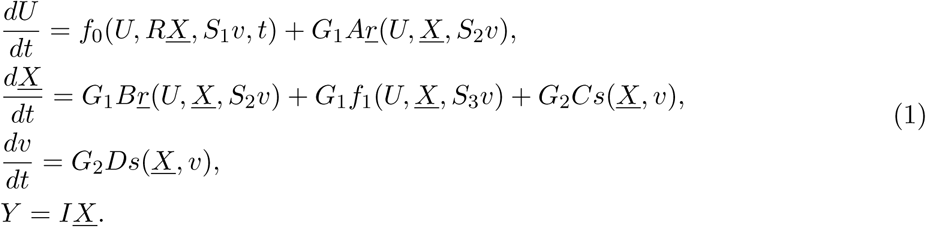

Here, the variable *t* represents time, *U* is the input signal (the concentration of the input species), *<u>X</u>* is a vector of concentrations of the species of the signaling system, *Y* is the output signal (the concentration of the output species) and *υ* is the state variable of the downstream system. In the cases that follow, *υ* is the normalized concentration of the complex formed by the output species Y and its target binding sites p in the downstream system.

The internal dynamics of the upstream system are captured by the reaction-rate vector *f*_0_. The internal dynamics of the signaling system are captured by the reaction-rate vector *f*_1_. The reaction-rate vector *<u>r</u>* is the reaction flux resulting from the reactions between species of the upstream system and those of the signaling system, with corresponding stoichiometric matrices *A* and *B.* The reaction rate vector *s* represents the additional reaction flux due to the binding-unbinding of the output protein with the target sites in the downstream system, with corresponding stoichiometric matrices *C* and *D.* These additional reaction fluxes affect the temporal behavior of the input and the output, often slowing them down, as demonstrated previously [11].

The parameter *R* accounts for decay/degradation of complexes formed by the input species with species of the signaling system, thus leading to an additional channel for removal of the input species through their interaction with the signaling system. Similarly, scalar *S*_1_ represents decay of complexes formed by the input species with species of the downstream system. This additional decay leads to an effective increase in decay of the input, thus affecting its steady-state. As species of the signaling system are sequestered by the downstream system, their free concentration changes. This is accounted for by the vectors *S*_2_ and *S*_3_.

The retroactivity to the input ℛ indicated in Fig. 1A therefore equals (*R, <u>r</u>, S*_1_), which leads to both steady-state and temporal effects on the input response. The retroactivity to the output 𝒮 of Fig. 1A equals (*S*_1_, *S*_2_, *S*_3_, *s*), which leads to an effect on the output response. For ideal unidirectional signal transmission, the effects of ℛ and 𝒮 must be small. The ideal input of Fig. 1B, *U*_ideal_, is the input when retroactivity to the input ℛ is zero, i.e., when *R* = *S*_1_ = *<u>r</u>* = 0. The isolated output of Fig. 1C, *Y*_is_, is the output when retroactivity to the output 𝒮 is zero, i.e., when *S*_1_ = *S*_2_ = *S*_3_ = *s* = 0.

The positive scalar *G*_1_ captures the timescale separation between the reactions of the signaling system and the dynamics of the input. Since we consider relatively slow inputs, we have that *G*_1_ ≫ 1. The positive scalar *G*_2_ captures the timescale separation between the binding-unbinding rates between the output Y and its target sites p in the downstream system and the dynamics of the input. Since binding-unbinding reactions also operate on a fast timescale, we have that *G*_2_ ≫ 1. We define 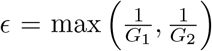 and thus, E ≪ 1. This allows us to apply techniques from singular perturbation to simplify and analyze model (1), to arrive at the results presented in the next section. Details of this analysis are shown in SI Section 5.1.

In Section 3, we outline a procedure to determine if a given signaling system satisfies Def. 1. For this, we introduce the following definitions. We assume that there exist matrices *M* and *P*, and invertible matrices *T* and Q such that:

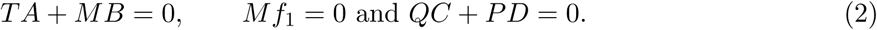

This assumption is usually satisfied in signaling systems [32]. Further, we have:

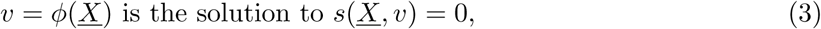

and

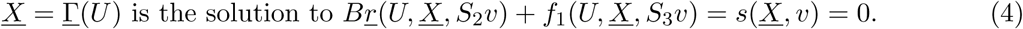

We note that, for system (1), terms *S*_2_, *S*_3_, functions *f*_1_ and <u>Γ</u> depend on the vector of total protein concentrations, <u>Θ</u>.

### 2.4 Simulations

Our theoretical analysis for the various systems is verified using simulations of the full ODE systems run on MATLAB, using the numerical ODE solvers ode23s and ode15s. All simulation parameters are picked from the biologically relevant ranges given in [35], and are listed in Table 1 in SI Section 5.2.

## 3 Results

The main result if this paper is two-fold. First, we provide a general procedure to determine whether any given signaling system enables unidirectional signal transmission. Second, using this procedure, we analyze the unidirectional signal transmission ability of both common and less frequent signaling architectures.

### 3.1 Procedure to determine unidirectional signal transmission

We outline a procedure to determine whether any given signaling system can enable unidirectional signaling in Fig. 3. First, the reaction-rate equations of the signaling system are written in form (1), allowing us to note the terms *S*_1_, *S*_2_, *S*_3_ and *R* for Step 2. The remaining terms for Step 2 are computed using equations (2)-(4) in Section 2.3. The terms in Steps 3, 4 and 5 are computed using the terms in Step 2. The upperbound on | *U*(*t*) − *U*_ideal_(*t*) | is proportional to the terms found in Step 3, and thus, as these are made small according to Test (i), Def. 1(i) is satisfied. The analysis giving rise to these terms is shown in Theorem 1 in SI Section 5.1. Similarly, the upperbound on |*Y_is_(t*) − *Y*(*t*)| is proportional to the terms in Step 4, and thus, as these are made small according to Test (ii), Def. 1(ii) is satisfied. This is derived in Theorem 2 in SI Section 5.1. Theorem 3 in SI Section 5.1 shows that the input-output relationship for the signaling system can be computed by Step 5. If this input-output relationship satisfies Test (iii), Def. 1(iii) is satisfied. Once Tests (i)-(iii) are satisfied, Test (iv) checks if all the requirements for Def. 1 can be achieved simultaneously by tuning <u>Θ</u>. If this is possible, the signaling system is said to be able to transmit a unidirectional signal.

**Figure 3:**
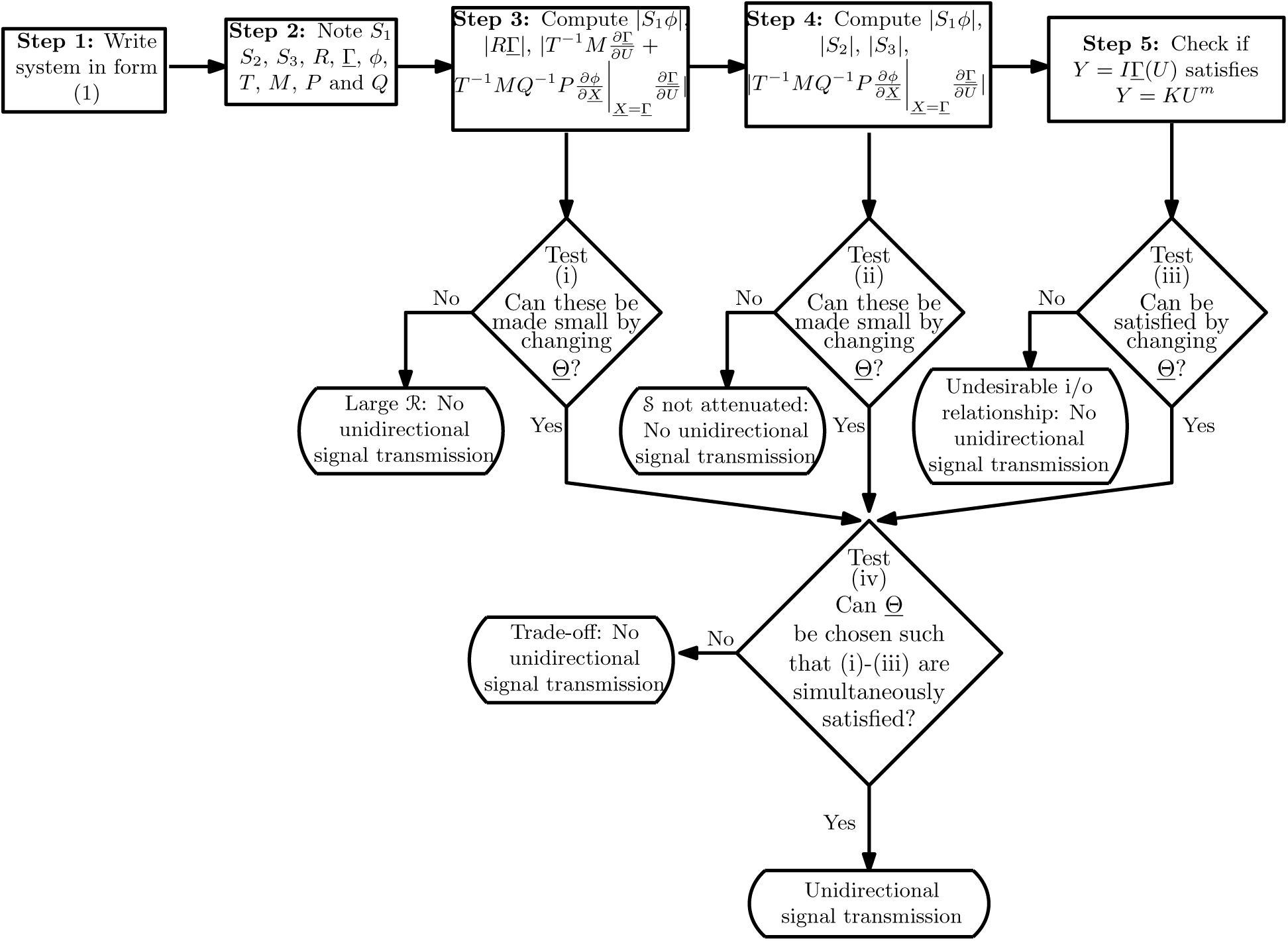
Procedure to determine if a given signaling system satisfies Def. 1 for unidirectional signal transmission.

As an example of the application of the procedure, we consider once again the single PD cycle of Section 2.2. Steps 1-5 for this system are shown in SI Section 5.3. We find that in order to satisfy Test (i), we must have small *X_T_*. Further, to satisfy Test (ii), we must have large *X_T_* and *M_T_*. Finally, computing *I*<u>Γ</u> from Step 5, we find that the input-output relationship has 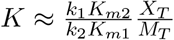 with *m* = 1 when *K*_*m*1_, *K_m2_* ≫ 1. These results are consistent with those described in Section 2.2 as well as previous theoretical and experimental work [8], [11], [33]. We see that there exists a trade-off between (i) and (ii), i.e., between imparting a small retroactivity to the input and attenuating retroactivity to the output. Thus, <u>Θ</u> cannot be chosen such that all 3 requirements are simultaneously satisfied. Test (iv) fails, and the single PD cycle cannot achieve unidirectional signal transmission.

This way, the above procedure can be used to identify ways to tune the total protein concentration of a signaling system such that it satisfies Def. 1. Using this procedure, we analyze a number of signaling architectures, including double phosphorylation systems, phosphotransfer systems, and multi-stage signaling architectures composed of these. For these architectures, we consider two types of input signals: a kinase input (highly represented in natural systems), where the input regulates the rate of phosphorylation, and a substrate input (less frequent in natural systems), where the input regulates the rate of production of the substrate.

### 3.2 Double phosphorylation cycle with input as kinase

Here, we consider a double phosphorylation cycle with a common kinase Z for both phosphorylation cycles as the input and the doubly phosphorylated substrate X^**^ as the output. This architecture is found in the second and third stages of the MAPK cascade, where the kinase phosphorylates both the threonine and tyrosine sites in a distributive process [42]. This configuration is shown in Fig. 4A. Referring to Fig. 1A, the input signal *U* is the concentration *Z* of the kinase and the output signal *Y* is the concentration *X^**^* of the doubly phosphorylated substrate X.

**Figure 4:**
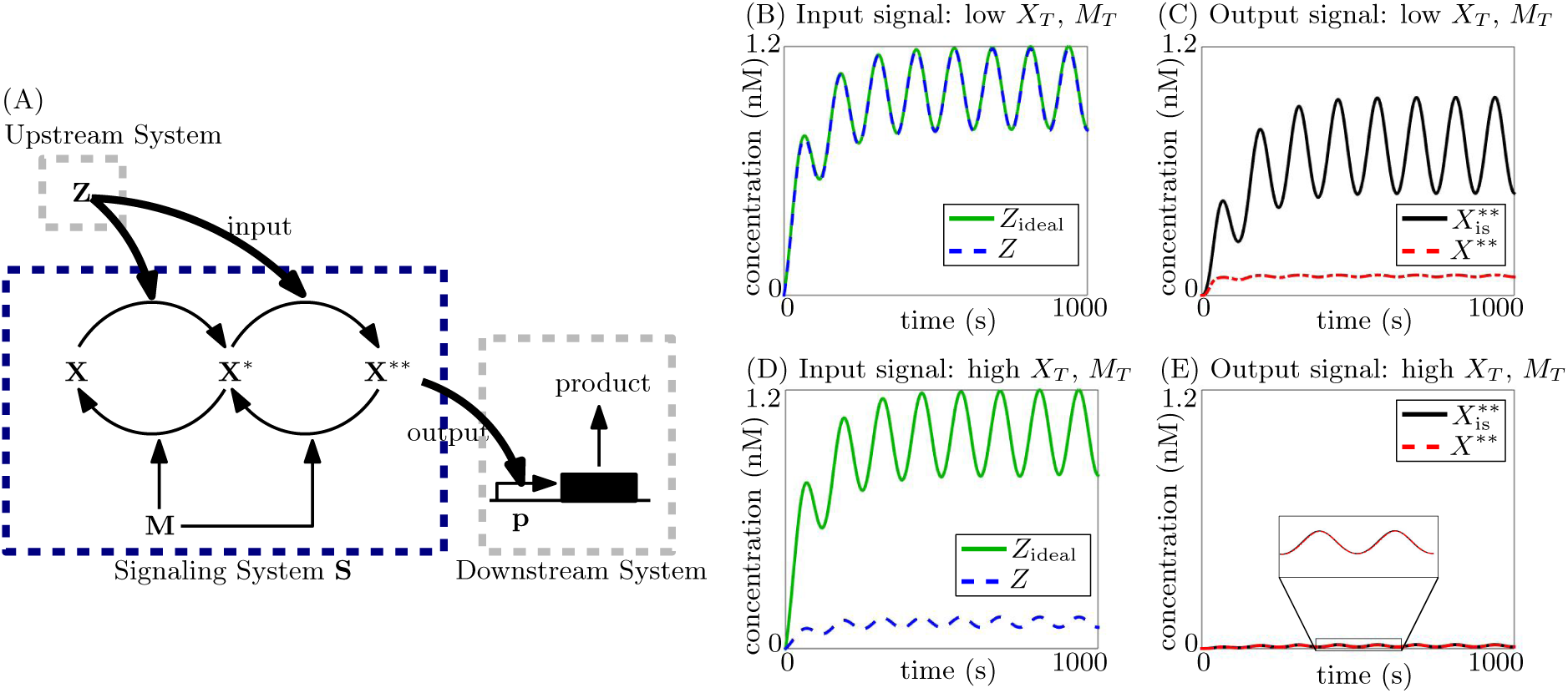
Tradeoff between small retroactivity to the input and attenuation of retroactivity to the output in a double phosphorylation cycle. (A) Double phosphorylation cycle, with input Z as the kinase: X is phosphorylated by Z to X^*^, and further on to X^**^. Both these are dephosphorylated by the phosphatase M. X^**^ is the output and acts on sites p in the downstream system, which is depicted as a gene expression system here. (B)-(E) Simulation results for ODE model (34) shown in SI Section 5.4. Simulation parameters are given in Table 1 in SI Section 5.2. The ideal system is simulated for *Z*_ideal_ with *X*_T_ = *M*_T_ = *p_T_* = 0. The isolated system for 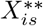 is simulated with *p_T_* = 0.

The input kinase is produced at a time-varying rate *k*(*t*). All species dilute with a rate constant *δ*, and the total promoter concentration in the downstream system is *p_T_*. The total substrate and phosphatase concentrations are *X_T_* and *M_T_*, respectively. The Michaelis-Menten constants for the two phosphorylation and the two dephosphorylation reactions are *K*_*m*1_, *K*_*m*3_, *K*_*m*2_ and *K*_*m*4_, respectively. The catalytic reaction rate constants of these reactions are *k*_1_, *k*_3_, *k*_2_ and *k*_4_, respectively. The system’s chemical reactions are shown in SI Section 5.4 eqns. (33). As explained before, the parameters that we tune to investigate retroactivity effects are the total protein con-centrations of the phosphorylation cycle, that is, *X_T_* and *M_T_*. Specifically, using the procedure in Fig. 3, we tune *X*_T_ and *M_T_* to verify if this system can transmit a unidirectional signal, according to Definition 1. Steps 1-5 are detailed in SI Section 5.4. We therefore find what follows.

(i) Retroactivity to the input: Evaluating the terms in Step 3, we find that to satisfy Test (i), we must have small and small 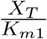 and small 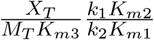. Thus, to have small retroactivity to the input, the parameter *X*_T_ must be small. (Evaluation of terms in Step 3 is shown in SI Section 5.4).
(ii) Retroactivity to the output: Evaluating the terms in Step 4, we find that to satisfy Test (ii), we must have small 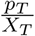 and 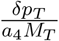. Thus, to attenuate retroactivity to the output, we must have large *X_T_* and *M_T_*. (Evaluation of terms in Step 4 is shown in SI Section 5.4).
(iii) Input-output relationship: Computing *I*<u>Γ</u> find that 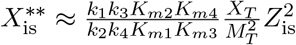 when *K*_*m*1_, *K*_*m*2_, *K*_*m*3_, *K*_*m*4_ ≫ *Z*_is_, 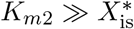, 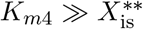 and *M_T_* ≫ *Z*_is_. Under these assumptions, this system satisfies Test (iii) by tuning the ratio 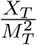 to achieve a desired *K* with *m* = 2. (Evaluation of Step 5 is shown in SI Section 5.4).

This system shows opposing requirements to satisfy Tests (i) and (ii), similar to the single phosphorylation cycle. Thus, while each of the requirements of Tests (i)-(iii) are individually satisfied, the system does not satisfy Test (iv), showing a trade-off that prevents unidirectional signal transmission. Retroactivity to the input is large when substrate concentration *X*_T_ (and *M_T_*) increases, because the input Z must phosphorylate a large amount of substrate thus leading to a large reaction flux to Z due to the phosphorylation reaction. However, if *X*_T_ (and *M*_T_) is made small, the system cannot attenuate the retroactivity to the input, since as the output X^**^ is sequestered by the downstream system, there is not enough substrate available for the signaling system. Therefore, Tests (i) and (ii) cannot be independently satisfied.

These mathematical predictions can be appreciated from the numerical simulations of Figs. 4B-4E and this result is summarized in Fig. 10B.

### 3.3 Regulated autophosphorylation followed by phosphotransfer

We now consider a signaling system composed of a phosphotransfer system, whose phosphate donor receives the phosphate group via autophosphorylation regulated by protein Z. An instance of this architecture is found in the bacterial chemotaxis network, where the autophosphorylation of protein CheA is regulated by a transmembrane receptor (e.g., Tar). CheA then transfers the phosphate group to protein CheY in a phosphotransfer reaction. CheY further undergoes dephosphorylation catalyzed by the phosphatase CheZ [43], [44], [45]. A similar mechanism is also present in the ubiquitous two-component signaling networks, where the sensor protein autophosphorylates upon binding to a stimulus (e.g., a ligand) and then transfers the phosphate group to the receptor protein [46], [15]. We model this regulated autophosphorylation as a phosphorylation reaction with kinase as input, since in both cases, first an intermediate complex is formed and the protein then undergoes phosphorylation. This architecture is shown in Fig. 5A. In this case, the input signal *U* of Fig. 1A is *Z*, which is the concentration of the kinase/stimulus Z that regulates the phosphorylation of the phosphate donor X_1_, which then transfers the phosphate group to protein X_2_. The output signal *Y* in Fig. 1A is then 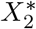 which is the concentration of the phosphorylated substrate 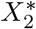. Protein 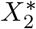 is dephosphorylated by phosphatase M. Total concentrations of proteins X_1_, X_2_ and M are *X*_*T*1_, *X*_*T*2_ and *M_T_*, respectively. The Michaelis-Menten constants for the phosphorylation of X_1_ by Z and dephosphorylation of 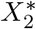 by M are *K*_*m*1_ and *K_m3_*, and the catalytic rate constants of these are *k*_***1***_ and *k*_3_, respectively. The association rate constant of complex formation by 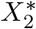 and X_1_ is *a_3_.* These reactions are shown in eqns. (53) in SI Section 5.5. The total concentration of promoter sites in the downstream system is *p_T_*. The input Z is produced at a time-varying rate *k*(*t*). As before, the parameters we change to analyze the system for unidirectional signal transmission are its total protein concentrations, *X*_*T*1_, *X*_*T*2_ and *M_T_*. Using the procedure in Fig. 3, we analyze the system’s ability to transmit unidirectional signals as per Definition 1 as *X*_*T*1_, *X*_*T*2_ and *M_T_* are varied. This is done as follows. (Steps 1-5 for this system are shown in SI Section 5.5).

(i) Retroactivity to the input: Evaluating the terms in Step 3, we find that to satisfy Test (i), we must have small 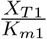. Thus, for small retroactivity to the input, we must have small *X*_*T*1_. (Evaluation of terms in Step 3 is shown in SI Section 5.5).
(ii) Retroactivity to the output: Evaluating the terms in Step 4, we find that to satisfy Test (ii), 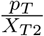 and 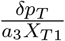 must be small. Thus, for a small retroactivity to the output, we must have large *X*_*T*1_ and *X*_*T*2_. (Evaluation of terms in Step 4 is shown in SI Section 5.5).
(iii) Input-output relationship: Evaluating *I*<u>T</u> as in Step 5, we find that 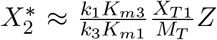 when *K*_*m*1_ ≫ *Z*_is_ and 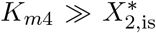. Under these assumptions, this system satisfies Test (iii), where a desired *K* can be achieved by tuning the ration 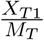 with *m* = 1. (Evaluation of Step 5 is shown in SI Section 5.5).

**Figure 5:**
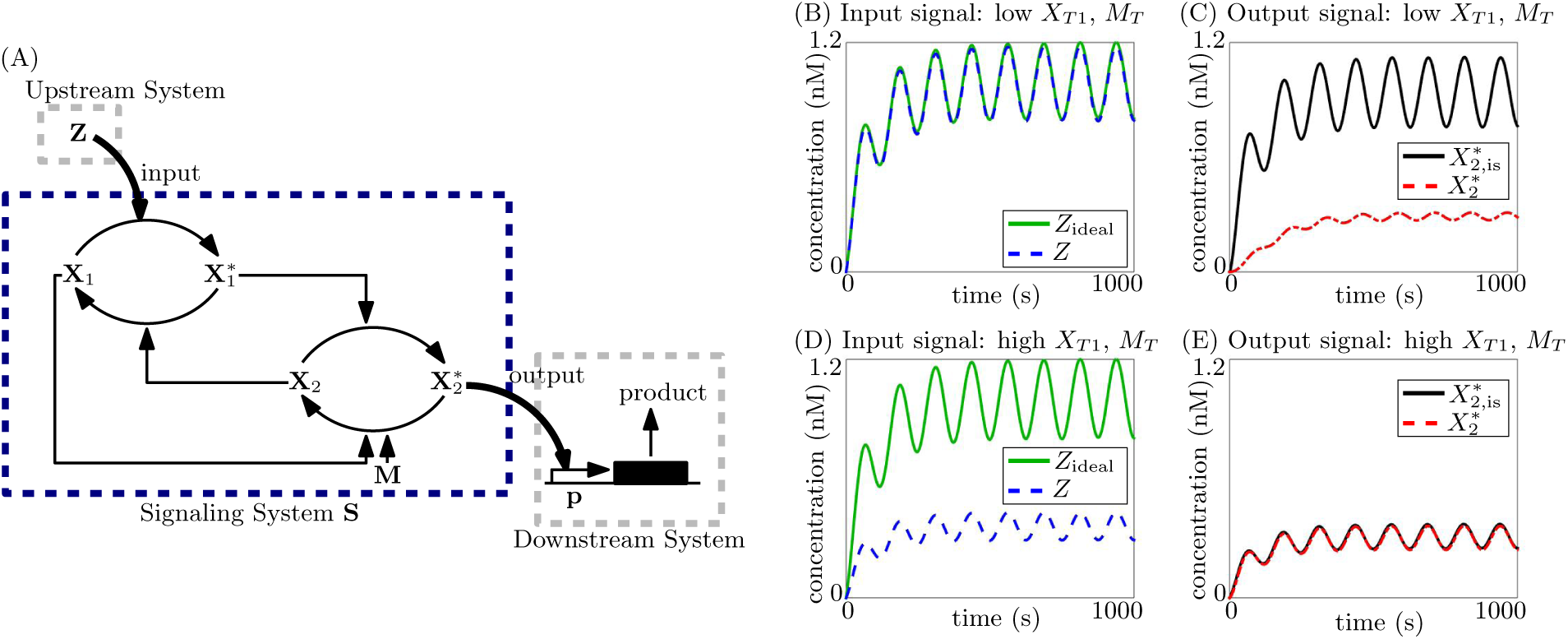
Tradeoff between small retroactivity to the input and attenuation of retroactivity to the output in a phosphotransfer system. (A) System with phosphorylation followed by phosphotransfer, with input Z as the kinase: Z phosphorylates X_1_ to 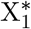. The phosphate group is transferred from 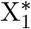 to X_2_ by a phosphotransfer reaction, forming 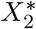, which is in turn dephosphorylated by the phosphatase M. 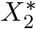 is the output and acts on sites p in the downstream system, which is depicted as a gene expression system here. (B)-(E) Simulation results for ODE (54) in SI Section 5.5. Simulation parameters are given in Table 1 in SI Section 5.2. Ideal system is simulated for *Z*_ideal_ with *X*_*T*1_ = *X*_*T*2_ = *M_T_* = *p_T_* = 0. Isolated system is simulated for 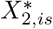 with *p_T_* = 0.

In light of (i) and (ii), we note that Tests (i) and (ii) cannot be simultaneously satisfied. Test (iv) fails, and the system shows a trade-off in attenuating retroactivity to the input and output. Retroactivity to the input can be made small, by making *X*_*T*1_ (and *M_T_*) small, since kinase Z must phosphorylate less substrate. However, the system with low *X*_*T*1_ is unable to attenuate retroactivity to the output, which requires that *X*_*T*1_ be large. This is because, as the output 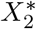 is sequestered by the downstream system and undergoes decay as a complex, this acts as an additional channel of removal for the phosphate group from the system, which was received from 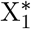. If *X*_*T*1_ (and *M_T_*) is small, this removal of the phosphate group affects the amount of 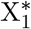 in the system to a larger extent that when *X*_*T*1_ is large. Thus, there exists a trade-off between requirements (i) and (ii) of Def. 1, and the system does not allow unidirectional signal transmission.

This mathematical analysis is demonstrated in the simulation results shown in Figs. 5B-5E and the discussion is summarized in Fig. 10B.

### 3.4 Cascade of single phosphorylation cycles

We have now seen three systems that show a trade-off between attenuating retroactivity to the output and imparting a small retroactivity to the input: the single phosphorylation cycle, the double phosphorylation cycle and the phosphotransfer system, all with a kinase as input. In all three cases, the trade-off is due to the fact that, as the total substrate concentration is increased to attenuate the effect of retroactivity on the output, the system applies a large retroactivity to the input. Thus, the requirements (i) and (ii) of Def. 1 cannot be independently achieved. In [34], a cascade of phosphotransfer systems was found to apply a small retroactivity to the input and to attenuate retroactivity to the output. Further, cascades of single and double PD cycles are ubiquitous in cellular signaling, such as in the MAPK cascade [14], [47]. The two-component signaling system (Section 3.3) is also often the first stage of a cascade of signaling reactions [46], [15]. Motivated by this, here we consider a cascade of PD cycles to determine how a cascaded architecture can overcome this trade-off. We have found that single and double PD cycles, and the phosphotransfer system, show similar properties with respect to unidirectional signal transmission. Thus, our findings are applicable to all systems composed of cascades of single stage systems, such as the single PD cycle, the double PD cycle and the phosphotransfer system analyzed in Section 3.3 (simulation results for cascades of different systems are in SI 5.6 Fig. 12 and Fig. 13).

We consider a cascade of two single phosphorylation cycles, shown in Fig. 6A. The input signal is *Z*, the concentration of kinase Z. Z phosphorylates substrate X_1_ to 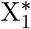, which acts as a kinase for substrate X_2_, phosphorylating it to 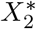. Both 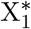 and 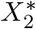 are dephosphorylated by a common phosphatase M. The output signal is 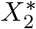, the concentration of 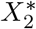.

**Figure 6:**
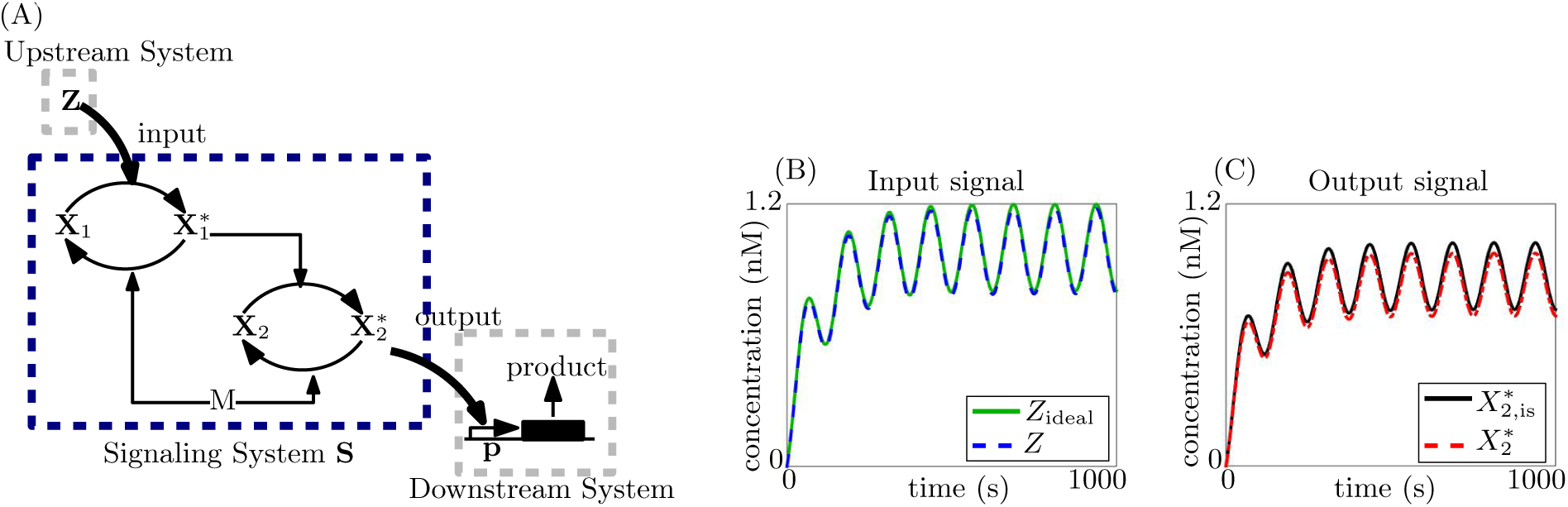
Tradeoff between small retroactivity to the input and attenuation of retroactivity to the output is overcome by a cascade of single phosphorylation cycles. (A) Cascade of 2 phosphorylation cycles that with kinase Z as the input: Z phosphorylates X_1_ to 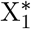, 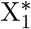 acts as the kinase for X_2_, phosphorylating it to 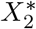, which is the output, acting on sites p in the downstream system, which is depicted as a gene expression system here. Both 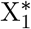 and 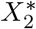 are phosphorylated by phosphatase M. (B), (C) Simulation results for ODEs (69)-(86) in SI Section 5.6 with N = 2. Simulation parameters are given in Table 1 in SI Section 5.2. Ideal system is simulated for *Z*_ideal_ with *X*_*T*1_ = *X*_*T*2_ = *M_T_* = *p_T_* = 0. Isolated system is simulated for 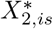 with *p_T_* = 0.

The input *Z* is produced at a time-varying rate *k*(*t*), and all species dilute with rate constant *δ*. The substrate of the cycles are produced at constant rates *kx*_1_ and *kx*_2_, respectively, and the phosphatase is produced at a constant rate *k_M_*. We then define 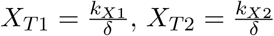 and 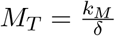. The concentration of promoter sites in the downstream system is *p_T_*. The Michaelis-Menten constants for the phosphorylation and dephosphorylation reactions are *K*_*m*1_ and *K_m2_*, respectively (assuming identical reaction-rate parameters for both cycles), and catalytic rate constants are *k*_1_ and *k*_2_. The chemical reactions for this system are shown in eqns. (62)-(68) in SI Section 5.6. As before, the parameters we vary to analyze this system's ability to transmit unidirectional signals are *X*_*T*1_, *X*_*T*2_ and *M*_T_. Using the procedure in Fig. 3, we seek to tune these to satisfy the requirements of Def. 1. We find what follows. (Steps 1-5 are detailed in SI Section 5.6).

(i) Retroactivity to the input: Evaluating the terms in Step 3, we _nd that to satisfy Test (i), 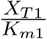 must be small. Thus, to have a small retroactivity to the input, *X*_*T*1_ must be small. (Evaluation of terms in Step 3 is shown in SI Section 5.6).
(ii) Retroactivity to the output: As before, we evaluate the terms in Step 4, and find that to satisfy Test (ii), we must have small 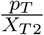 and 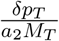. Thus, to attenuate retroactivity to the output, *M_T_* and *X*_*T*2_ must be large. (Evaluation of terms in Step 4 is shown in SI Section 5.9).
(iii) Input-output relationship: Evaluating *I*<u>Γ</u> as in Step 5, we find that the input-output relationship is 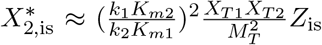 when *K*_*m*1_, *K*_*m*2_ ≫ *Z*_is_. (Details are shown in SI Section 5.6).

The ratio 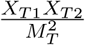 can thus be tuned such that the system satisfies Test (iii) with *m* = 1. However, as 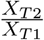 increases beyond a point, the second stage of the cascade affects the first stage, and the output begins to saturate with respect to the input, thus not satisfying Test (iii). In SI 5.6, we have shown that this non-linearity can be reduced by additional cycles, between the first and second cycle, in the cascade up to a certain number of cycles. That is, there exists an optimal number of cycles in the cascade for which the term leading to a non-linear input-output response (shown in eqn. (90) in SI Section 5.6) is minimized. This is because, each downstream cycle affects the response of the cycle directly upstream to it, making it non-linear. For each cycle, these non-linearities add up, and thus the number of terms contributing to the total non-linearity increase with the number of cycles. However, additional cycles reduce the non-linear effect of each individual stage. These two opposing effects make it so that the net non-linearity in the output of the final stage has an optimum.

Finally, we see that Test (iv) is satisfied for this system, since Tests (i)-(iii) can be satisfied simultaneously. We thus note that the trade-off between attenuating retroactivity to the output and imparting small retroactivity to the input, found in single-stage systems is broken by having a cascade of two cycles. This is because the input kinase Z only directly interacts with the first cycle, and thus when *X*_*T*1_ is made small, the upstream system faces a small reaction flux due to the phosphorylation reaction, making retroactivity to the input small. The downstream system sequesters the species 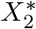, and when *X*_*T*2_ is made high, there is enough substrate X_2_ available for the signaling system to be nearly unaffected, thus attenuating retroactivity to the output. This is verified in Figs. 6B,6C. The trade-off found in the single cycle in Figs. 2B-2E is overcome by the cascade, where we have tuned *M_T_* to satisfy requirement (iii) of Def. 1. When the total substrate concentration for a single cycle is low, the retroactivity to the input is small (Fig. 2B) but the retroactivity to the output is not attenuated (Fig. 2C). When the total substrate concentration of this cycle is increased, the retroactivity to the output is attenuated (Fig. 2D) but the input, and therefore the output, are highly changed due to an increase in the retroactivity to the input (Figs. 2D, 2E). When the same two cycles are cascaded, with the low substrate concentration cycle being the first and the high substrate concentration cycle being the second (and *M_T_* tuned to maintain the same gain *K* as the single cycles), retroactivity to the input is small and retroactivity to the output is attenuated (Figs. 6B, 6C). Thus, cascading two cycles overcomes the trade-off found in a single cycle.

These results are summarized in Fig. 10E. While the system demonstrated here is a cascade of single phosphorylation cycles, the same decoupling is true for cascaded systems composed of double phosphorylation cycles and phosphorylation cycles followed by phosphotransfer, which as we saw in the previous sub Section 5.9s, show a similar kind of trade-off. Cascades of such systems, with the first system with a low substrate concentration and the last system with a high substrate concentration thus both, impart a small retroactivity to the input, and attenuate retroactivity to the output and are therefore able to transmit unidirectional signals. This can be seen via simulation results in SI Section 5.6, where a cascade of a phosphotransfer system and a single PD cycle is seen in Fig. 12 and a cascade of a single PD cycle and a double PD cycle is seen in Fig. 13.

### 3.5 Phosphotransfer with the phosphate donor undergoing autophosphorylation as input

Here, we consider a signaling system composed of a protein X_1_ that undergoes autophosphorylation and then transfers the phosphate group to a substrate X_2_, shown in Fig. 7A. In Section 3.3, we considered a system with regulated autophosphorylation, where the input is a ligand/kinase. In this Section 5.9, motivated by proteins that undergo autophosphorylation and then transfer the phosphate group, we consider a system where the input is the protein undergoing autophosphorylation (substrate input). Based on our literature review, we have not found instances of such systems in nature, and in this Section 5.9 we investigate whether they might pose a disadvantage to unidirectional signaling. The input signal *U* of Fig. 1A is *X*_1_, the concentration of protein X_1_ which undergoes autophosphorylation, and the output signal *Y* of Fig. 1A is 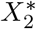, the concentration of phosphorylated protein 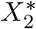. The total protein concentrations of substrate X_2_ and phosphatase M are *X*_*T*2_ and *M_T_*, respectively. The total concentration of promoters in the downstream system is *P_T_*. Autophosphorylation of a protein typically follows a conformational change that either allows the protein to dimerize and phosphorylate itself, or the conformational change stimulates the phosphorylation of the monomer [48]. Here, we model the latter mechanism for autophosphorylation as a single step with rate constant *π*_1_. The Michaelis-Menten constant for the dephosphorylation of 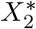 by M is *K*_*m*3_ and the association, dissociation and catalytic rate constants for this reaction are *a_3_, d_3_* and *k*_3_. The association and dissociation rate constants for the complex formed by 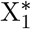 and X_2_ are *a*_1_ and *d*_1_, the dissociation rate constant of this complex into X_1_ and 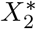 is *d*_2_, and the corresponding reverse association rate constant is *a*_2_. The input protein X_1_ is produced at a time-varying rate *k*(*t*). Details of the chemical reactions of this system are shown in SI Section 5.7 eqn. (98). We use the procedure in Fig. 3 to analyze this system as per Def. 1 by varying the total protein concentrations *X*_*T*2_ and *M_T_*. This is done as follows. (Steps 1-5 are detailed in SI Section 5.7).

**Figure 7:**
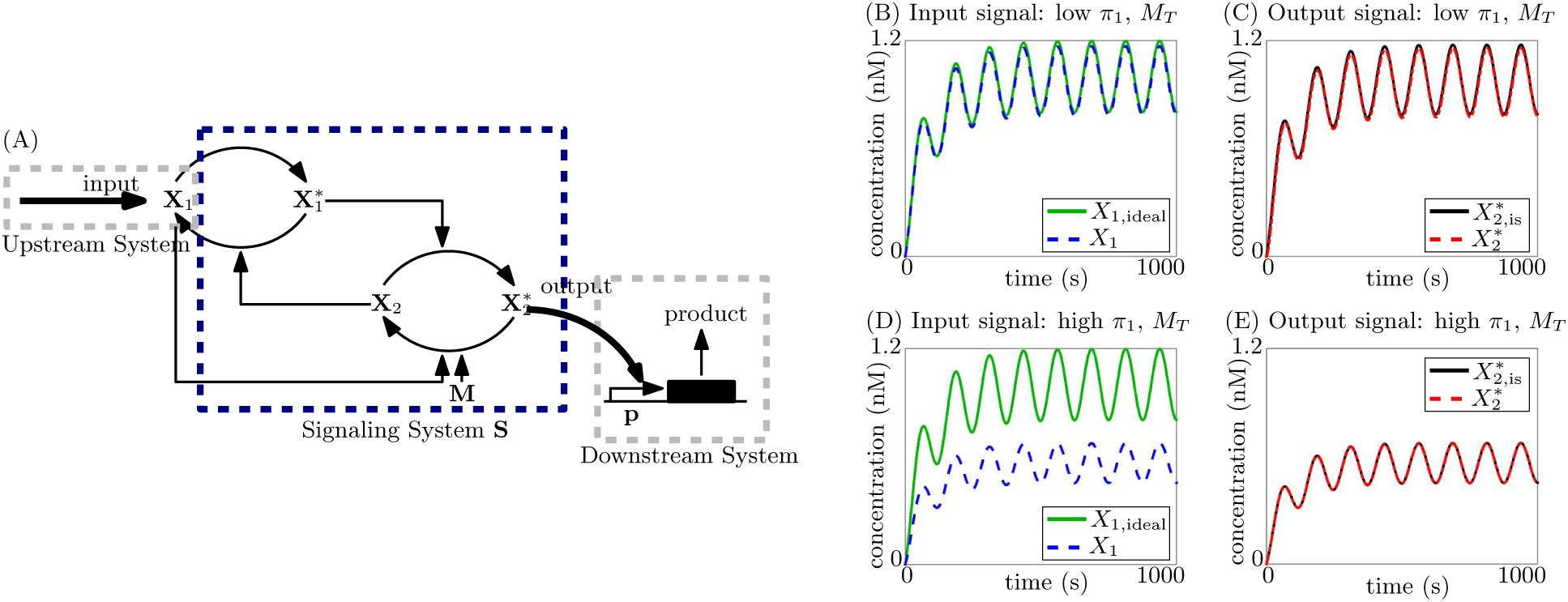
Attenuation of retroactivity to the output by a phosphotransfer system. (A) System with autophosphorylation followed by phosphotransfer, with input as protein X_1_ which autophosphorylates to 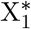 The phosphate group is transferred from 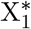 to X_2_ by a phosphotransfer reaction, forming 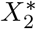, which is in turn dephosphorylated by the phosphatase M. 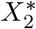 is the output and acts on sites p in the downstream system, which is depicted as a gene expression system here. (B)-(E) Simulation results for ODE (99) in SI Section 5.7. Simulation parameters are given in Table 1 in SI Section 5.9. Ideal system is simulated for *X*_1,ideal_ with *X*_*T*2_ = *M_T_* = *π*_1_ = *p_T_* = 0. Isolated system is simulated for 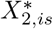 with *p_T_* = 0.

(i) Retroactivity to input: Evaluating the terms in Step 3, we find that to satisfy Test (i), 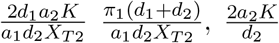 and 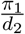 must be small, where 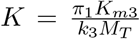 However, not all these terms can be made smaller by varying *X*_*T*2_ and *M_T_* alone. Thus, the retroactivity to the input, and whether or not Test (i) is satisfied, depends on the reaction rate constants of the system, and it is not possible to tune it using total protein concentrations alone. (Evaluation of terms in Step 3 is shown in SI Section 5.7).
(ii) Retroactivity to output: Evaluating the terms in Step 4, we find that to satisfy Test (ii), we must have a small 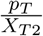 and 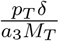. Thus, to attenuate retroactivity to the output, *X*_*T*2_ and *M_T_* must be large. (Evaluation of terms in Step 4 is shown in SI Section 5.7).
(iii) Input-output relationship: Evaluating *I*<u>Γ</u> as in Step 5, we find that the input-output relationship is 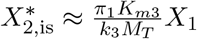 when 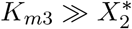 and thus, this system can satisfy Test (iii) by tuning *M_T_* to achieve a desired *K* with *m* = 1. (Details of Step 5 are shown in SI Section 5.7).

Thus, we find that the retroactivity to the input cannot be made small by changing concentrations alone. The retroactivity to the output can be attenuated by having a large *X*_*T*2_ and *M_T_*, since these can compensate for the sequestration of 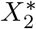 by the downstream system. This signaling system can therefore satisfy Tests (ii) and (iii) for unidirectional signal transmission. While satisfying these requirements does not increase the retroactivity to the input, thus making it possible for it to satisfy Test (i) as well, retroactivity to the input depends on the reaction-rate parameters, in particular, on the forward reaction rate constant *π*_1_ of autophosphorylation of X_1_. If this is large, the autophosphorylation reaction applies a large reaction flux to the upstream system, thus resulting in a large retroactivity to the input. If *π*_1_ is small, this flux is small, and thus retroactivity to the input is small. By the way we have defined cascades (as signals between stages transmitted through a kinase), any cascade containing this system would have it as a first stage. Therefore, even cascading this system with different architectures would not overcome the above limitation. These mathematical predictions can be appreciated in the simulation results shown in Figs. 7B-7E. The result is summarized in Fig. 10C.

### 3.6 Single cycle with substrate input

Here, we consider a single phosphorylation cycle where the input signal *U* of Fig. 1A is *X*, the concentration of the substrate X, and the output signal *Y* is X^*^, the concentration of the phosphorylated substrate. We consider this system motivated by the various transcription factors that undergo phosphorylation before activating or repressing their targets, such as the transcriptional activator NRI in the *E. Coli* nitrogen assimilation system [49]. However, to the best of our knowledge, based on our literature review, signals are more commonly transmitted through kinases, as opposed to being transmitted by the substrates of phosphorylations. Since these are less represented than the others in natural systems, we ask whether they have any disadvantage for unidirectional transmission, and in fact they do. Note that the system analyzed in Section 3.5 is a system that takes as input a kinase that undergoes autophosphorylation before donating the phosphate group, and is not the same as the system considered here, where the input is a substrate of enzymatic phosphorylation.

The signaling system we consider, along with the upstream and downstream systems, is shown in Fig. 8A. The input protein X is produced at a time-varying rate *k*(*t*). It is phosphorylated by kinase Z to the output protein X^*^, which is in turn dephosphorylated by phosphatase M. X^*^ then acts as a transcription factor for the promoter sites in the downstream system. All the species in the system decay with rate constant *δ*. The total concentration of promoters in the downstream system is *p_T_*. The total kinase and phosphatase concentrations are *Z*_T_ and *M_T_*, respectively, which are the parameters of the system we vary. The Michaelis-Menten constants of the phosphorylation and dephosphorylation reactions are *K*_*m*1_ and *K*_*m*2_, and the catalytic rate constants are *k*_1_ and *k*_2_. The chemical reactions of this system are shown in eqn. (109) in SI Section 5.8. Using the procedure in Fig. 3, we analyze if this system can transmit a unidirectional signal according to Definition 1 by varying *Z*_T_ and *M*_T_. This is done as follows. (Steps 1-5 in SI Section 5.8).

**Figure 8:**
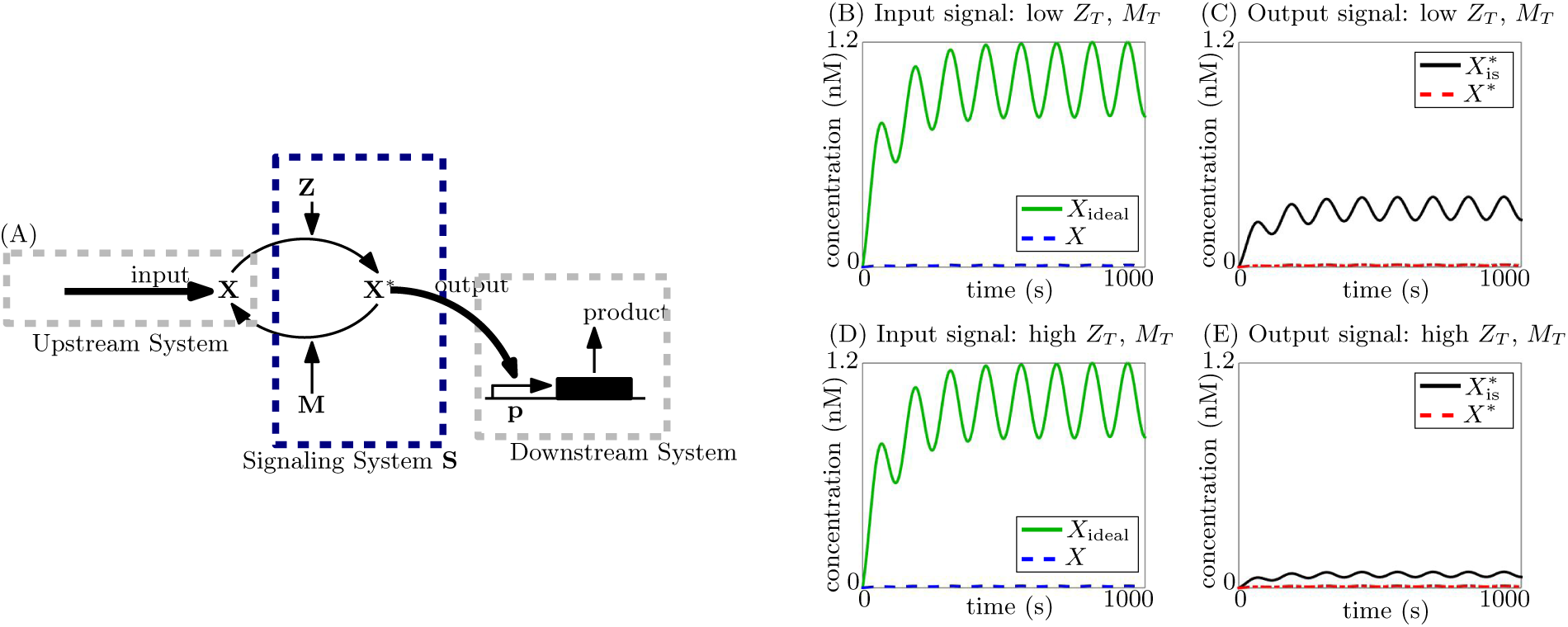
Inability to attenuate retroactivity to the output or impart small retroactivity to the input by single phosphorylation cycle with substrate as input. (A) Single phosphorylation cycle, with input X as the substrate: X is phosphorylated by the kinase Z to X^*^, which is dephosphorylated by the phosphatase M back to X. X^*^ is the output and acts as a transcription factor for the promoter sites p in the downstream system. (B)-(E) Simulation results for ODEs (110), (111) in SI Section 5.8. Simulation parameters are given in Table 1 in SI Section 5.2. Ideal system is simulated for *X*_ideal_ with *Z*_T_ = *M_T_* = *p_T_* = 0. Isolated system is simulated for 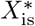 with *p_T_* = 0.

(i) Retroactivity to the input: Evaluating the terms in Step 3, we find that they cannot be made small by changing *Z*_T_ and *M_T_*, and therefore, Test (i) fails and retroactivity to the input cannot be made small. (Evaluation of terms in Step 3 is shown in SI Section 5.8).
(ii) Retroactivity to the output: Similarly, we evaluate the terms in Step 4 and find that they cannot be made small by varying *Z*_T_ and *M_T_*. Thus, Test (ii) fails and retroactivity to the output cannot be attenuated by tuning these parameters. (Evaluation of terms in Step 4 is shown in SI Section 5.8).
(iii) Input-output relationship: Evaluating *I*<u>Γ</u> as in Step 5, we find that the input-output relationship is linear with gain 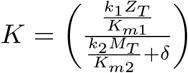 when *K*_*m*1_, *K*_*m*2_ ≫ *X*, that is:

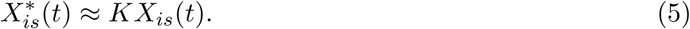

The input-output relationship is thus linear, i.e., *m* =1, and *K* can be tuned by varying *Z_T_* and *M_T_*. The system thus satisfies Test (iii). (Details of Step 5 are shown in SI Section 5.8).

Thus, we find that a signaling system composed of a single phosphorylation cycle with substrate as input cannot transmit a unidirectional signal, since it can neither make retroactivity to the input small nor attenuate retroactivity to the output. This is because, the same protein X is the input (when unmodified) and the output (when phosphorylated). Thus, when X undergoes phosphorylation, the concentration of input X is reduced by conversion to X^*^, thus applying a large retroactivity to the input. Now, when X^*^ is sequestered by the downstream system, this results in a large flux to both X and X^*^, and thus the retroactivity to the output is also large. Cascading such a system would also not enhance its ability to transmit unidirectional signals: if the system were used as the first stage to a cascade, it would apply a large retroactivity to the input for the aforementioned reasons. The way we have defined cascades above, with non-initial stages receiving their input via a kinase, this system cannot be the second stage of a cascade since it takes its input in the form of the substrate. These results are demonstrated in the simulation results shown in Fig. 8B-8E and summarized in Fig. 10F.

### 3.7 Double cycle with substrate input

Finally, we consider a double phosphorylation cycle with input signal *U* of Fig. 1A as the concentration of the substrate, *X*, and the output signal *Y* as the concentration of the doubly phosphorylated substrate, *X^**^*. Similar to the single phosphorylation cycle, we consider this system to model cases where the input species undergoes double phosphorylation before acting on its downstream targets, such as transcription factor FKHRL1, which is phosphorylated by Akt at its T23 and S253 sites [50]. In this system, the signal is transmitted by the kinase Akt and not the substrate. Based on our literature review, we have not found systems where the signal is transmitted by the substrate in such an architecture. We therefore consider this architecture to test whether it has a disadvantage for unidirectional signal transmission. The arrangement is shown in Fig. 9A. All species dilute with rate constant *δ*. The total concentration of promoters in the downstream system is *p_T_*. The total concentration of kinase Z and total concentration of phosphatase M are *Z_T_* and *M_T_*, respectively. The input X is produced at a time-varying rate *k*(*t*). Using the procedure described in Fig. 3, we vary *Z_T_* and *M_T_* to investigate if this system can transmit unidirectional signals according to Def. 1. (Steps 1-5 are detailed in SI Section 5.9). This is done as follows:

(i) Retroactivity to the input: Evaluating the terms in Step 3, we find that they cannot be made small by tuning *Z_T_* and *M_T_*, and thus, Test (i) is not satisfied. (Evaluation of terms in Step 3 is shown in SI Section 5.9).
(ii) Retroactivity to the output: Evaluating the terms in Step 4, we find that these cannot be made small by tuning *Z_T_* and *M_T_*. Thus, Test (ii) is not satisfied. (Evaluation of terms in Step 4 is shown in SI Section 5.9).
(iii) Input-output relationship: Evaluating *Ι*<u>Γ</u> as in Step 5, we find that 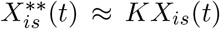 for *t ∈* [*t_b_*, *t_f_*] for large Michaelis-Menten constants, where *K* can be tuned by tuning the total kinase and phosphatase concentrations *Z_T_* and *M_T_*. Thus, the system satisfies Test (iii) with *m* = 1 and a desired *K*. (Details of Step 5 are shown in SI Section 5.9).

**Figure 9:**
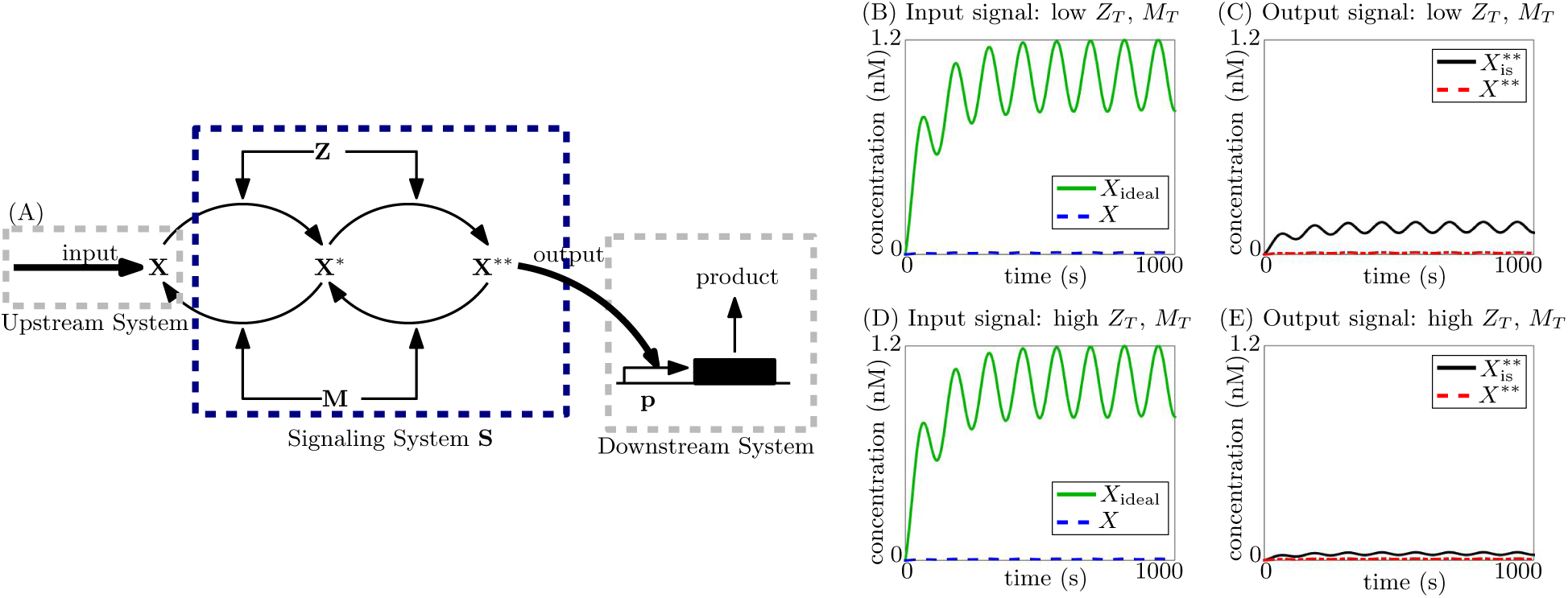
Inability to attenuate retroactivity to the output or impart small retroactivity to the input by double phosphorylation cycle with substrate as input. (A) Double phosphorylation cycle, with input X as the substrate: X is phosphorylated twice by the kinase K to X^*^ and X^**^, which are in turn dephosphorylated by the phosphatase M. X^**^ is the output and acts on sites p in the downstream system, which is depicted as a gene expression system here. (B)-(E) Simulation results for ODE (124) in SI Section 5.9. Simulation parameters are given in Table 1 in SI Section 5.2. Ideal system is simulated for *X*_ideal_ with *Z_T_* = *M_T_* = *p_T_* = 0. Isolated system is simulated for 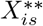 with *p_T_* = 0.

Thus, similar to the single cycle with substrate as input, the double cycle with substrate as input provides a linear input-output relationship but is not able to impart a small retroactivity to the input, nor is it able to attenuate retroactivity to the output, even upon cascading with other systems. These properties are shown in Fig. 9B-9E, and the results are summarized in Fig. 10G.

**Figure 10:**
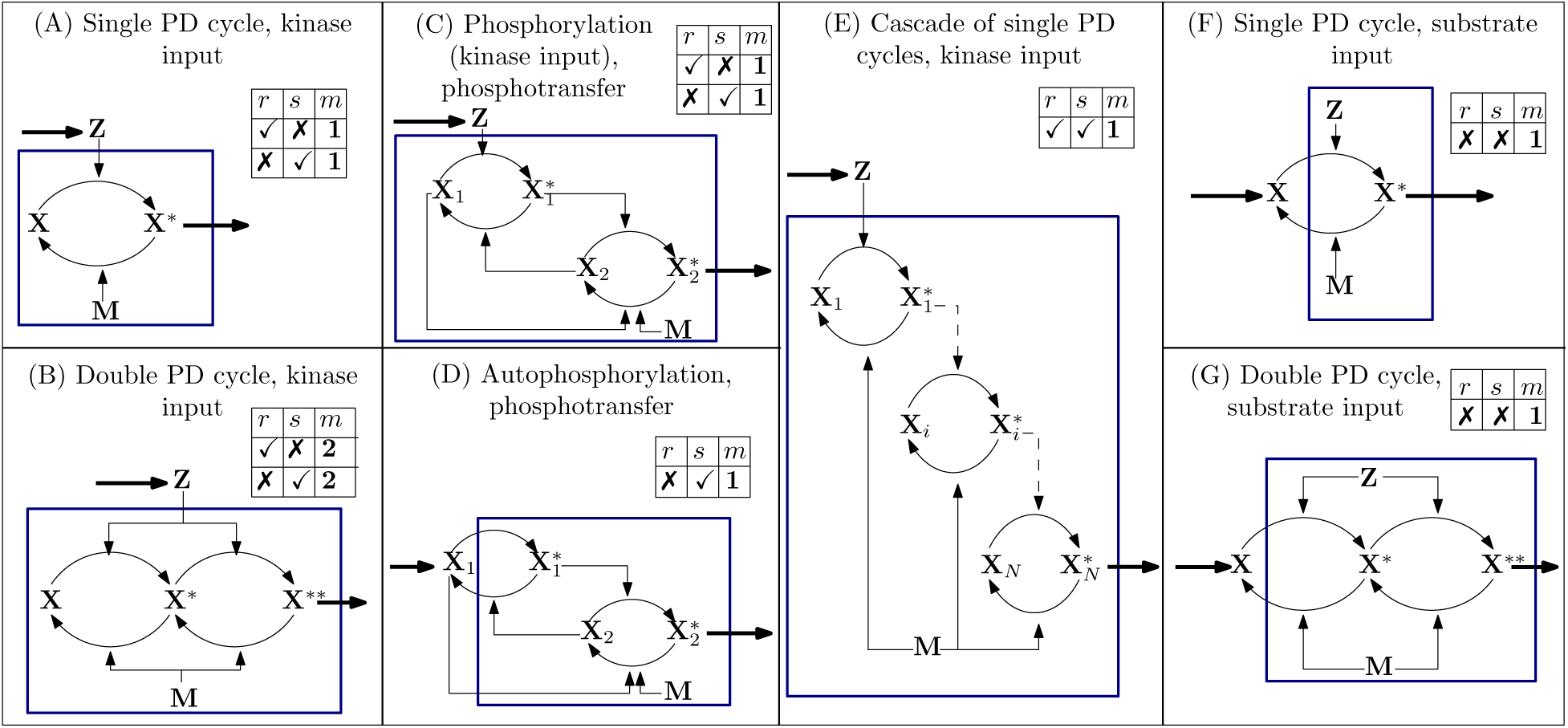
Table summarizing the results. For each inset table, a ✓ (✗) for column *r* implies the system can (cannot) be designed to minimize retroactivity to the input by varying total protein concentrations, a ✓ (✗) for column *s* implies the system can (cannot) be designed to attenuate retroactivity to the output by varying total protein concentrations, column *m* describes the input-output relationship of the system with *m* as described in Def. 1(iii). Inset tables with two rows imply that one of the two rows can be achieved for a set of values for the design parameters: thus, the two rows for systems (A), (B) and (C) show the trade-off between the ability to minimize retroactivity to the input (first row) and the ability to attenuate retroactivity to the output (second row). Note that this trade-off is overcome by the cascade (E).

## 4 Discussions

The goal of this work was to identify signaling architectures that can overcome retroactivity and thus allow the transmission of unidirectional signals. To achieve this, we have provided a procedure that can be used to analyze any signaling system composed of reactions such as phosphorylation-dephosphorylation and phosphotransfer. We have then considered different signaling architectures (Fig. 10), and have used this procedure to determine whether they have the ability to minimize retroactivity to the input and attenuate retroactivity to the output.

We have found that a main discriminating factor is whether the signaling architecture transmits information from kinases or from substrates. Specifically, phosphorylation cycles (single or double) and phosphotransfer systems that transmit information from an input kinase (Figs. 10A, 10B, 10C) show a trade-off between minimizing retroactivity to the input and attenuating retroactivity to the output, consistent with prior experimental studies [33], [51]. Yet, cascades of such systems (see, for example Fig. 10E) can break this trade-off. This is achieved when the first stage has low substrate concentration, thus imparting a small retroactivity to the input, and the last stage has high substrate concentration, thus attenuating retroactivity to the output. Interestingly, this low-high substrate concentration pattern appears in the MAPK signaling cascade in the mature Xenopus Oocyte, where the first stage is a phosphorylation cycle with substrate concentration in the nM range and the last two stages are double phosphorylation cycle with substrate concentration in the thousands of nM [25]. This low-high pattern indicates an ability to overcome retroactivity and transmit unidirectional signals, and while this structure may serve other purposes as well, it is possible that the substrate concentration pattern has evolved to more efficiently transmit unidirectional signals. By contrast, architectures that transmit information from a substrate (Figs. 10D, 10F, 10G) do not perform as well even when cascaded. Consistent with this finding, while architectures that transmit signals from an input kinase are highly represented in cellular signaling, such as in the MAPK cascade and two-component signaling [1]–[7], [16]–[19], those receiving signals through substrates are not as frequent in natural systems. It has also been reported that kinase-to-kinase relationships are highly conserved evolutionarily [52], implying that upon evolution, signaling mechanisms where kinases phosphorylate other kinases are conserved. These facts support the notion that cellular signaling has evolved to favor one-way transmission.

For graph-based methods for analyzing cellular networks [53], such as discovering functional modules based on motif-search or clustering, signaling pathway architectures that transmit unidirectional signals can then be treated as directed edges. On the contrary, analysis of signaling systems (such as those with a substrate as input) that do not demonstrate the ability to transmit unidirectional signals must take into account effects of retroactivity. These effects could result in crosstalk between different targets of the signaling system, since a change in one target would affect the others by changing the signal being transmitted through the pathway [13]. Our work provides a way to identify signaling architecture that overcome such effects and that can be treated as modules whose input/output behavior is largely independent from the context. Our findings further uncover a library of systems that transmit unidirectional signals, which could be used in synthetic biology to connect genetic components, enabling modular circuit design.

## 5 Supplementary Information

### 5.1 Assumptions and Theorems

For the general system (1), we make the following Assumptions:

#### Assumption 1

Phosphorylation-dephosphorylation and phosphotransfer reactions typically occur at rates of the order of second^-1^ [54], [55], much faster than transcription, translation and decay, which typically occur at rates of the order of hour^-1^ [56]. Then, *G*_1_ ≫ 1.

#### Assumption 2

Binding-unbinding reactions of the output with the promoter sites in the downstream system are much faster than transcription, translation and decay [57]. Then, *G*_2_ ≫ 1.

#### Assumption 3

The eigenvalues of 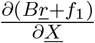 and 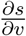 have strictly negative real parts.

#### Assumption 4

There exist invertible matrices *T* and *Q*, and matrices *M* and *P*, such that *TA* + *MB* = 0, *M f*_1_ = 0 and *QC* + *PD* = 0.

#### Assumption 5

Let *<u>X</u>* = <u>Ψ</u>(*U*, *υ*) be the locally unique solution to *f*_1_(*U*, *<u>X</u>*, *S*_3_*υ*) + *B<u>r</u>*(*U*, *<u>X</u>*, *S*_2_*υ*) = 0. We assume <u>Ψ</u>(*U*, *υ*) is Lipschitz continuous in *υ* with Lipschitz constant *L*_Ψ._

#### Assumption 6

Let *υ* = *ϕ*(*<u>Χ</u>*) be the locally unique solution to *s*(*<u>X</u>*, *υ*) = 0. Define the function *f*(*U*, *<u>X</u>*) = *<u>X</u>* − <u>Ψ</u>(*U*, *ϕ*(*<u>Χ</u>*)). Then the matrix 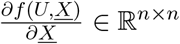 is invertible.

#### Assumption 7

Let <u>Γ</u>(*U*) be the locally unique solution to *B<u>r</u>*(*U*, *<u>X</u>*, *S*_2_*υ*) + *f*_1_(*U*, <u>X</u>, *S*_3_*υ*) = 0. We assume that <u>Γ</u>(*U*) is Lipschitz continuous with Lipschitz constant *L*_Γ_.

#### Remark 1

By definition of <u>Γ</u>(*U*), we have that <u>Γ</u>(*U*) = <u>Ψ</u>(*U, ϕ* (<u>Γ</u>*(U*))), since *υ* = *ϕ(<u>Χ</u>)* satisfies *s*(*<u>X</u>*, *υ*) = 0 and *<u>X</u>* = <u>Ψ</u>(*U*, *<u>X</u>*) satisfies *f*_1_(*U, <u>X</u>, S_3_*υ*)* + *B<u>r</u>*(*U*, *<u>X</u>*, *S*_2_*υ*) = 0. If *S*_2_ = *S*_3_ = 0, <u>Γ</u>(*U*) is independent of *υ*, which is denoted by <u>Γ</u>_*is*_(*U*). Then, <u>Γ</u>_is_(*U*) = <u>Ψ</u>(*U*, 0) since *S*_2_ = *S*_3_ = 0. Thus, the difference |<u>Γ</u>_*is*_(*U*) − <u>Γ</u>(*U*)| depends on *S*_2_ and *S*_3_, and is zero when *S*_2_ = *S*_3_ = 0. We thus sometimes denote <u>Γ</u>(*U*) as <u>Ψ</u>(*U*, *g*(*S*_2_, *S*_3_)*ϕ*(<u>Γ</u>(*U*))), where *g*(*S*_2_, *S*_3_) = 0 if both *S*_2_ = *S*_3_ = 0. Further, since as ||*S*_2_|| and ||*S*_3_|| decrease, the dependence of f_1_(*U*, *<u>X</u>*, *S*_3_*υ*) + *B<u>r</u>*(*U*, *<u>X</u>*, *S*_2_*υ*) on *υ* decreases, by the implicit function theorem, *g*(*S_2_, S_3_*) decreases as ||*S*_2_|| and ||*S*_3_|| decrease.

#### Remark 2

On picking *S*_2_ and *S*_3_ for the systems: *S*_2_ and *S*_3_ are picked such that they appear in the form of *Y* + *S*_2_*υ* and *Y* + *S*_3_*υ* in the ODEs when written in form (1).

#### Assumption 8

The function *f*_0_(*U*, *t*) is Lipschitz continuous in *U* with Lipschitz constant *L*_0_. The function *<u>r</u>*(*U*, *<u>X</u>*, *υ*) is Lipschitz continuous in *<u>X</u>* and *υ*.

#### Assumption 9

The system:

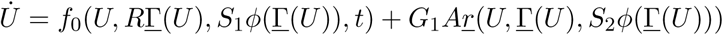

is contracting [58] with parameter λ.

We now state the following result from [51]:

#### Lemma 1

*If the following system:*

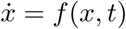

*is contracting with contraction rate λ, then, for the perturbed system:*

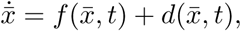

*where there exists a d̅* ≥ 0 *such that* |*d*(*x̅*, *t*) | ≥ *d̅ for all x̅, t, the difference in trajectories for the actual and perturbed system is given by:*

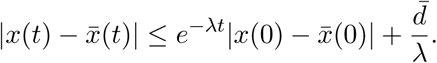

We state the following result, adapted from [32], for system (1):

#### Lemma 2

*Under Assumptions 1-4*, 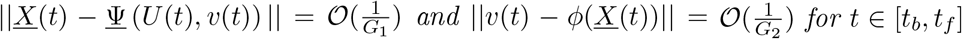, *where* <u>Ψ</u>(*U*, *υ*) *is defined in Assumption 5, ϕ*(*<u>X</u>*) *is defined in Assumption 6 and *t_b_* is such that t_i_*< *t_b_* < *t_f_ and t_b_* − *t_i_ decreases as G*_1_ *and G*_2_ *increase.*

*Proof of Lemma 2.* We bring the system to standard singular perturbation form, by defining *<u>w</u> =* Q*<u>X</u>* + *P*υ** and *z* = *TU* + *M*(*<u>X</u> + Q^-1^P*υ**). Under Assumption 4, we obtain the following system:

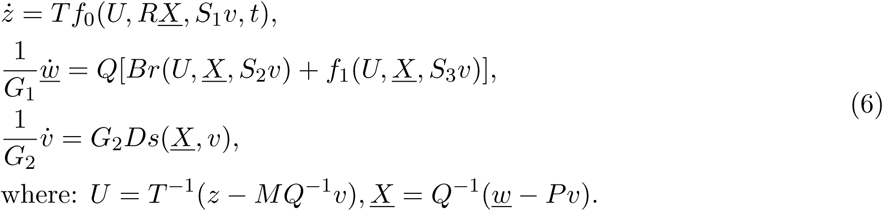

Under Assumptions 1-3, this system is in the standard singular perturbation form with 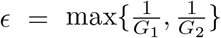. We define function *<u>W</u>*(*z*, *υ*), such that <u>w</u> = *<u>W</u>* is a solution to *(Br* + *f*_1_)(*z*, *<u>w</u>*, *υ*) = 0 and function *V*(*<u>w</u>*) such that *υ* = V is a solution to *s*(*<u>w</u>*, *υ*) = 0. Applying singular perturbation, we then have 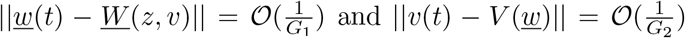. Rewriting these expressions in terms of the original variables, we use the definitions in Assumptions 5 and 6, we have 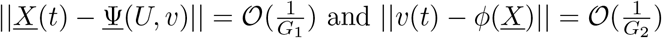.

#### Lemma 3

*Under Assumptions 1-6,* ||<u>X</u>(*t*) − <u>Γ</u>(*U*(*t*))|| = 𝒪 (ϵ), *for t* ϵ [*t_b_*, *t_f_*], *where* <u>Γ</u>(*U*) *is defined in Remark 1.*

*Proof of Lemma 3.* From Lemma 2, we have:

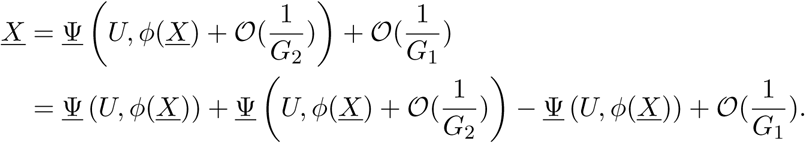

Under Assumption 5, using the Lipschitz continuity of <u>Ψ</u>(*U*, *υ*) we have:

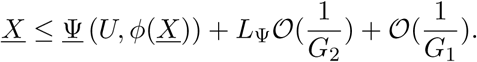

By definition of 𝒪, we have:

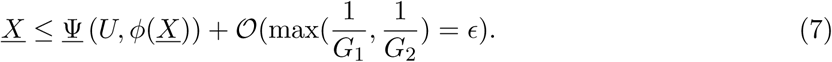

By equation (7), *f* (*U*, *<u>U</u>*) ≤ 𝒪 ϵ, where the function *f* is defined in Assumption 4. By definition of <u>Γ</u>(*U*), we have *f*(*U*, <u>Γ</u>(*U*)) = <u>Γ</u>(*U*) − <u>Ψ</u>(*U*, *ϕ*(<u>Γ</u>(*U*))) = 0. Therefore:

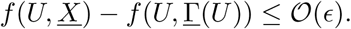

Under Assumption 5, *f* (*U*, *<u>X</u>*) is differentiable. Applying the Mean Value theorem [59], we have:

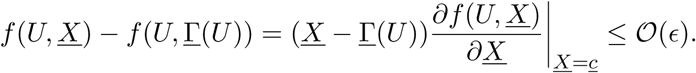

Under Assumption 6, the matrix 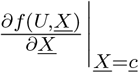 is invertible. Thus,

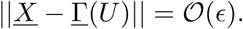

#### Lemma 4

*Under Assumptions 1-6, 8-9, for t ∈* [ *t_b_, t_f_*], *| U* (*t*) *− U̅* (*t*)*| = 𝒪*(*∈*) *where U̅ is such that:*

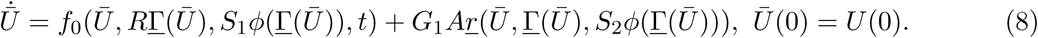

*Proof of Lemma 4.*

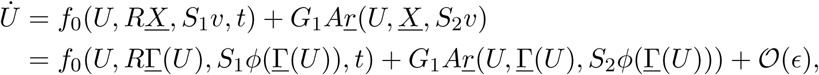

by Lemmas 2 and 3, since the functions *f_0_* and *<u>r</u>* are Lipschitz continuous under Assumption 8. Applying Lemma 1 to this system under Assumption 9, we have |*U*(*t*) − *U̅*(*t*) | = 𝒪(ϵ).

The first Theorem gives an upperbound on the retroactivity to the input. The terms in Step 3 and the corresponding Test (i) in the procedure in Fig. 3 arise from this result.

#### Theorem 1

*The effect of retroactivity to the input is given by:*

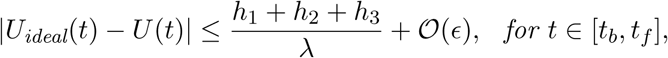

*Proof of Theorem 1.* By definition of *U*_ideal_, we have from (1):

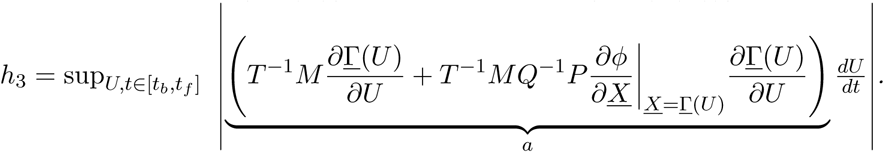

We define *U̅* such that its dynamics are given by (8), that is:

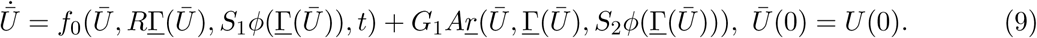

By the Lipschitz continuity of *f*_0_ under Assumption 8, we have:

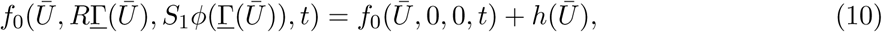

where |*h*(*U̅*)| ≤ *L*_o_|*R*<u>Γ</u>(*U̅*)| + *L_o_*|*S*_1_*ϕ*(<u>Γ</u>(*U̅*))| Thus, |*h*(*U̅*)| ≤ *h*_1_ + *h*_2_.

Further define *z* = *TU* + *M<u>X</u>* + *MQ*^-1^*P*υ*.* Then,

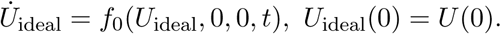

from eqns. (1). Using the expression of ˙ from (1), we then see that

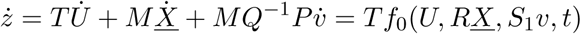

By Lemma 2 we have 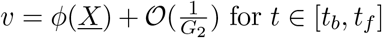 By Lemma 3 we have <u>X</u> =<u>Γ</u>(*U*)+𝒪 (ϵ) for *t ∈* [*t_b_*, *t_f_*]. *Thus,*

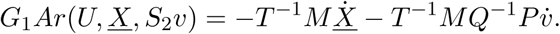

This implies that

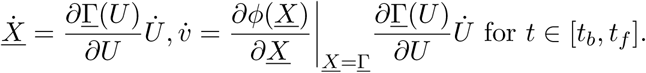

Then, under Assumption 8, due to the Lipschitz continuity of *<u>r</u>* and Lemmas 2 and 3,

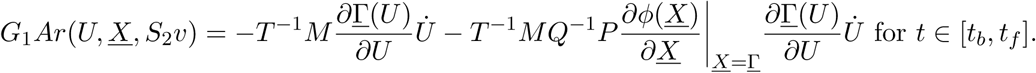

for *t ∈* [*t_b_, t_f_*]. Changing variables does not change the result, i.e., we define *q*(*U̅*) such that 
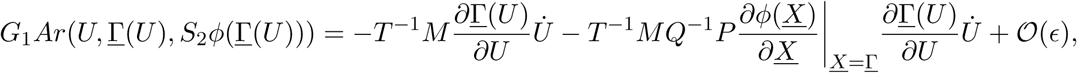

From the definition of *h*_3_ in Theorem 1, we have that |q(*U̅*)| ≤ *h*_3_ + 𝒪( ϵ). Thus, the dynamics of *U̅* as given by eqn. (9) can be rewritten using eqn. (10) and q(*U̅*) = G_1_A*r*(*<u>U̅</u>*, <u>Γ</u>(*U̅*), *S*_2_*ϕ*(<u>Γ</u>( *U̅*))) as:

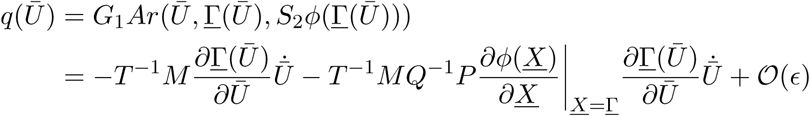

Using Lemma 1 we have that

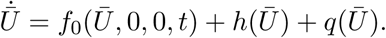

for *t ∈* [*t_b_*, *t_f_*]. From the triangle inequality, we know that |*U*_ideal_(*t*) − *U*(*t*)| ≤ |*U*_idea1_(*t*) − *U̅* (*t*)| + |*U̅*(*t*) − *U*(*t*)|. Using Theorem 4, we have:

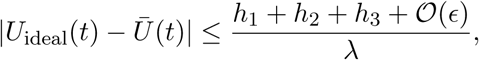

The next Theorem gives an upperbound on the retroactivity to the output. The terms in Step 4 and the corresponding Test (ii) in the procedure in Fig. 3 arise from this result.

#### Theorem 2

*The effect of retroactivity to the output is given by:*

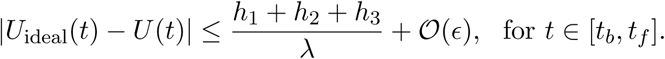

*Proof of Theorem 2.* By definition, *Y* (*t*) = *<u>IX</u>*(*t*). Under Lemma 3, this implies that *Y* (*t*) = *I*<u>Γ</u>(*U*(*t*)) + 𝒪( ϵ). The isolated output is then *Y*_is_(*t*) = *I*<u>Γ</u>_is_(*U*_is_(*t*)) + 𝒪( ϵ). Thus,

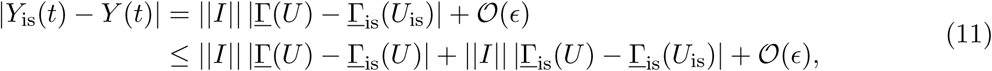

by the triangle inequality. By definition, as seen in Remark 1, <u>Γ</u>(*U*) = <u>Ψ</u>(*U*, g(*S*_2_, *S*_3_)*ϕ*(<u>Γ</u>(*U*))), where g(*S*_2_, *S*_3_) = 0 for *S*_2_ = *S*_3_ = 0. Also seen in Remark 1, <u>Γ</u>_is_(*U*) = <u>Ψ</u>(*U*, 0). Then, under Assumption 5,

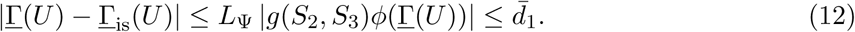

Under Assumption 7,

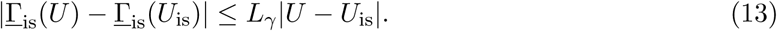

We now define *z* = *TU* + *M<u>X</u>* + *MQ^-1^P*υ*.* Then, from eqn. (1),

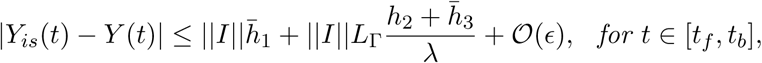

Then,

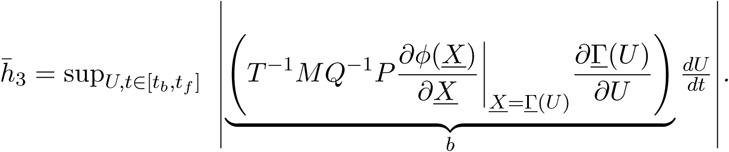

Comparing the equation above to eqns. (1) we have

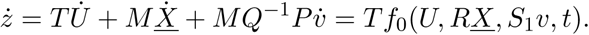

Thus we have that

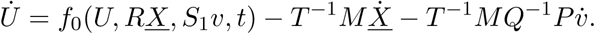

Thus, defining *U̅* as in eqn. (8), we have:
>
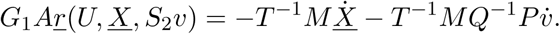

By the Lipschitz continuity of *f*_0_ under Assumption 8, this can be written as:

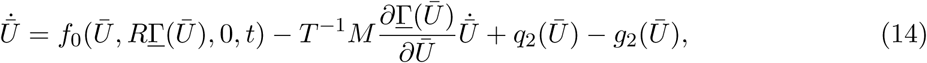

where |q_2_(*U̅*)| ≤ *L*_0_|*S*_1_ *ϕ*(<u>Γ</u>( *U̅*))| for all *U̅*. Thus, from the definition of *h_2_* in Theorem 2, we have that |q_2_(*U̅*)| ≤ *h*_2_. Further, we have

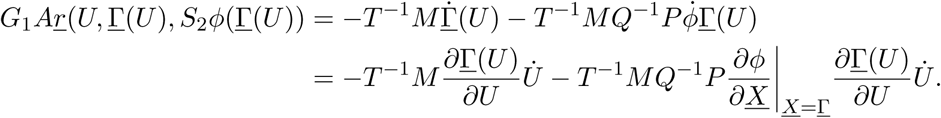

Since *˙* = *f*_0_(*U*, *R<u>X</u>*, *S*_1_*υ*, *t*) − *T*^−1^M˙ *− T^−1^MQ|^−1^P*υ**̇, the isolated input dynamics are by definition: ˙_*is*_ = *f*_0_(*U*, *R<u>X</u>*, 0, *t*) − *T*^-1^ *M*X. By Lemma 3 and under Assumption 8, this can be written as:

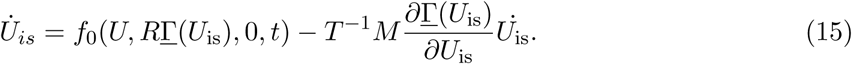

Applying Lemma 1 to systems (14) and (15) under Assumption 9, we have: 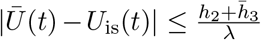 By the triangle inequality and Lemma 4,

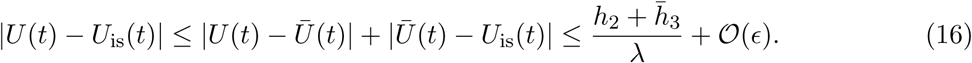

Using (11), (12), (13) and (16), we obtain the desired result.

The final Theorem gives an approximation of the input-output relationship. Step 5 and the corresponding Test (iii) in the procedure in Fig. 3 arise from this result.

#### Theorem 3

*The relationship between Y_is_*(*t*) *and U_is_*(*t*) *is given by:*

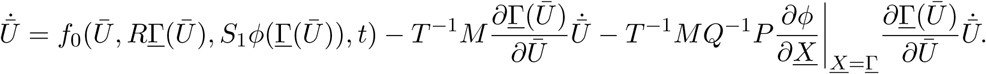

*Proof of Theorem 3.* From Remark 1, we see that <u>Γ</u>_*is*_(*U_is_*) = <u>Ψ</u>(*U_is_*, 0). From Lemma 2, we have ||*<u>X</u>*_is_(*t*) − <u>Ψ</u>(*U*_is_, 0)|| = 𝒪(ϵ). Thus, for *y_is_* = *I<u>X</u>*_is_, we have

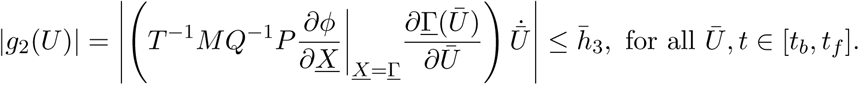

### 5.2 Table of simulation parameters

**Table 1.** Table of simulation parameters for Figures 2, 4-9.

| System | Simulation parameters |
| --- | --- |
| Single phosphorylation cycle with kinase input (Fig. 2) | $k(t) = 0.01(1 + \sin(0.05t))$, $\delta = 0.01s^{-1}$, $k_1 = k_2 = 600s^{-1}$, $a_1 = a_2 = 18nM^{-1}s^{-1}$, $d_1 = d_2 = 2400s^{-1}$, $k_{on} = 10nM^{-1}s^{-1}$, $k_{off} = 10s^{-1}$. Ideal system ($Z_{ideal}$): $X_T = M_T = p_T = 0$. Isolated system ($X_{is}^*$) with low substrate concentration: $X_T = M_T = 10nM$, $p_T = 0$. Actual system ($X^*, Z$) with low substrate concentration: $X_T = M_T = 10nM$, $p_T = 100nM$. Isolated system ($X_{is}^*$) with high substrate concentration: $X_T = M_T = 1000nM$, $p_T = 0$. Actual system ($X^*, Z$) with high substrate concentration: $X_T = M_T = 1000nM$, $p_T = 100nM$. |
| Double phosphorylation cycle with kinase input (Fig. 4) | $k(t) = 0.1(1 + \sin(0.05t))$, $\delta = 0.01s^{-1}$, $k_1 = k_2 = k_3 = k_4 = 600s^{-1}$, $a_1 = a_2 = a_3 = a_4 = 18nM^{-1}s^{-1}$, $d_1 = d_2 = d_3 = d_4 = 2400s^{-1}$, $k_{on} = 10nM^{-1}s^{-1}$, $k_{off} = 10s^{-1}$. Ideal system ($Z_{ideal}$): $X_T = M_T = p_T = 0$. Isolated system ($X_{is}^{**}$) with low substrate concentration: $X_T = 10nM$, $M_T = 3nM$, $p_T = 0$. Actual system ($X^{**}, Z$) with low substrate concentration: $X_T = 10nM$, $M_T = 3nM$, $p_T = 100nM$. Isolated system ($X_{is}^*$) with high substrate concentration: $X_T = 1200nM$, $M_T = 39nM$, $p_T = 0$. Actual system ($X^{**}, Z$) with high substrate concentration: $X_T = 1200nM$, $M_T = 39nM$, $p_T = 100nM$. |
| Phosphotransfer with phospho-donor phosphorylated by kinase input (Fig. 5) | $k(t) = 0.01(1 + \sin(0.05t))$, $\delta = 0.01s^{-1}$, $k_1 = k_2 = k_4 = 15s^{-1}$, $a_1 = a_2 = a_3 = a_4 = 18nM^{-1}s^{-1}$, $d_1 = d_2 = d_3 = d_4 = 2400s^{-1}$, $k_{on} = 10nM^{-1}s^{-1}$, $k_{off} = 10s^{-1}$. Ideal system ($Z_{ideal}$): $X_{T1} = X_{T2} = M_T = p_T = 0$. Isolated system ($X_{2,is}^*$) with low $X_{T1}$, $M_T$: $X_{T1} = M_T = 3nM$, $X_{T2} = 1200nM$, $p_T = 0$. Actual system ($X_2^*, Z$) with low $X_{T1}$: $X_{T1} = M_T = 3nM$, $X_{T2} = 1200nM$, $p_T = 100nM$. Isolated system ($X_{2,is}^*$) with high $X_{T1}$, $M_T$: $X_{T1} = M_T = 300nM$, $X_{T2} = 1200nM$, $p_T = 0$. Actual system ($X_2^*, Z$) with high $X_{T1}$: $X_{T1} = M_T = 300nM$, $X_{T2} = 1200nM$, $p_T = 0$. |
| Cascade of phosphorylation cycles (Fig. 6) | $k(t) = 0.01(1 + \sin(0.05t))nM.s^{-1}, \delta = 0.01s^{-1}, a_1 = a_2 = 18(nM.s)^{-1}, d_1 = d_2 = 2400s^{-1}, k_1 = k_2 = 600s^{-1}$. Ideal system ($Z_{ideal}$): $X_{T1} = X_{T2} = M_T = p_T = 0$. Isolated system ($X_{2, is}^*$): $X_{T1} = 3nM, X_{T2} = 1000nM, M_T = 54nM, p_T = 0$. Actual system ($Z, X_2^*$): $X_{T1} = 3nM, X_{T2} = 1000nM, M_T = 54nM, p_T = 100nM$. |
| Phosphotransfer with phospho-donor undergoing autophosphorylation as input (Fig. 7) | $k(t) = 0.01(1 + \sin(0.05t)), \delta = 0.01s^{-1}, k_3 = 600s^{-1}, a_1 = a_2 = a_3 = 18nM^{-1}s^{-1}, d_1 = d_2 = d_3 = 2400s^{-1}, k_{on} = 10nM^{-1}s^{-1}, k_{off} = 10s^{-1}, X_{T2} = 1200nM$. Ideal system ($X_{1, ideal}$): $\pi_1 = M_T = p_T = 0$. Isolated system ($X_{2, is}^*$) with low $\pi_1$: $\pi_1 = 30nM, M_T = 9nM, p_T = 0$. Actual system ($Z, X_2^*$) with low $\pi_1$: $\pi_1 = 30nM, M_T = 9nM, p_T = 100nM$. Isolated system ($X_{2, is}^*$) with high $\pi_1$: $\pi_1 = 1500nM, M_T = 420nM, p_T = 0$. Actual system ($Z, X_2^*$) with high $\pi_1$: $\pi_1 = 1500nM, M_T = 420nM, p_T = 100nM$. |
| Single cycle with substrate as input (Fig. 8) | $k(t) = 0.01(1 + \sin(0.05t)), \delta = 0.01s^{-1}, k_1 = k_2 = 600s^{-1}, a_1 = a_2 = 18nM^{-1}s^{-1}, d_1 = d_2 = 2400s^{-1}, k_{on} = 10nM^{-1}s^{-1}, k_{off} = 10s^{-1}$. Ideal system ($X_{ideal}$): $Z_T = M_T = p_T = 0$. Isolated system ($X_{is}^*$) with low $Z_T, M_T$: $Z_T = M_T = 100nM, p_T = 0$. Actual system ($X, X^*$) with low $Z_T, M_T$: $Z_T = M_T = p_T = 100nM$. Isolated system ($X_{is}^*$) with high $Z_T, M_T$: $Z_T = M_T = 1000nM, p_T = 0$. Actual system ($X, X^*$) with low $Z_T, M_T$: $Z_T = M_T = 1000nM, p_T = 100nM$. |
| Double cycle with substrate as input (Fig. 9) | $k(t) = 0.01(1 + \sin(0.05t)), \delta = 0.01s^{-1}, k_1 = k_2 = k_3 = k_4 = 600s^{-1}, a_1 = a_2 = a_3 = a_4 = 18nM^{-1}s^{-1}, d_1 = d_2 = d_3 = d_4 = 2400s^{-1}, k_{on} = 10nM^{-1}s^{-1}, k_{off} = 10s^{-1}$. Ideal system ($X_{ideal}$): $Z_T = M_T = p_T = 0$. Isolated system ($X_{is}^*$) with low $Z_T, M_T$: $Z_T = M_T = 150nM, p_T = 0$. Actual system ($X, X^*$) with low $Z_T, M_T$: $Z_T = M_T = 150nM, p_T = 100nM$. Isolated system ($X_{is}^*$) with high $Z_T, M_T$: $Z_T = M_T = 1000nM, p_T = 0$. Actual system ($X, X^*$) with low $Z_T, M_T$: $Z_T = M_T = 1000nM, p_T = 100nM$. |

### 5.3 Single cycle with kinase input

The reactions for this system are:

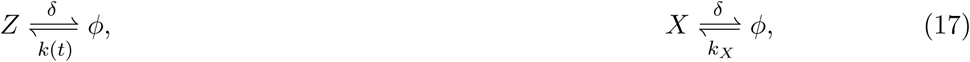

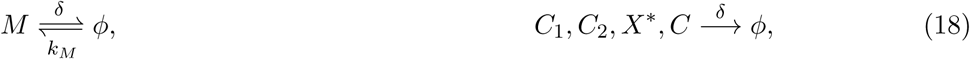

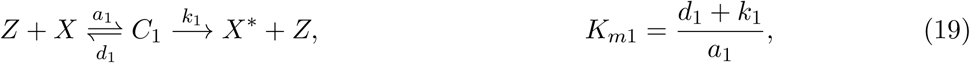

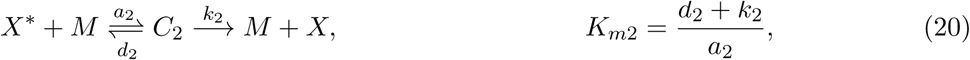

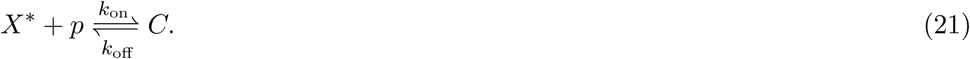

Using reaction-rate equations, and the conservation law for the promoter *p_T_ = p* + *C*, the ODEs for this system are then:

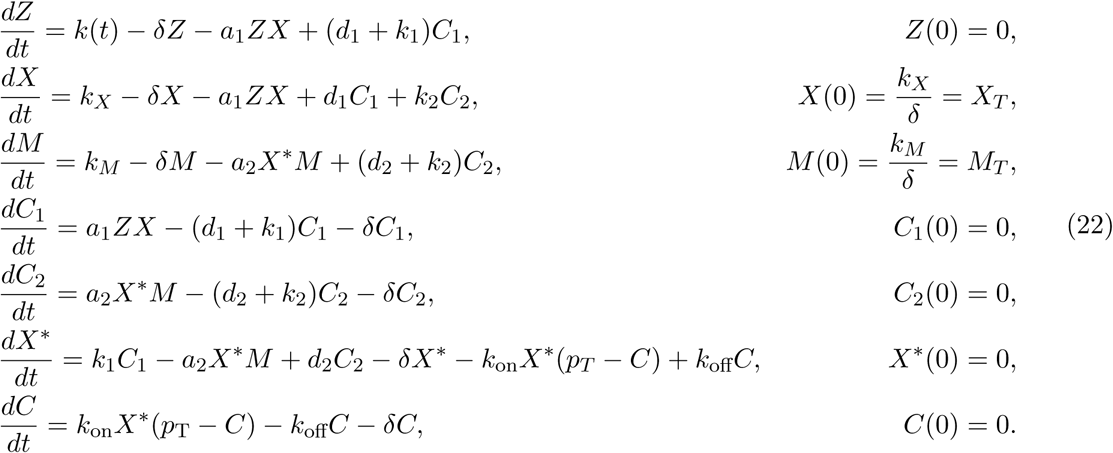

For the system defined by (22), let *M*_T_ = *M* + *C*_2_. Then the dynamics of *M_T_* are 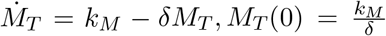. This gives a constant 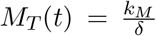. The variable *M* = *M_T_* − *C*_2_ is then eliminated from the system. Similarly, we define *X_T_* = *X* + *C*_1_ + *C*_2_ + *X*^*^ + *C*, whose dynamics become 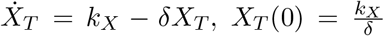. Thus, 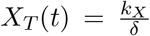 is a constant. The variable *X* = *X_T_ − C*_1_ − *C*_2_ − *X*^*^ − *C* can then be eliminated from the system. Further, we non-dimensionalize *C* with respect to *p_T_*, such that c 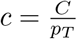 The system thus reduces to:

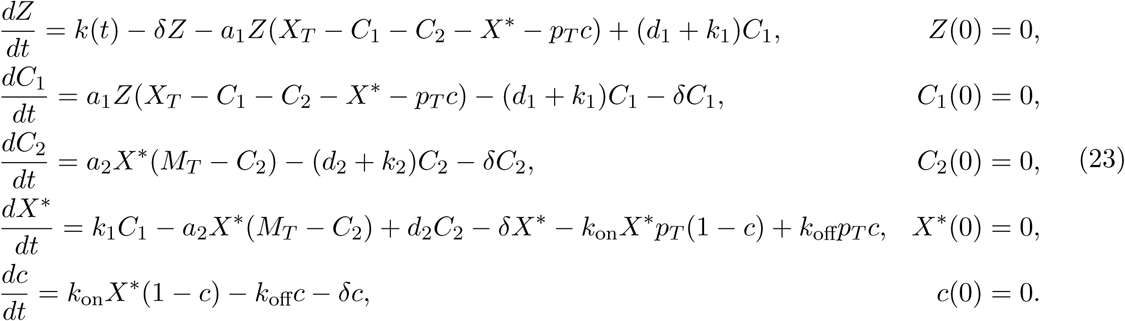

#### Step 1

Based on eqns. (23), we bring the system to form (1) as shown in Table 2.

**Table 2.** System variables, functions and matrices for a double phosphorylation cycle with the kinase for both cycles as input brought to form (1).

|  |  |  |  |
| --- | --- | --- | --- |
| $U$ | $Z$ | $v$ | $c$ |
| $\underline{X}$ | $[C_1 \ C_2 \ X^*]_{3 \times 1}^T$ | $Y, I$ | $X^*, [0 \ 0 \ 1]_{1 \times 3}$ |
| $G_1$ | $\max \left\{ \frac{a_1 X_T}{\delta}, \frac{d_1}{\delta}, \frac{k_1}{\delta}, \frac{a_2 X_T}{\delta}, \frac{d_2}{\delta}, \frac{k_2}{\delta} \right\}$ | $G_2$ | $\max \left\{ \frac{k_{\text{on}} p_T}{\delta}, \frac{k_{\text{off}}}{\delta} \right\}$ |
| $f_0(U, R\underline{X}, S_1 v, t)$ | $k(t) - \delta Z - \delta C_1$ | $s(\underline{X}, v)$ | $\frac{1}{G_2} (k_{\text{on}} X^* (1 - c) - k_{\text{off}} c - \delta c)$ |
| $r(U, \underline{X}, S_2 v)$ | $\frac{1}{G_1} \left[ -a_1 Z X_T (1 - \frac{X^*}{X_T} - \frac{C_1}{X_T} - \frac{C_2}{X_T} - \frac{p_T}{X_T} c) + (d_1 + k_1) C_1 + \delta C_1 \right]_{1 \times 1}$ | | |
| $f_1(u, \underline{x}, S_3 v)$ | $\frac{1}{G_1} \begin{bmatrix} 0 \\ a_2 X^* (M_T - C_2) - (d_2 + k_2) C_2 - \delta C_2 \\ k_1 C_1 - a_2 M_T (X^* + \frac{\delta p_T}{a_2 M_T} c) + a_2 X^* C_2 + d_2 C_2 - \delta X^* \end{bmatrix}_{3 \times 1}$ | | |
| $A$ | $1$ | $D$ | $1$ |
| $B$ | $[-1 \ 0 \ 0]_{3 \times 1}^T$ | $C$ | $[0 \ 0 \ -p_T]_{3 \times 1}^T$ |
| $R$ | $[1 \ 0 \ 0]_{1 \times 3}$ | $S_1$ | $0$ |
| $S_2$ | $\frac{p_T}{X_T}$ | $S_3$ | $\frac{\delta p_T}{a_2 M_T}$ |
| $T$ | $1$ | $M$ | $[1 \ 0 \ 0]_{1 \times 3}$ |
| $Q$ | $\mathbb{I}_{3 \times 3}$ | $P$ | $[0 \ 0 \ p_T]_{3 \times 1}^T$ |

#### Step 2

We now solve for <u>Ψ</u>, *ϕ* and <u>Γ</u> as defined by Assumptions 5, 6 and 7. The other terms required for Step 2 are noted in Table 2.

Solving for *<u>X</u>* = <u>Ψ</u>(*U*, *υ*) setting (*Br* + *f*_1_)_3×1_ = 0, we have:

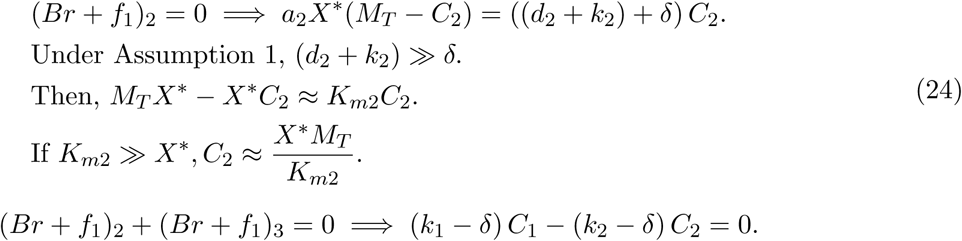

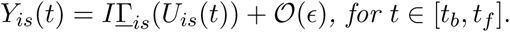

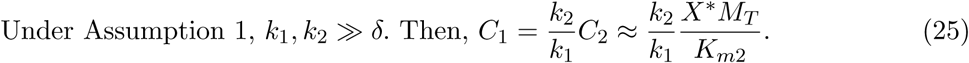

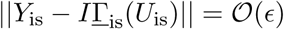

Note that as the input *Z* becomes very large, the output *X*^*^ saturates to 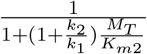 Since this violates condition (iii) of Def. 1, we must have 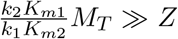. This gives a range of input z for which condition (iii) of Def. 1 is satisfied. Once the input increases so that 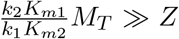 are no longer satisfied, condition (iii) does not hold. Under these conditions, the expression for *X*^*^ is then:

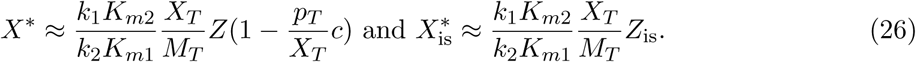

From (24)-(26), we have <u>Ψ</u>(*U*, *υ*) given by:

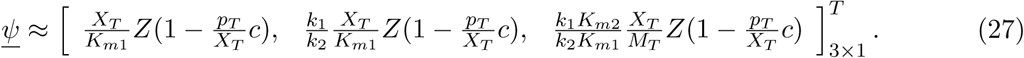

Solving for *ϕ* by setting *s*(*<u>X</u>*; *υ*) = 0, we have:

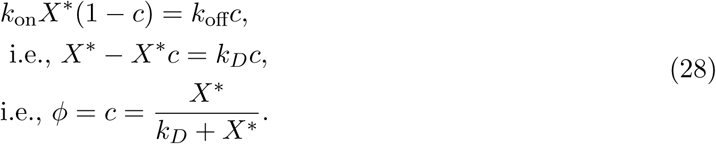

We can use (27) and (28) to find <u>Γ</u> as defined in Remark 1, and find that it satisfies Assumption 7. We then state without proof the following claims for this system:

##### Claim 1

*For the matrix B and functions r, f*_1_ *and s defined in Table 2, Assumption 3 is satisfied for this system.*

##### Claim 2

*For the functions f*_***0***_ *and <u>r</u> and matrices R, S*_1_ *and A defined in Table 2, and the functions <u>γ</u> and ϕ as found above, Assumption 9 is satisfied for this system.*

For matrices *T, Q, M, P* defined in Table 2, we see that Assumption 4 is satisfied. Further, for <u>Ψ</u> and *ϕ* defined by (27) and (28), Assumption 5 and 6 are satisfied. Thus, Theorems 1, 2 and 3 can be applied to this system to check if the system can transmit unidirectional signals according to Definition 1 by varying *X_T_* and *M_T_*.

#### Step 3 and Test (i)

Retroactivity to the input: Using Theorem 1, we see that since *S*_1_ = 0, Further, 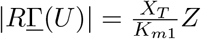. Evaluating the final term, we see that:

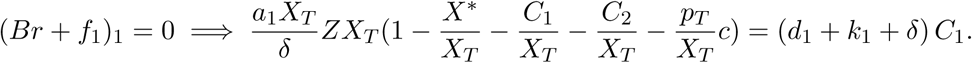

Thus, for a small retroactivity to the input, we must have small 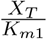.

#### Step 4 and Test (ii)

Retroactivity to the output: We see that *S*_1_ = 0. Further, the term 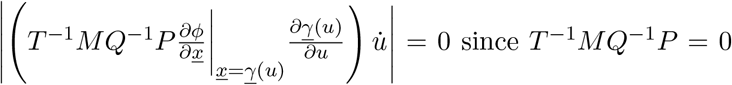 from Table 2. Further, we see that 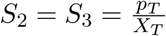 must be small. Thus, to decrease the retroactivity to input, *X_T_* must be increased.

#### Step 5 and Test (iii)

Input-output relationship: Evaluating *Y_is_* = *I<u>Γ</u>*_*is*_ + 𝒪(ϵ). Under Remark 1, 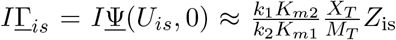 from (27). Thus, the dimensionless input-output behavior is approximately linear. Thus, from Def. 1(iii) we have that *m* = 1 and 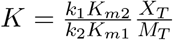 which can be tuned by tuning the substrate and phosphatase concentrations *X_T_, M*_T_.

**Test (iv)** fails since Tests (i) and (ii) have opposing requirements from the total protein concentrations.

### 5.4 Double cycle with input as kinase of both phosphorylations

The reactions for this system are then:

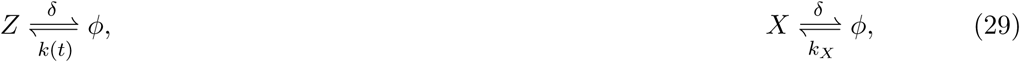

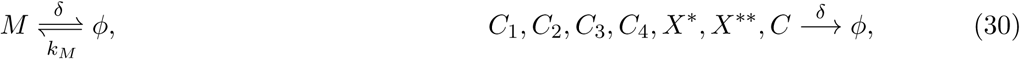

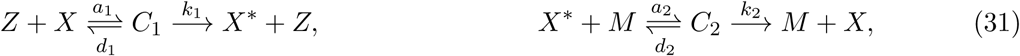

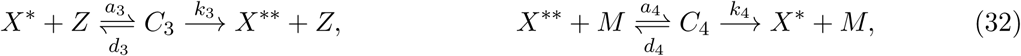

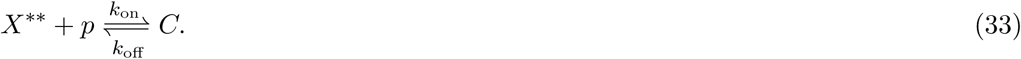

Using the reaction-rate equations, the ODEs for this system are:

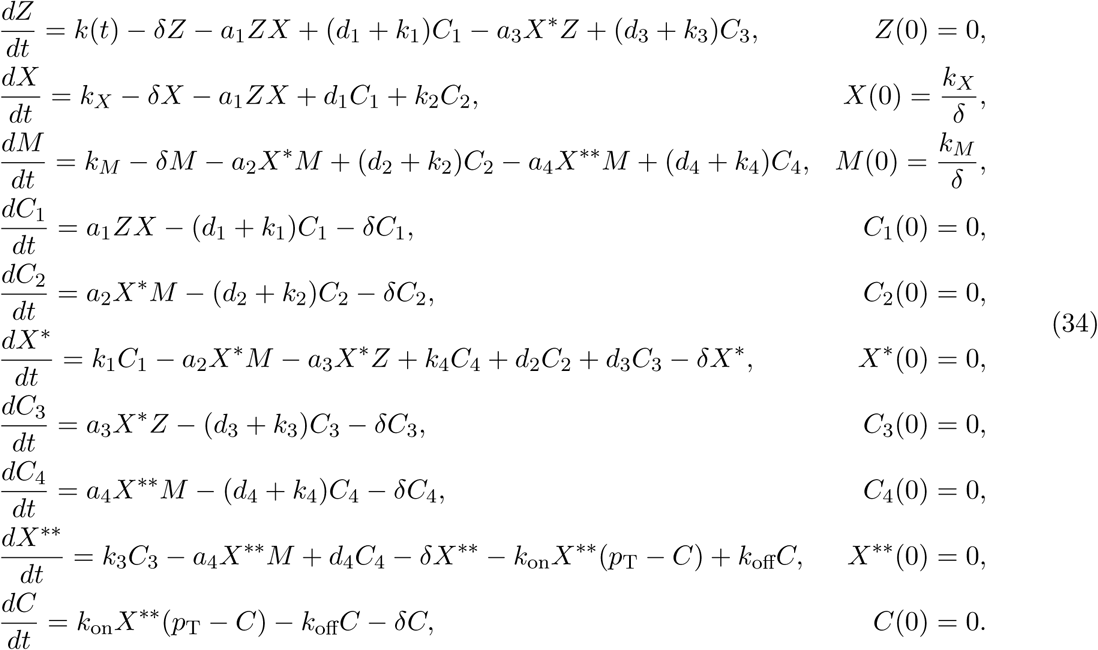

For system (34), let *M_T_* = *M* + *C*_2_ + *C*_4_. Then its dynamics are 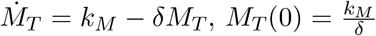. This gives a constant 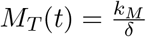. The variable *M* = *M_T_*− *C*_2_ − *C*_4_ can then be eliminated from the system. Similarly, defining *X_T_* = *X* + *C*_1_ + *C*_2_ + *X*^*^ + *C*_3_ + *C*_4_ + *X*^**^ + *C* gives a constant 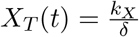, and *X* can be eliminated from the system as *X* = *X_T_*−*X*^*^ −*X*^**^ −*C*_1_ −*C*_2_−*C*_3_−*C*_4_−*C*. Further, we define 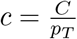 which the dimensionless form of *C*. The system then reduces to:

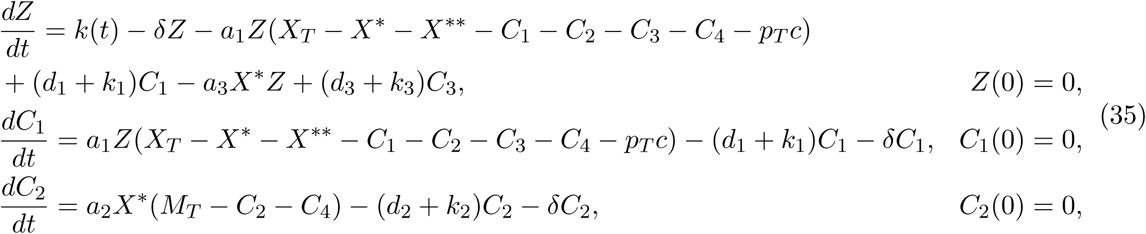

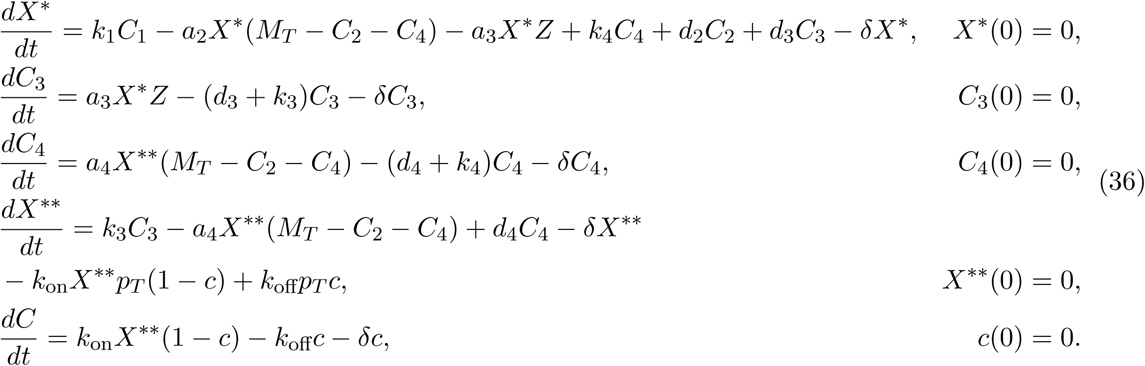

This system (35), (36) is brought to form (1) as shown in Table 3.

**Table 3.** System variables, functions and matrices for a double phosphorylation cycle with the kinase for both cycles as input brought to form (1).

|  |  |  |  |
| --- | --- | --- | --- |
| $U$ | $Z$ | $v$ | $c$ |
| $\underline{x}$ | $[C_1 \ C_2 \ X^* \ C_3 \ C_4 \ X^{**}]_{6 \times 1}^T$ | $Y, I$ | $X^{**}, [0 \ 0 \ 0 \ 0 \ 0 \ 1]_{1 \times 6}$ |
| $G_1$ | $\max \left\{ \frac{a_1 X_T}{\delta}, \frac{d_1}{\delta}, \frac{k_1}{\delta}, \frac{a_2 M_T}{\delta}, \frac{d_2}{\delta}, \frac{k_2}{\delta}, \frac{a_3 X_T}{\delta}, \frac{d_3}{\delta}, \frac{k_3}{\delta}, \frac{a_4 M_T}{\delta}, \frac{d_4}{\delta}, \frac{k_4}{\delta} \right\}$ | $G_2$ | $\max \left\{ \frac{k_{\text{on}} p_T}{\delta}, \frac{k_{\text{off}}}{\delta} \right\}$ |
| $f_0(U, R\underline{X}, S_1 v, t)$ | $k(t) - \delta Z - \delta C_1 - \delta C_3$ | $s(\underline{X}, v)$ | $\frac{1}{G_2} (k_{\text{on}} X^{**} (1 - c) - k_{\text{off}} c - \delta c)$ |
| $r(U, \underline{X}, S_2 v)$ | $\frac{1}{G_1} \begin{bmatrix} -a_1 Z X_T (1 - \frac{X^*}{X_T} - \frac{X^{**}}{X_T} - \frac{C_1}{X_T} - \frac{C_2}{X_T} - \frac{C_3}{X_T} - \frac{C_4}{X_T} - \frac{p_T}{X_T} c) + (d_1 + k_1) C_1 + \delta C_1 \\ -a_3 Z X^* + (d_3 + k_3) C_3 + \delta C_3 \end{bmatrix}_{2 \times 1}$ | | |
| $f_1(U, \underline{X}, S_3 v)$ | $\frac{1}{G_1} \begin{bmatrix} 0 \\ a_2 X^* (M_T - C_2 - C_4) - (d_2 + k_2) C_2 - \delta C_2 \\ k_1 C_1 - a_2 X^* (M_T - C_2 - C_4) - a_3 X^* Z + k_4 C_4 + d_2 C_2 + d_3 C_3 - \delta X^* \\ 0 \\ a_4 X^{**} (M_T - C_2 - C_4) - (d_4 + k_4) C_4 - \delta C_4 \\ k_3 C_3 - a_4 M_T (X^{**} + \frac{\delta p_T}{a_4 M_T} c) a_4 X^{**} (C_2 + C_4) + d_4 C_4 - \delta X^{**} \end{bmatrix}_{6 \times 1}$ | | |
| $A$ | $[1 \ 1]_{1 \times 2}$ | $D$ | 1 |
| $B$ | $\begin{bmatrix} -1 & 0 & 0 & 0 & 0 & 0 \\ 0 & 0 & 0 & -1 & 0 & 0 \end{bmatrix}_{6 \times 2}^T$ | $C$ | $[0 \ 0 \ 0 \ 0 \ 0 \ -p_T]_{6 \times 1}^T$ |
| $R$ | $[1 \ 0 \ 0 \ 1 \ 0 \ 0]_{1 \times 6}$ | $S_1$ | 0 |
| $S_2$ | $\frac{p_T}{X_T}$ | $S_3$ | $\frac{\delta p_T}{a_4 M_T}$ |
| $T$ | 1 | $M$ | $[1 \ 0 \ 0 \ 1 \ 0 \ 0]_{1 \times 6}$ |
| $Q$ | $\mathbb{I}_{6 \times 6}$ | $P$ | $[0 \ 0 \ 0 \ 0 \ 0 \ p_T]_{6 \times 1}^T$ |

#### Steps 1 and 2

For the system brought to form (1) as seen in Table 3, we now solve for <u>Ψ</u> and *ϕ* as defined by Assumptions 5 and 6.

Solving for *<u>X</u>* = <u>Ψ</u> by setting (*Br* + *f*_1_)_6×1_ = 0, we have:

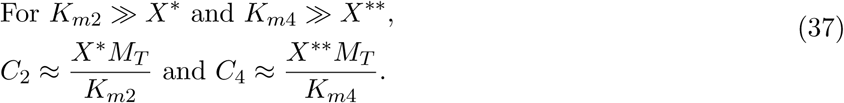

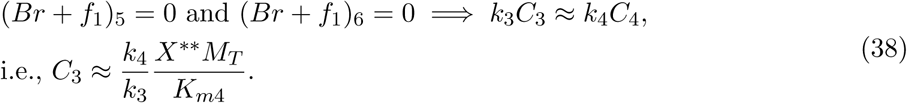

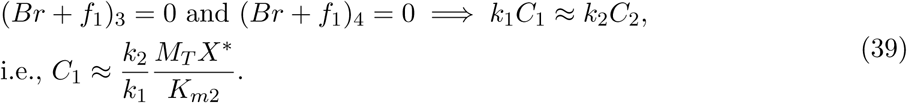

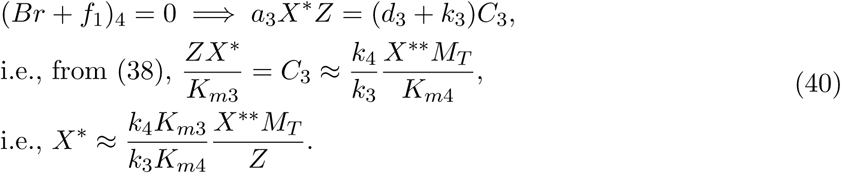

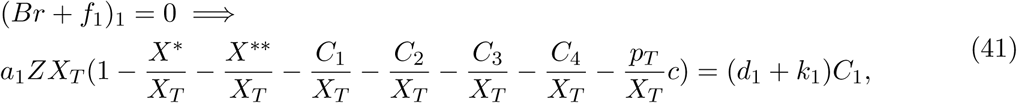

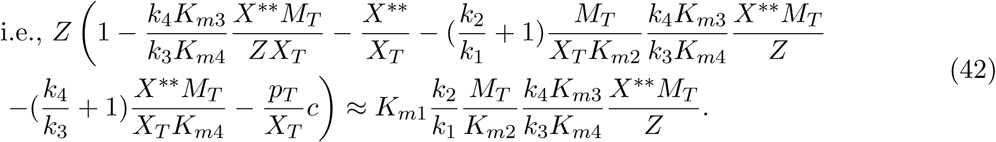

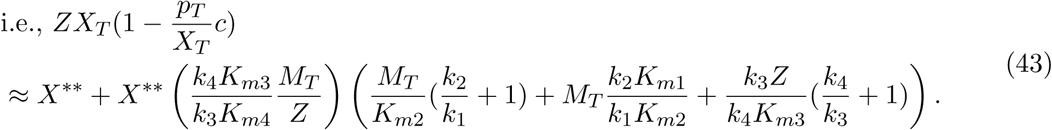

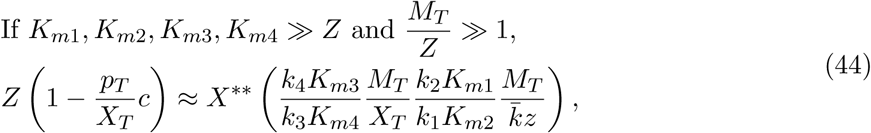

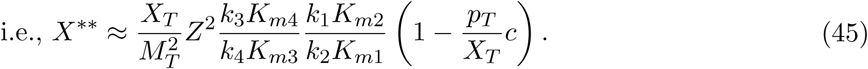

Thus, from (37)-(45), we have the function <u>Ψ</u>(*U*; *υ*):

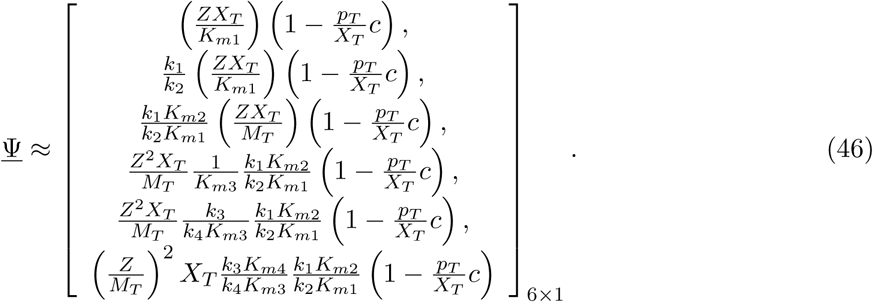

Solving for *ϕ* by setting *s*(*<u>X</u>*; *υ*) = 0, we have:

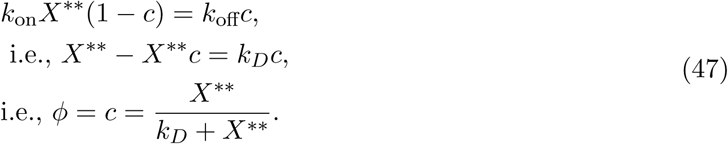

We can use (46) and (47) to find <u>Γ</u> as defined in Remark 1, and find that it satisfies Assumption 7. We then state the following claims without proof for this system:

##### Claim 3

*For the matrix B and the functions r, f*_**1**_ *and s defined in Table 3, Assumption 3 is satisfied for large K*_*m*1_, *K*_*m*2_, *K*_*m*3_, *K*_*m*4_.

##### Claim 4

*For the functions f*_0_ *and <u>r</u> and matrices R, S*_***1***_ *and A defined in Table 3, and the functions <u>γ</u>and ϕ as found above, Assumption 9 is satisfied for this system.*

For matrices *T, Q, M, P* defined in Table 3, we see that Assumption 4 is satisfied. Further, for <u>Ψ</u> and *ϕ* defined by (46) and (47), Assumptions 5 and 6 are satisfied. Thus, Theorems 1, 2 and 3 can be applied to this system.

#### Results: Step 3 and Test (i)

Retroactivity to the input: We see that since *S*_1_ = 0 from Table 3. Further, 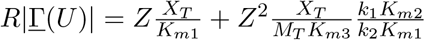. For the final term we evaluate:

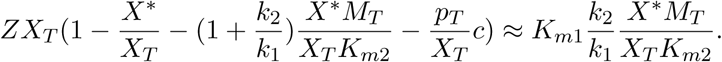

Thus, for small retroactivity to the input, we must have small 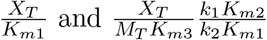.

#### Step 4 and Test (ii)

Retroactivity to the output: From Table 3, we see that *S*_**1**_ = 0. Further, since *T^-^*^***1***^*MQ^-1^P* = 0, the expression 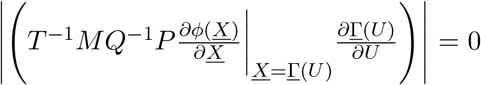. *S*_3_ = 0, thus, we must have a small 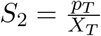. Thus, for a small retroactivity to the output, we must have a large *X*_T_.

#### Step 5 and Test (iii)

Input-output relationship: From eqn. (46), we have that:

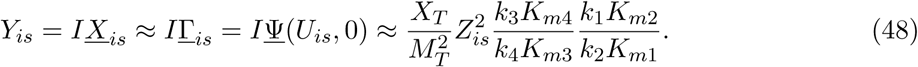

**Test (iv)** fails since Tests (i) and (ii) have opposing requirements from the total protein concentrations.

### 5.5 Phosphotransfer with kinase as input

The reactions for this system are:

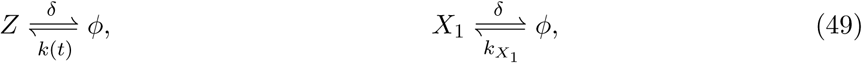

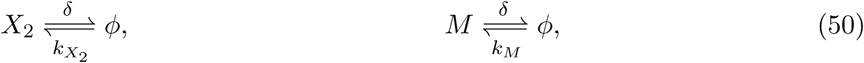

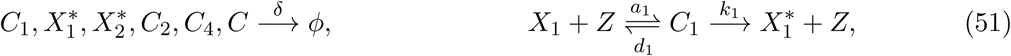

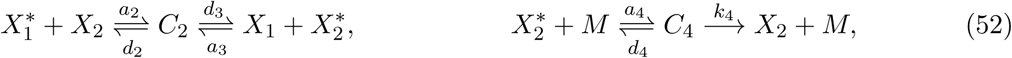

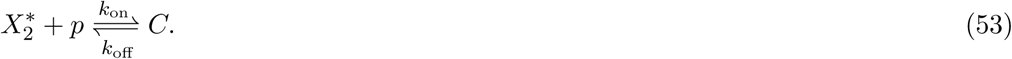

The ODEs based on the reaction rate equations are:

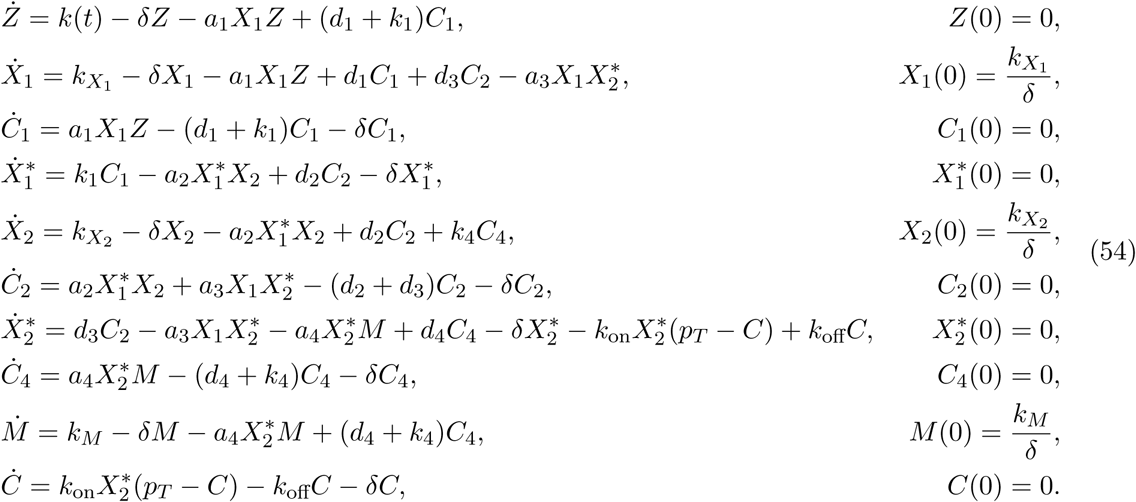

For (54), define 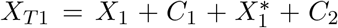. Then, 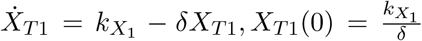. Thus, 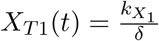 is a constant at all time *t* > 0. Similarly, 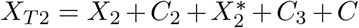 is a constant with 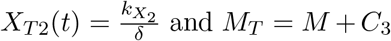 is a constant with 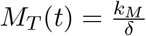 for all time *t* > 0. Thus, the variables, 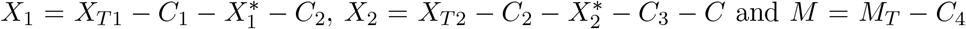 can be eliminated from the system. Further, we define 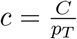. The reduced system is then:

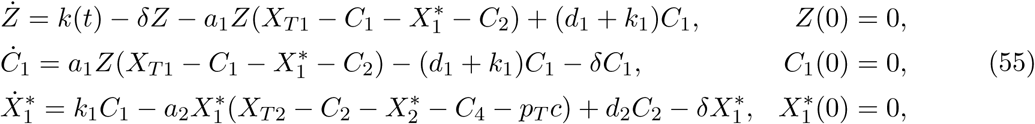

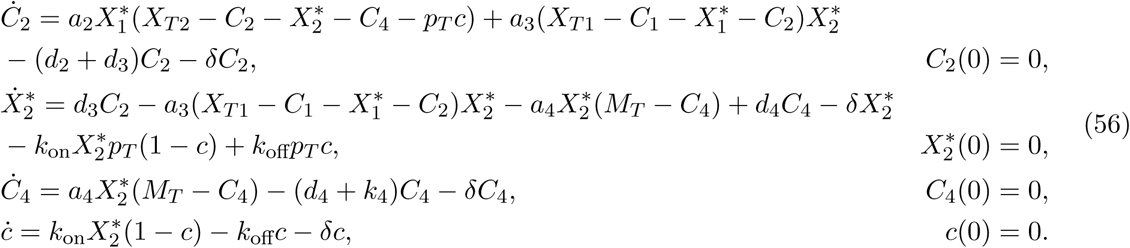

#### Step 1

This system (55), (56) is brought to form (1) as shown in Table 4.

**Table 4.** System variables, functions and matrices for a phosphotransfer system with kinase as input brought to form (1).

|  |  |  |  |
| --- | --- | --- | --- |
| $U$ | $Z$ | $v$ | $c$ |
| $\underline{X}$ | $[C_1 \ X_1^* \ C_2 \ X_2^* \ C_4]_{5 \times 1}^T$ | $Y, I$ | $X_2^*, [0 \ 0 \ 0 \ 1 \ 0]_{1 \times 5}$ |
| $G_1$ | $\max \left\{ \frac{a_1 X_{T1}}{\delta}, \frac{d_1}{\delta}, \frac{k_1}{\delta}, \frac{a_2 X_{T2}}{\delta}, \frac{d_2}{\delta}, \frac{d_3}{\delta}, \frac{a_3 X_{T1}}{\delta}, \frac{a_4 M_T}{\delta}, \frac{d_4}{\delta}, \frac{k_4}{\delta} \right\}$ | $G_2$ | $\max \left\{ \frac{k_{on} p_T}{\delta}, \frac{k_{off}}{\delta} \right\}$ |
| $f_0(U, R\underline{X}, S_1 v, t)$ | $k(t) - \delta Z - \delta C_1$ | $s(\underline{X}, v)$ | $\frac{1}{G_2} (k_{on} X_2^* (1 - c) - k_{off} c - \delta c)$ |
| $r(U, \underline{X}, S_2 v)$ | $\frac{1}{G_1} (-a_1 Z (X_{T1} - C_1 - X_1^* - C_2) + (d_1 + k_1) C_1 + \delta C_1)$ | | |
| $f_1(U, \underline{X}, S_3 v)$ | $\frac{1}{G_1} \begin{bmatrix} 0 \\ k_1 C_1 - a_2 X_1^* X_{T2} (1 - \frac{C_2}{X_{T2}} - \frac{X_2^*}{X_{T2}} - \frac{C_4}{X_{T2}} - \frac{p_T}{X_{T2}} c) + d_2 C_2 - \delta X_1^*, \\ a_2 X_1^* X_{T2} (1 - \frac{C_2}{X_{T2}} - \frac{X_2^*}{X_{T2}} - \frac{C_4}{X_{T2}} - \frac{p_T}{X_{T2}} c) - (d_2 + d_3) C_2 + a_3 (X_{T1} - C_1 - X_1^* - C_2) X_2^* - \delta C_2, \\ d_3 C_2 - a_4 X_2^* (M_T - C_4) + d_4 C_4 + a_3 (C_1 + X_1^* + C_2) X_2^* - a_3 X_{T1} (X_2^* + \frac{\delta p_T}{a_3 X_{T1}} c) - \delta X_2^*, \\ a_4 X_2^* (M_T - C_4) - (d_4 + k_4) C_4 - \delta C_4 \end{bmatrix}_{5 \times 1}$ | | |
| $A$ | $1$ | $D$ | $1$ |
| $B$ | $[-1 \ 0 \ 0 \ 0 \ 0]_{5 \times 1}^T$ | $C$ | $[0 \ 0 \ 0 \ -p_T \ 0]_{5 \times 1}^T$ |
| $R$ | $[1 \ 0 \ 0 \ 0 \ 0]_{1 \times 5}$ | $S_1$ | $0$ |
| $S_2$ | $0$ | $S_3$ | $\frac{p_T}{X_{T2}}, \frac{\delta p_T}{a_3 X_{T1}}$ |
| $T$ | $1$ | $M$ | $[1 \ 0 \ 0 \ 0 \ 0]_{1 \times 5}$ |
| $Q$ | $\mathbb{I}_{5 \times 5}$ | $P$ | $[0 \ 0 \ 0 \ p_T \ 0]_{5 \times 1}^T$ |

#### Step 2

We now solve for the functions <u>Ψ</u> and *ϕ* as defined by Assumptions 5 and 6.

Solving for *<u>X</u>* = <u>Ψ</u> by setting (*Br* + *f*_1_)_5_ = 0, we have:

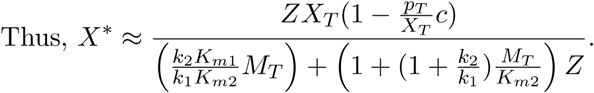

Solving the above 2 simultaneously, we obtain:

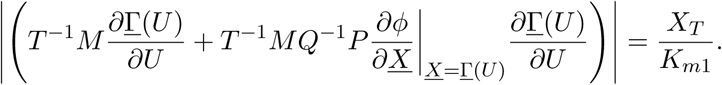

Thus, we have the function <u>Ψ</u>(*U*, *υ*):

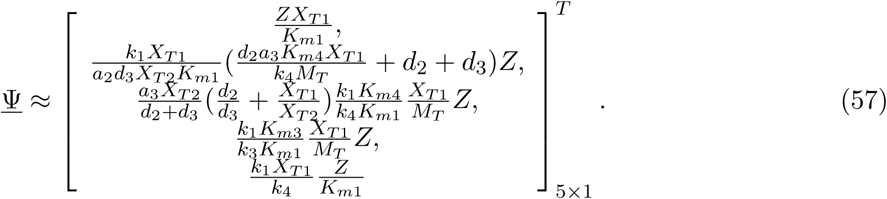

Solving for *ϕ* by setting *s*(*<u>X</u>*; *υ*) = 0, we have:

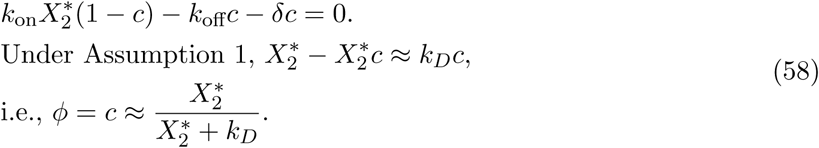

Finding <u>Γ</u> from (57) and (58) under Remark 1, we see that it satisfies Assumption 7. For matrices *T, Q, M* and *P* as seen in Table 4, we see that Assumption 4 is satisfied. Functions *f*_0_ and *<u>r</u>* in Table 4 satisfy Assumptions 8. For the functions <u>Ψ</u>, *ϕ* and <u>Γ</u>, Assumptions 5, 6 and 7 are satisfied. We also claim without proof that Assumptions 3 and 9 are satisfied for this system. Theorems 1, 2 and 3 can then be applied to this system.

#### Results: Step 3 and Test (i)

Retroactivity to the input: *S*_1_ = 0 from Table 4. Further, 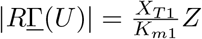. Finally, we evaluate the following expression:

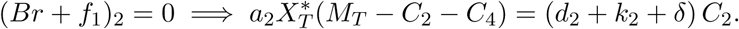

Thus, for small retroactivity to the input, we must have small 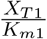.

#### Step 4 and Test (ii)

Retroactivity to the output: We see from Table 4 that *S*_1_ =0 and further, *T*^−1^*MQ*^−1^*P* = 0. Since *S*_2_ = 0, we must have a small 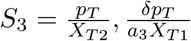. Thus, for a small retroactivity to the output, we must have a large *X*_*T*2_ and 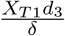 compared to *p_T_*.

#### Step 5 and Test (iii)

Input-output relationship: From (57), we see that

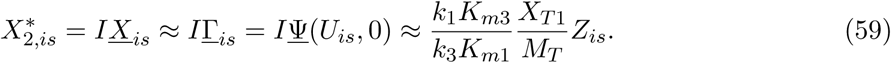

**Test (iv)** fails since Tests (i) and (ii) have opposing requirements from the total protein concentrations.

### 5.6 N-stage cascade of single phosphorylation cycles with common phosphatase

The two-step reactions for the cascade are shown below. The reactions involving species of the first cycle are given by:

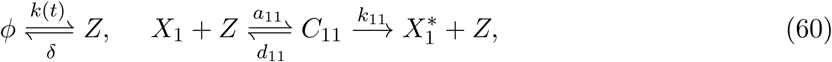

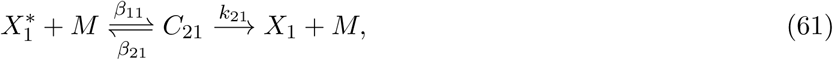

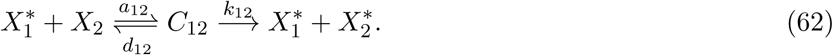

The reactions involving species of the *i*^th^ cycle, for *i* ∈ [2;*N* − 1], are given by:

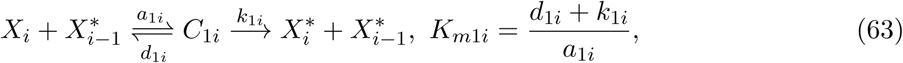

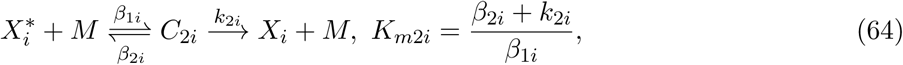

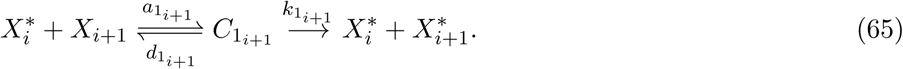

And those for the final cycle are given by:

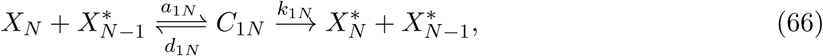

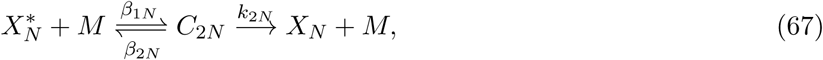

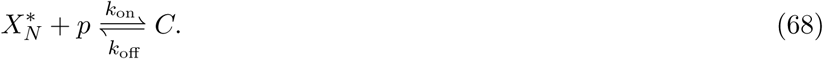

The production and dilution of the proteins and other species gives:

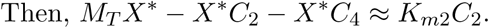

The reaction rate equations for the system are then given below, for time *t ∈* [*t_i_, t_f_*]. For the input,

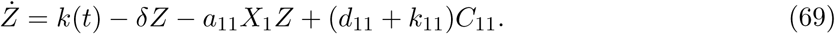

For the first cycle,

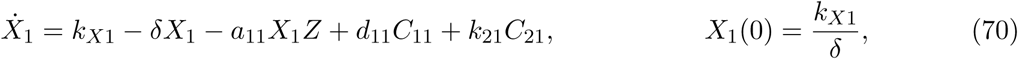

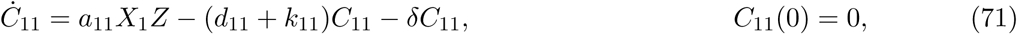

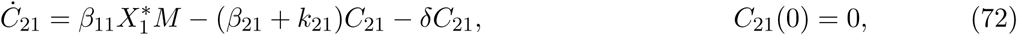

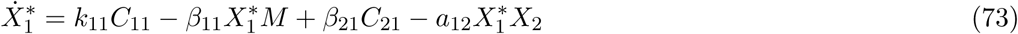

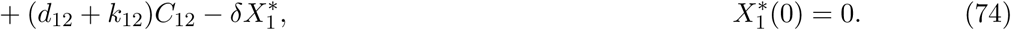

For the *i*^th^ cycle, where *i* ∈ [2, *N* − 1]:

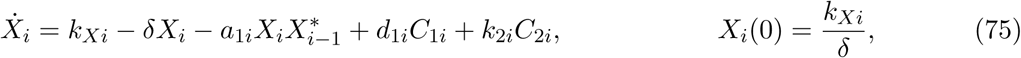

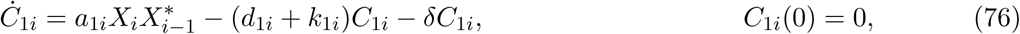

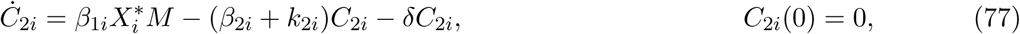

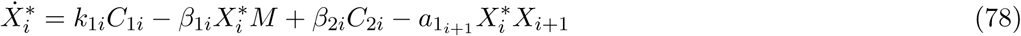

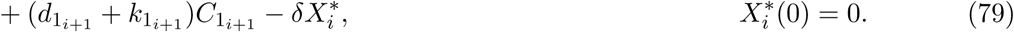

For the last, *N*^th^, cycle:

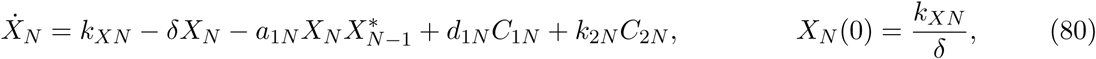

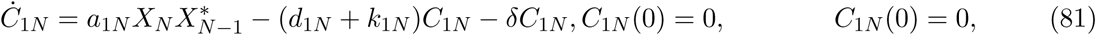

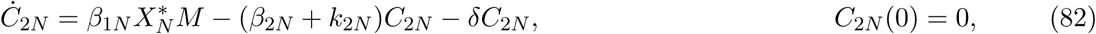

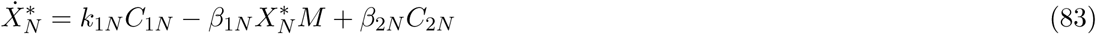

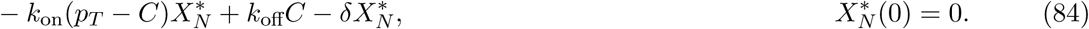

For the common phosphatase:

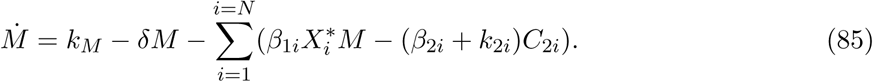

For the downstream system,

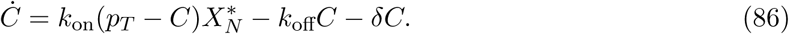

**Step 1:** Seeing that 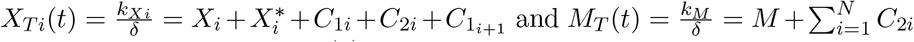 we reduce the system above to bring it to form (1) as seen in Table 5, with 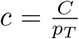. We make the following Assumptions for the system:

**Table 5.**
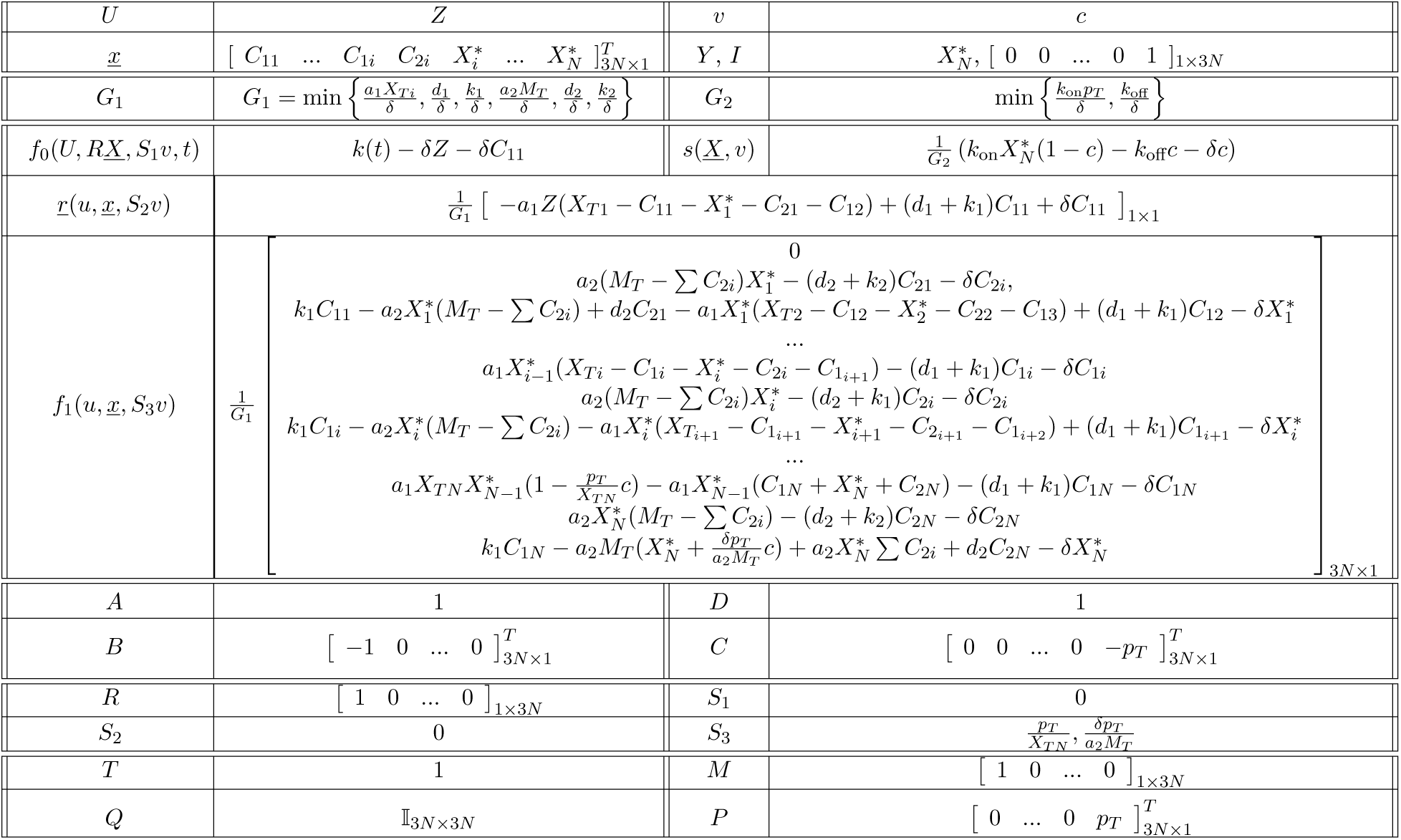
System variables, functions and matrices for an N-stage cascade of phosphorylation cycles with the kinase as input to the first cycle brought to form (1).

#### Assumption 10

All cycles have the same reaction constants, i.e., ∀_*i*_ ∈ [1, *N*], *k*_1*i*_ = *k*_1_, *k*_2*i*_ =*k*_2_, *a*_1*i*_ = *a*_1_, *β*_1*i*_ =*a*_2_, *d*_1*i*_= *d*_1_, *β_2i_=d_2._* Then, *K_m1i_*, = *Km*_1_, *Km*_2*i*_= *K*_*m*2_. Define 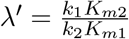

#### Assumption 11

∀*t* and ∀*i* ∈ [1, *N*], 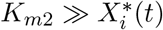.

**Step 2:** We now solve for <u>Ψ</u> by setting (*Br* + *f*_1_)_3*n*×1_ = 0. Under Assumption 11, this is given by:

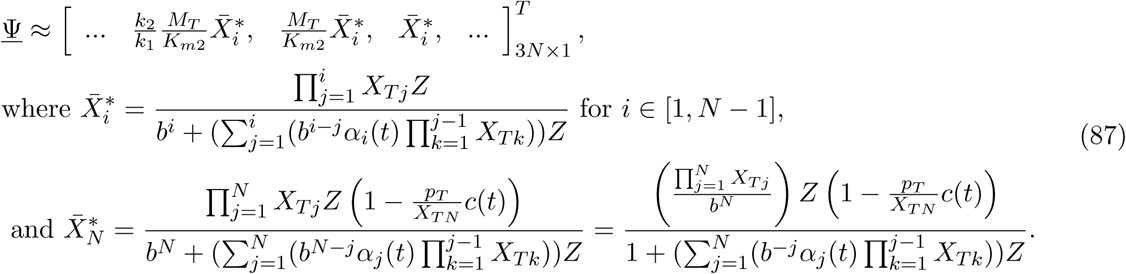

Here, *α_j_(t)* ≤ 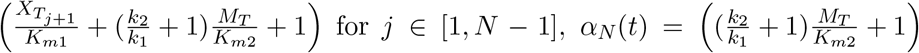 and 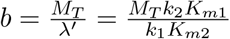.

Solving for *ϕ* by setting *s*(*<u>X</u>*, *υ*) = 0, we have:

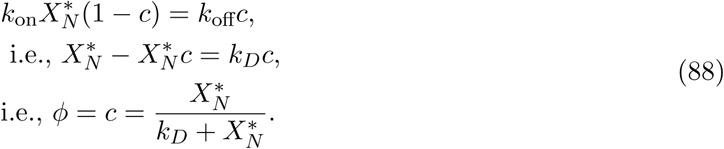

We can use (87) and (88) to find <u>Γ</u> as defined in Remark 1, and find that this satisfies Assumption 7. Note that this <u>Γ</u> differs from <u>Ψ</u> only in the last 3 terms, involving 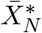 Functions <u>Ψ</u> and *ϕ* satisfy Assumptions 5 and 6. Further, from Table 5, we see that matrices *T*, *Q*, *M* and *P* satisfy Assumption 4, and functions *f*_0_ and *<u>r</u>* satisfy Assumption 8. We further assume that Assumptions 3 and 9 are satisfied for this system. Thus, Theorems 1, 2 and 3 can be applied to this system.

**Results: Step 3 and Test (i)** Retroactivity to the input: Since *S*_1_ = 0 from Table 5, under Claim 1, *h_2_* = 0. Further, 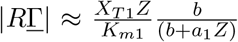, and thus, to make *h*_1_ small, we must have small 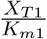. For the final term, we see that *T*^-1^*M* = [1 0 … 0] and T*^-1^MQ^-1^P* = 0. Since *T*^-1^*M* only has an entry on the first term, and since 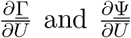 differ only in the last 3 terms, we can compute the final term using (87). This gives the following expression:

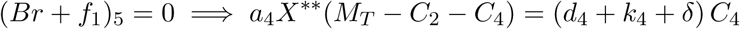

Thus, for a small retroactivity to the input, 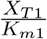 must be small.

**Step 4 and Test (ii)** Retroactivity to the output: From Table 5, we have that *S*_1_ = 0, *T*^**−**1^*MQ*^**−**1^ *P* = 0, *S*_2_ = 0, and 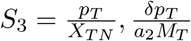 Thus, for a small retroactivity to the output, *X_TN_* and *M_T_* must be large.

**Step 5 and Test (iii)** Input-output relationship: From (87), we see that

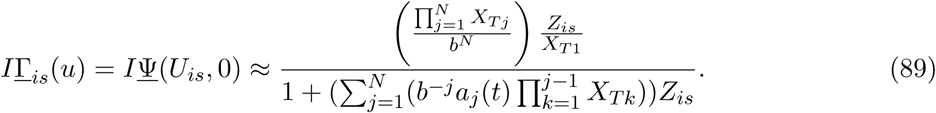

Note that 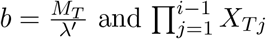 are constants, and the linear gain is 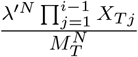

The upper bound for 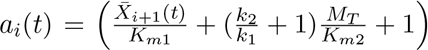, is given by seeing that the maximum value for *X*̅_*i*+1_ is *Χ*_*Ti+*1_. Let the maximum value of *Z*(*t*) for which the input-output relationship is approximately linear be *Z*_max_. We then have:

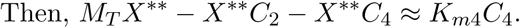

where 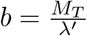. Thus, for the input-output relationship to not saturate, ∊_3_*Z*_max_ must be small. To maximize *Z*_max_, the range in which the input-output relationship is linear, we must then minimize ∊_3_. We see that, to make ∊_3_ small, we must have a large b and small *X*_*Ti*+1_. Since, to satisfy Test (ii), we saw before that *X_TN_* must be large, we have *X_Ti_*+_1_ ≤ *X_TN_*. However, as seen from the expression of *I*<u>Γ</u>_*is*_, increasing *b* also decreases the input-output gain. For simplicity, the next arguments are made to achieve unit gain for the original input *Z_is_*(*t*) and output 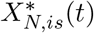. For unit gain, 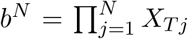. Since *X_T_j__* ≤ *X_TN_*, *j* ∊ [2, *N*], the maximum possible 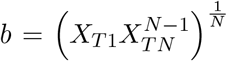, which occurs when *X_Tj_* = *X_TN_*, *j* ∊ [2, *N*]. Thus, following this argument, for unit gain and maximum linear range of the input for any N, we have *X_Tj_* = *X_TN_*, *j* ∊ [2, *N*] and 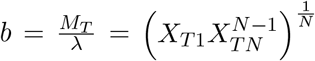. substituting 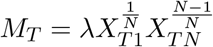, and using the geometric series sum, we obtain the following expression for ∊_3_:

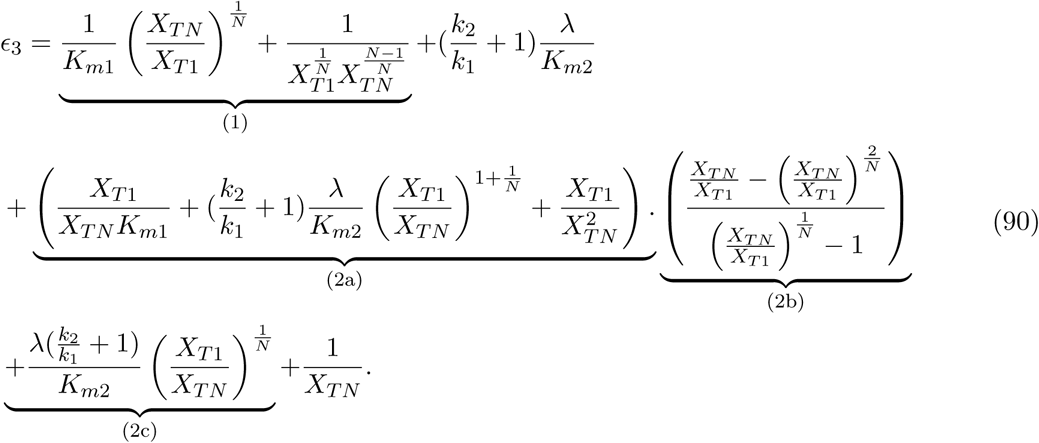

Starting from *N* = 2, we see that since *X*_*T*1_ *< X_TN_*, term (1) decreases with *N*, terms (2a), (2b) and (2c) increase with N and as *N* → ∞, ∊_3_ → ∞. The function ∊3 is continuous, and therefore, there exists an optimal number of cycles N for which the linear operating range of the input, *Z*_max_ is maximized.

To satisfy Test (iii) then, the cascade must have ∊_3_ be small, so that *m* = 1 as defined in requirement (iii) of Def. 1. As discussed above, there is an optimal *N̅* at which ∊_3_ is minimized, all other parameters remaining the same. We see from Fig. 11, that with load, the number of cycles needed increase, since *X_TN_* increases as load *ρ_T_* is increased. Note that, it may not be necessary to have *N̅* cycles to achieve a desirable result, i.e., a sufficiently large operating range. However, it is possible that no *N* is capable of producing linearity for the desired operating range, since ∊3 is bounded below.

**Figure 11:**
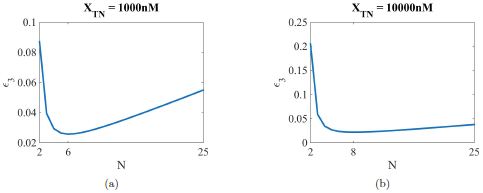
Figures showing the variation of *∊*_3_ with *N*, for different *X_TN_*. Parameter values are: *K*_*m*1_ = *K*_*m*2_ = 300*nM*, *k*_1_ = *k*_2_ = 600*s*^−1^, *λ* = 1, (a) *X_TN_* = 1000*nM*, where resulting *N̅* = 6 and (b) *X_TN_* = 10000*nM*, where resulting *N̅* = 8.

**Figure 12:**
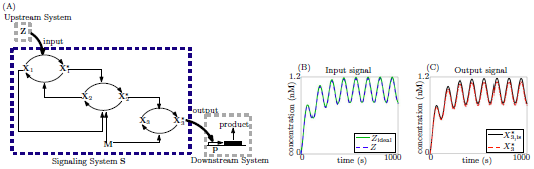
Tradeoff between small retroactivity to the input and attenuation of retroactivity to the output is overcome by a cascade of a phosphotransfer system with a single phosphorylation cycle. (A) Cascade of a phosphotransfer system that receives its input through a kinase Z phosphorylating the phosphate donor, and a phosphorylation cycle: Z phosphorylates X_1_ to 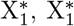 transfers the phosphate group in a reversible reaction to X_2_. 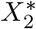 further acts as the kinase for X_3_, phosphorylating it to 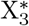, which is the output, acting on sites p in the downstream system, which is depicted as a gene expression system here. Both 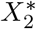 and 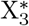 are dephosphorylated by phosphatase M. (B), (C) Simulation results for ODE model (91), (92). Simulation parameters: *k*(*t)* = 0.01(1 + *sin*(0.05*t*))*nM.s*^−1^, *δ* = 0.01*s*^−1^, *a*_1_ = *a*_2_ = *d*_3_ = *a*_4_ = *a*_5_ = *a*_6_ = 18*nM*^−1^*s*^−1^, *d*_1_ = *d*_2_ = *a*_3_ = *d*_4_ = *d*_5_ = *d_6_* = 2400*s*^−1^, *k*_1_ = *k*_4_ = *k*_5_ = *k*_6_ = 600*s*^−1^. (B) Effect of retroactivity to the input: for the ideal input *Z*_ideal_, system is simulated with X_*T*1_ = *X*_*T*2_ = *X*_*T*3_ = *M_T_* = *p_T_* = 0; for actual input *Z*, system is simulated with *X*_*T*1_ = 3*nM*, *X*_*T*2_ = 1200*nM*, *X*_*T*3_ = 1200*nM*, *M_T_* = 3*nM*, *p_T_* = 100*nM*. (C) Effect of retroactivity to the output: for the isolated output 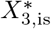, system is simulated with *X*_*T*1_ = *3nM*, *X*_*T*2_ = 1200*nM*, *X*_*T*3_ = 1200*nM*, *M_T_* = 3*nM, p_T_* = 0; for the actual output 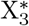, system is simulated with *X*_*T*1_ = 3*nM*, *X*_*T*2_ = 1200*nM*, *X*_*T*3_ = 1200*nM*, *M_T_* = 3*nM, pT* = 100*nM*.

**Figure 13:**
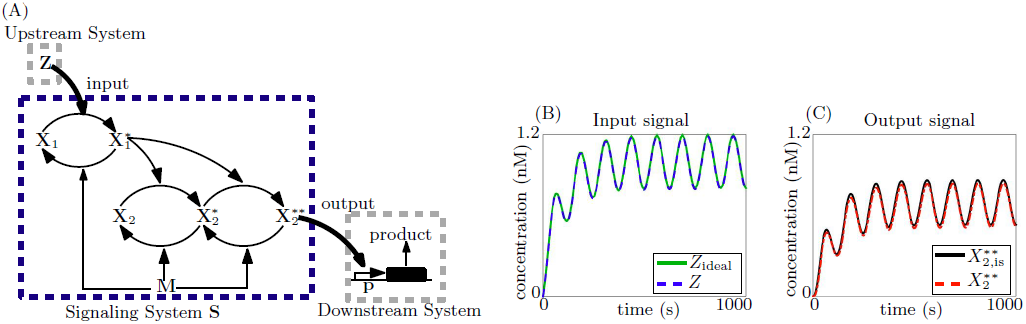
Tradeoff between small retroactivity to the input and attenuation of retroactivity to the output is overcome by a cascade of a single phosphorylation cycle and a double phosphorylation cycle. (A) Cascade of a a single phosphorylation and a double phosphorylation cycle with input kinase Z: Z phosphorylates X_1_ to 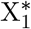, 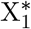 further acts as the kinase for X_2_, phosphorylating it to 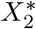 and 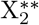, which is the output, acting on sites p in the downstream system, which is depicted as a gene expression system here. All phosphorylated proteins 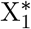, 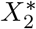 and 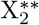 are dephosphorylated by phosphatase M. (B), (C) Simulation results for ODE model (93), (94). Simulation parameters: *k*(*t*) = 0.01(1 + *sin*(0.05*t*))*nM.s*^−1^, *δ* = 0.01*s*^−1^, *a*_1_ = *a*_2_ = *a*_3_ = *a*_4_ = *a*_5_ = *a*_6_ = 18*nM*^−1^*s*^−1^, *d*_1_ = *d*_2_ = *d*_3_ = *d*_4_ = *d*_5_ = *d*_6_ = 2400*s*^−1^, *k*_1_ = *k*_2_ = *k*_3_ = *k*_4_ = *k*_5_ = *k*_6_ = 600*s*^−1^. (B) Effect of retroactivity to the input: for the ideal input *Z*_ideal_, system is simulated with *X*_*T*1_ = *X*_*T*2_ = *X*_*T*3_ = *M_T_* = *p_T_* = 0; for actual input *Z*, system is simulated with X_*T*1_ = 3*nM*, *X*_*T*2_ = 1200*nM*, *M_T_* = 9*nM, p_T_* = 100*nM*. (C) Effect of retroactivity to the output: for the isolated output 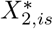, system is simulated with *X*_*T*1_ = 3*nM*, *X*_*T*2_ = 1200*nM*, **M_T_** = 9*nM, p_T_* = 0; for the actual output 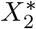, system is simulated with *X*_*T*1_ = 3*nM*, *X*_*T*2_ = 1200*nM*, *M_T_* = 9*nM,p_T_* = 100*nM*.

**Test (iv)** succeeds since the requirements to satisfy Tests (i), (ii) and (iii) do not conflict with each other.

#### 5.6.1 Simulation results for other cascades

##### Phosphotransfer + single cycle

Equations:

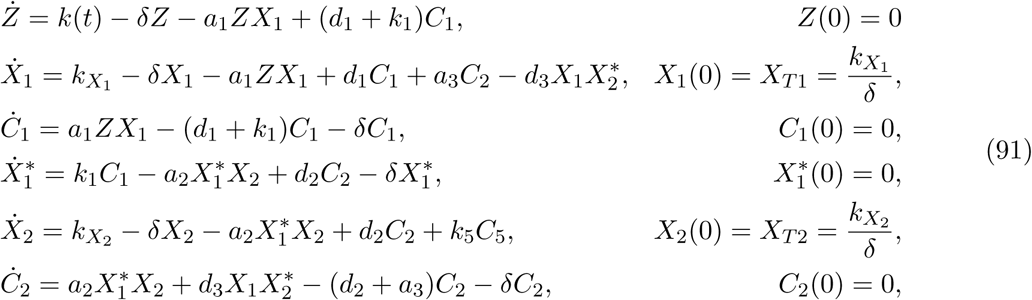

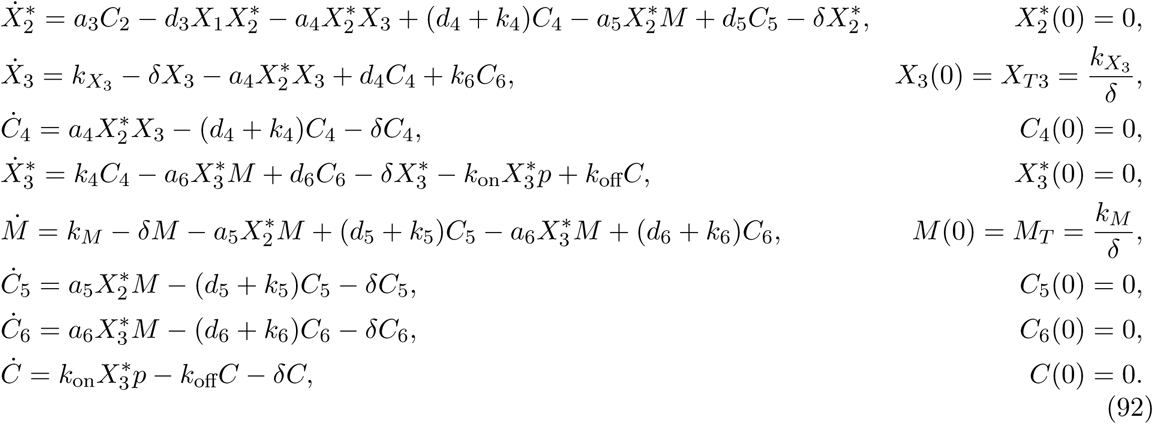

##### Single + Double cycle

Equations:

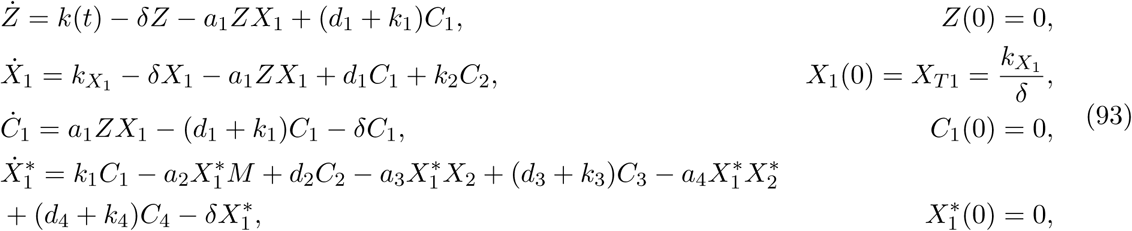

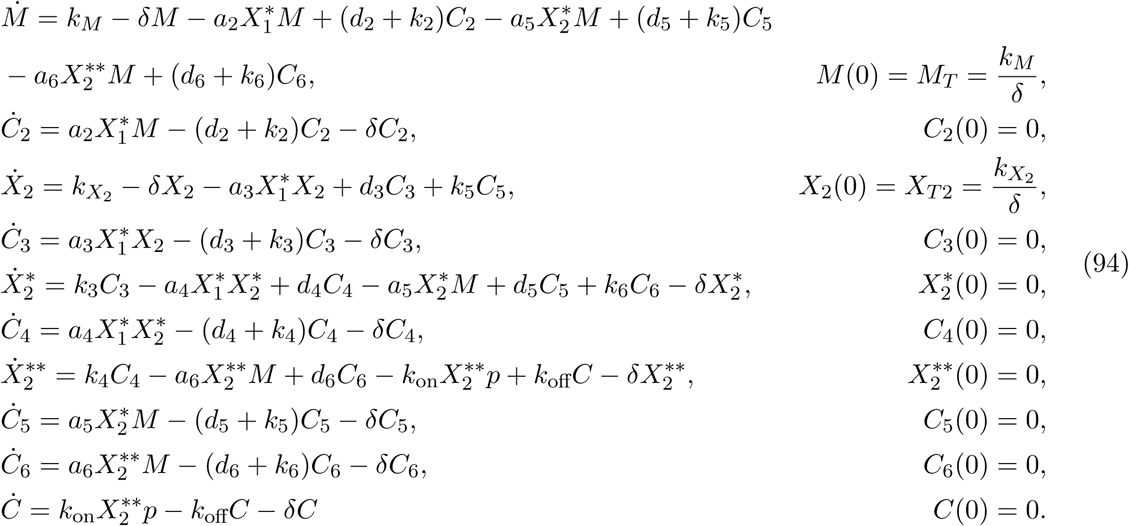

### 5.7 Phosphotransfer with autophosphorylation

The reactions for this system are then:

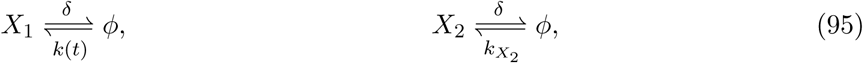

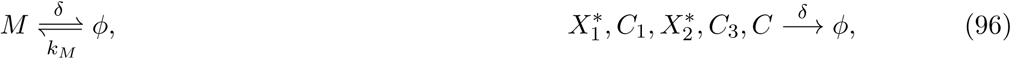

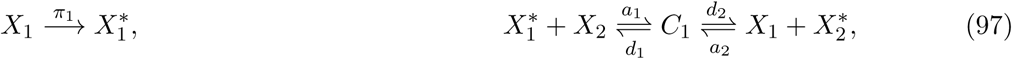

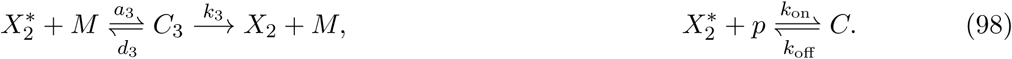

The ODEs based on the reaction rate equations are:

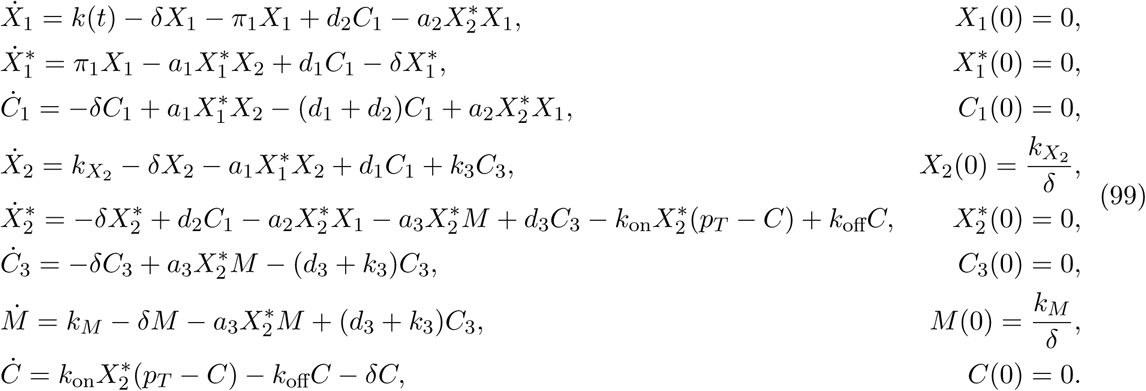

For system (99), define 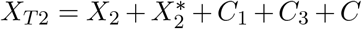 then 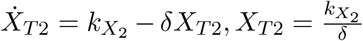. Thus, 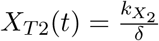 is a constant. Similarly, defining *M_T_* = *M* +*C*_3_ gives a constant 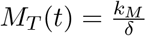. Thus, the variables 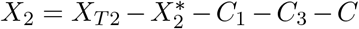 and *M* = *M*_T_ − *C*_3_ can be eliminated from the system. Further, we define 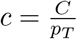. This system is then:

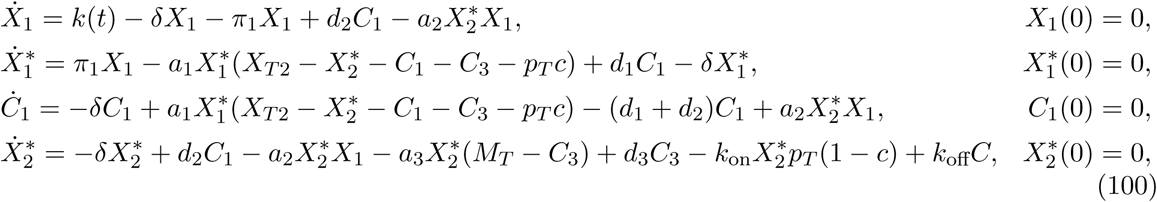

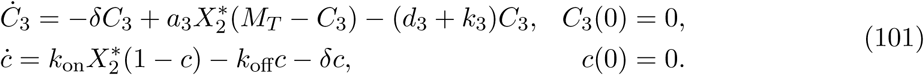

#### Steps 1 and 2

Based on eqns. (100), (101), we bring the system to form (1) as shown in Table 6. We now solve for the functions <u>Ψ</u> and *ϕ* as defined by Assumptions 5 and 6.

**Table 6.** System variables, functions and matrices for a phosphotransfer system with autophosphorylation brought to form (1).

|  |  |  |  |
| --- | --- | --- | --- |
| $U$ | $X_1$ | $v$ | $c$ |
| $\underline{X}$ | $[X_1^* \ C_1 \ X_2^* \ C_3]^T_{4 \times 1}$ | $Y, I$ | $X_2^*, [0 \ 0 \ 1 \ 0]_{1 \times 4}$ |
| $G_1$ | $\max \left\{ \frac{a_1 X_{T2}}{\delta}, \frac{d_1}{\delta}, \frac{d_2}{\delta}, \frac{a_2 X_{T1}}{\delta}, \frac{a_3 M_T}{\delta}, \frac{d_3}{\delta}, \frac{k_3}{\delta} \right\}$ | $G_2$ | $\max \left\{ \frac{k_{\text{on}} p_T}{\delta}, \frac{k_{\text{off}}}{\delta} \right\}$ |
| $f_0(U, R\underline{X}, S_1 v, t)$ | $k(t) - \delta X_1 - \delta C_1 - \delta X_1^*$ | $s(\underline{X}, v)$ | $\frac{1}{G_2} (k_{\text{on}} X_2^* (1 - c) - k_{\text{off}} c - \delta c)$ |
| $\underline{r}(U, \underline{X}, S_2 v)$ | $\frac{1}{G_1} \begin{bmatrix} -\pi_1 X_1 + \delta X_1^*, \\ d_2 C_1 - a_2 X_2^* X_1 + \delta C_1 \end{bmatrix}_{2 \times 1}$ | | |
| $f_1(U, \underline{X}, S_3 v)$ | $\frac{1}{G_1} \begin{bmatrix} -a_1 X_{T2} X_1^* (1 - \frac{C_1}{X_{T2}} - \frac{X_2^*}{X_{T2}} - \frac{C_3}{X_{T2}} - \frac{p_T}{X_{T2}} c) + d_1 C_1, \\ a_1 X_{T2} X_1^* (1 - \frac{C_1}{X_{T2}} - \frac{X_2^*}{X_{T2}} - \frac{C_3}{X_{T2}} - \frac{p_T}{X_{T2}} c) - d_1 C_1, \\ -\delta X_2^* + d_2 C_1 - a_2 X_2^* X_1 + a_3 X_2^* C_3 + d_3 C_3 - a_3 M_T (X_2^* + \frac{p_T \delta}{a_3 M_T} c), \\ -\delta C_3 + a_3 X_2^* (M_T - C_3) - (d_3 + k_3) C_3 \end{bmatrix}_{4 \times 1}$ | | |
| $A$ | $[1 \ 1]_{1 \times 2}$ | $D$ | 1 |
| $B$ | $\begin{bmatrix} -1 & 0 \\ 0 & -1 \\ 0 & 0 \\ 0 & 0 \end{bmatrix}_{4 \times 2}$ | $C$ | $\begin{bmatrix} 0 \\ 0 \\ -p_T \\ 0 \end{bmatrix}_{4 \times 1}$ |
| $R$ | $[1 \ 1 \ 0 \ 0]_{1 \times 4}$ | $S_1$ | 0 |
| $S_2$ | 0 | $S_3$ | $\frac{p_T}{X_{T2}}, \frac{p_T \delta}{a_3 M_T}$ |
| $T$ | $\mathbb{I}_{2 \times 2}$ | $M$ | $[1 \ 1 \ 0 \ 0]_{1 \times 4}$ |
| $Q$ | $\mathbb{I}_{4 \times 4}$ | $P$ | $[0 \ 0 \ p_T \ 0]^T_{4 \times 1}$ |

Solving for *<u>X</u>*= <u>Ψ</u> by setting (*Br* + *f*_1_)_4_ = 0, we have:

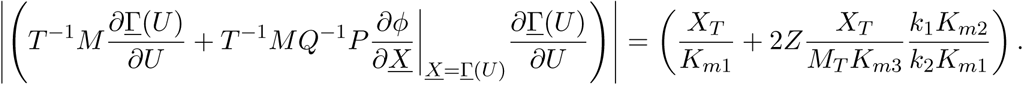

Thus, we have the function <u>Ψ</u>(*U*; *υ*):

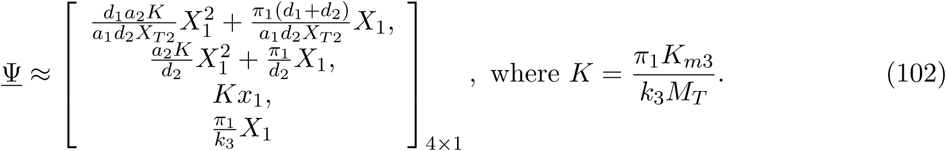

Solving for *ϕ* by setting *s*(*<u>X</u>*; *υ*) = 0, we have:

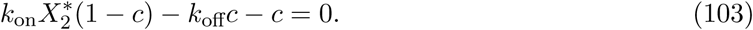

Under Assumption 1, 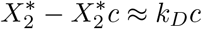,

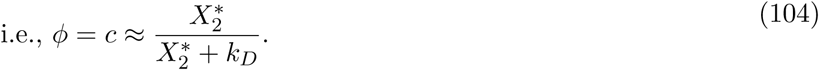

Again, we find <u>Γ</u> from (102) and (104) under Remark 1. This system satisfies Assumptions 3-9. Theorems 1-3 can then be applied.

#### Results: Step 3 and Test (i)

Retroactivity to input: We see that since *S*_1_ = 0 from Table 6. Further 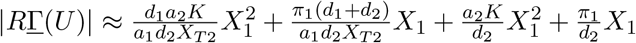 compute the final term, we see that:

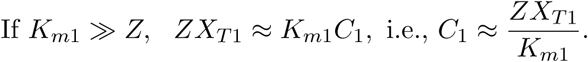

Thus, for a small retroactivity to the input, terms 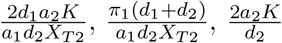 and 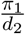 must be small. However, these terms cannot be made smaller by varying concentrations alone. Thus the retroactivity to the input depends on the reaction rate parameters of the system, and is harder to tune.

#### Step 4 and Test

(ii) Retroactivity to output: We see from Table 6 that *S*_1_ = 0, T^−1^*MQ*^−1^*P* = 0, *S*_2_ = 0 and 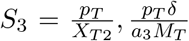. Thus, to attenuate retroactivity to the output, we must have large *XT*_2_ and *M_T_*.

**Step 5 and Test (iii)** Input-output relationship: From (102), we see that

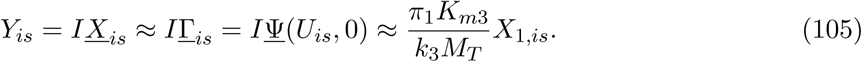

Thus, the dimensionless output 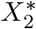 varies linearly with the dimensionless input X_1_, i.e., *m* = 1 and 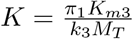

**Test (iv)** is not tested for since Test (i) failed.

### 5.8 Single cycle with substrate input

The reactions for this system are:

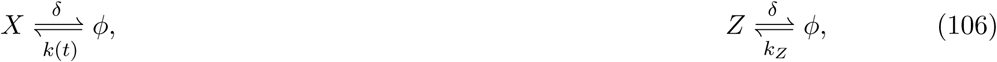

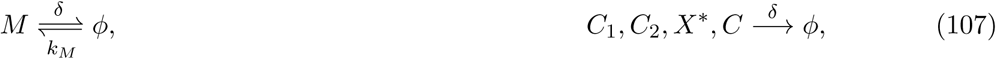

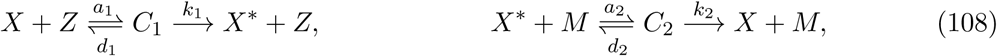

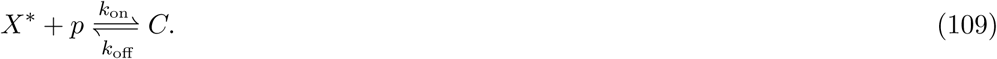

The corresponding ODEs based on the reaction rate equations are then:

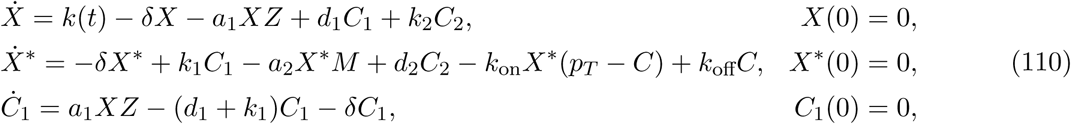

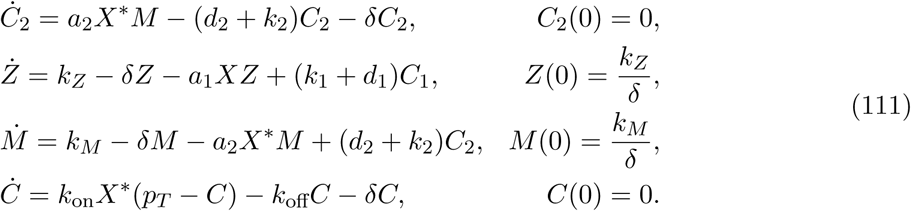

Let *Ζ*_T_ = *Z* + *C*_1_. Then, from the ODEs (110), (111) and the initial conditions, we see that 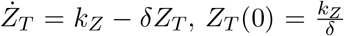. Thus, 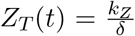 is a constant. Similarly, defining *M_T_* = *M* + *C*_2_ gives a constant 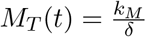. The variables *Z* = *Z*_T_ − *C*_1_ and *M* = *M_T_* − *C*_2_ can then be eliminated from the system. Further, we define 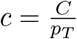. The reduced system is then:

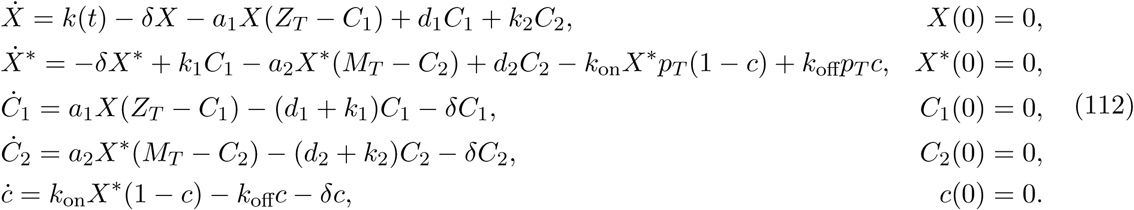

#### Steps 1 and 2

Based on the system of ODEs (112), we bring this system to form (1) as shown in Table 7. We now solve for the functions <u>Ψ</u> and *ϕ* as defined by Assumptions 5 and 6. Solving for *<u>X</u>* = <u>Ψ</u> by setting (*Br* + *f*_1_)_3×1_ = 0, we have:

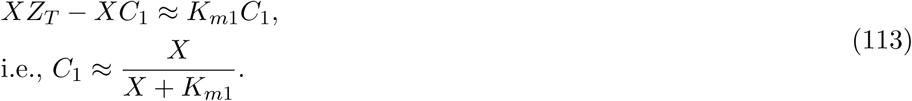

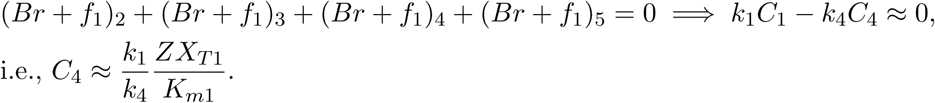

since (*d_2_ + k_2_*) ≫ *δ* under Assumption 1,

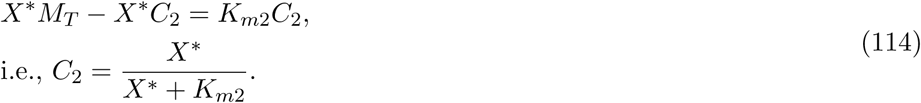

**Table 7.** System variables, functions and matrices for a single phosphorylation cycle with substrate as input brought to form (1).

|  |  |  |  |
| --- | --- | --- | --- |
| $U$ | $X$ | $v$ | $c$ |
| $\underline{X}$ | $[X^* \ C_1 \ C_2]_{3 \times 1}^T$ | $Y, I$ | $X^*, [1 \ 0 \ 0]_{1 \times 3}$ |
| $G_1$ | $\max \left\{ \frac{a_1 Z_T}{\delta}, \frac{d_1}{\delta}, \frac{k_1}{\delta}, \frac{a_2 M_T}{\delta}, \frac{d_2}{\delta}, \frac{k_2}{\delta} \right\}$ | $G_2$ | $\max \left\{ \frac{k_{\text{on}} p_T}{\delta}, \frac{k_{\text{off}}}{\delta} \right\}$ |
| $f_0(U, R\underline{X}, S_1 v, t)$ | $k(t) - \delta X - \delta X^* - \delta C_1 - \delta C_2 - \delta p_T c$ | $s(\underline{X}, v)$ | $\frac{1}{G_2} (k_{\text{on}} X^* (1 - c) - k_{\text{off}} c - \delta c)$ |
| $\underline{r}(U, \underline{X}, S_2 v)$ | $\frac{1}{G_1} [\delta(X^* + p_T c), -a_1 X(Z_T - C_1) + d_1 C_1 + \delta C_1, k_2 C_2 + \delta C_2]_{3 \times 1}^T$ | | |
| $f_1(U, \underline{X}, S_3 v)$ | $\frac{1}{G_1} [k_1 C_1 - a_2 X(M_T - C_2) + d_2 C_2, -k_1 C_1, a_2 X^*(M_T - C_2) - d_2 C_2]_{3 \times 1}^T$ | | |
| $A$ | $[1 \ 1 \ 1]_{1 \times 3}$ | $D$ | 1 |
| $B$ | $\begin{bmatrix} -1 & 0 & 0 \\ 0 & -1 & 0 \\ 0 & 0 & -1 \end{bmatrix}_{3 \times 3}$ | $C$ | $\begin{bmatrix} -p_T \\ 0 \\ 0 \end{bmatrix}_{3 \times 1}$ |
| $R$ | $[1 \ 1 \ 1]_{1 \times 3}$ | $S_1$ | $p_T$ |
| $S_2$ | $p_T$ | $S_3$ | 0 |
| $T$ | 1 | $M$ | $[1 \ 1 \ 1]_{1 \times 3}$ |
| $Q$ | $\mathbb{I}_{3 \times 3}$ | $P$ | $[p_T \ 0 \ 0]_{3 \times 1}^T$ |

Using (113) and (114), we have: 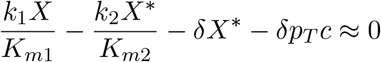,

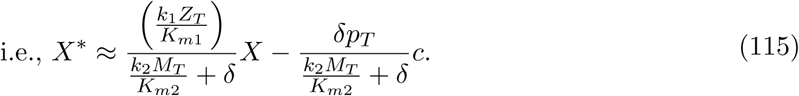

Thus, from equations (113)-(115), we have the function <u>Ψ</u>(*U*; *υ*):

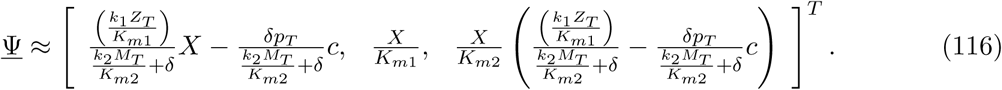

Solving for *υ* = *ϕ*(*<u>X</u>*) by setting *s*(*<u>X</u>*; *υ*) = 0, we have:

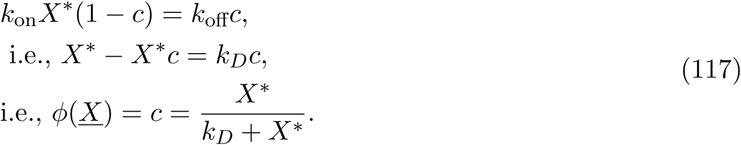

Using (116) and (117), <u>Γ</u> can be found as described in Remark 1. We find that this satisfies Assumption 7. We then state the following claims without proof for this system:

##### Claim 5

*For the matrix B and functions r, f*_1_ *and s defined in Table 7, Assumption 3 is satisfied for this system.*

##### Claim 6

*For the functions f*_0_ *and <u>r</u> and matrices R, S*_1_ *and A defined in Table 7, and the functions* <u>Γ</u> *and ϕ as found above, Assumption 9 is satisfied for this system.*

For matrices *T, Q, M* and *P* as seen in Table 7, we see that Assumption 4 is satisfied. For functions *f*_0_ and *<u>r</u>* defined in Table 7, Assumption 8 is satisfied. Further, for <u>Ψ</u> and *ϕ* defined by (116) and (117), Assumptions 5, 6 and 7 are satisfied. Thus, Theorems 1, 2 and 3 can be applied to this system.

#### Results: Step 3 and Test (i)

Retroactivity to the input: From Table 7, we see that *R* and *S*_1_ cannot be made small by changing system variables. Therefore, Test (i) fails and retroactivity to the input cannot be made small.

#### Step 4 and Test (ii)

Retroactivity to the output: From Table 7, we see that *S*_1_ and *S*_2_ cannot be made small. Therefore, Test (ii) fails and retroactivity to the output cannot be made small.

#### Step 5 and Test (iii)

Input-output relationship: Using Theorem 3, we see that

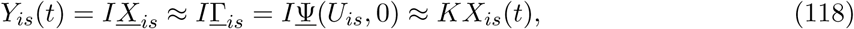

for *t ∈ [t_b_, t_f_]* from (116), where 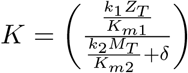.

**Test (iv)** is not tested for since Tests (i) and (ii) failed.

### 5.9 Double cycle with substrate input

The reactions for this system are:

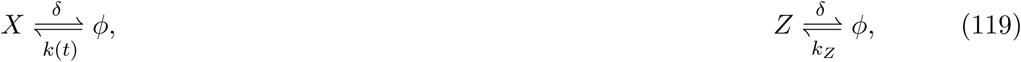

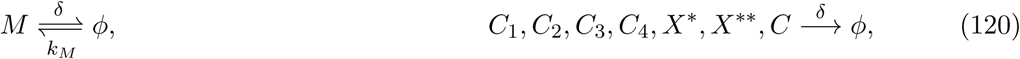

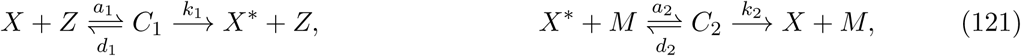

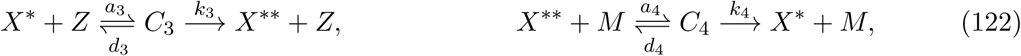

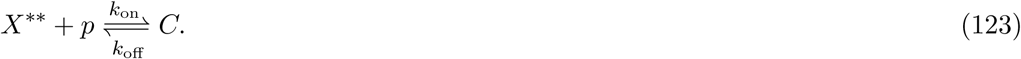

The ODEs based on the reaction rate equations are:

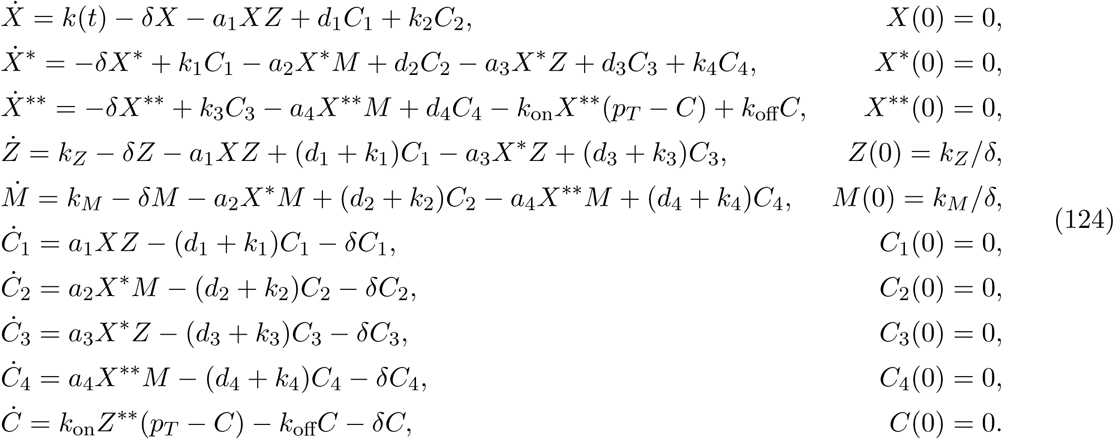

Define *Z_T_* = *Z* + *C*_1_ + *C*_3_. Then, the dynamics of *Z*_T_, seen from (124), are: 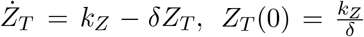 Thus, 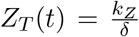 is a constant at all time *t*. Similarly, for *M_T_* = *M* + *C*_2_ + *C*_4_, 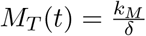 is a constant for all t. Thus, the variables *Z = Z*_T_ *− C*_1_ *− C*_2_ and *M* = *M_T_ − C*_2_ *− C*_4_ can be eliminated from the system. Further, we define 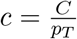. The reduced system is then:

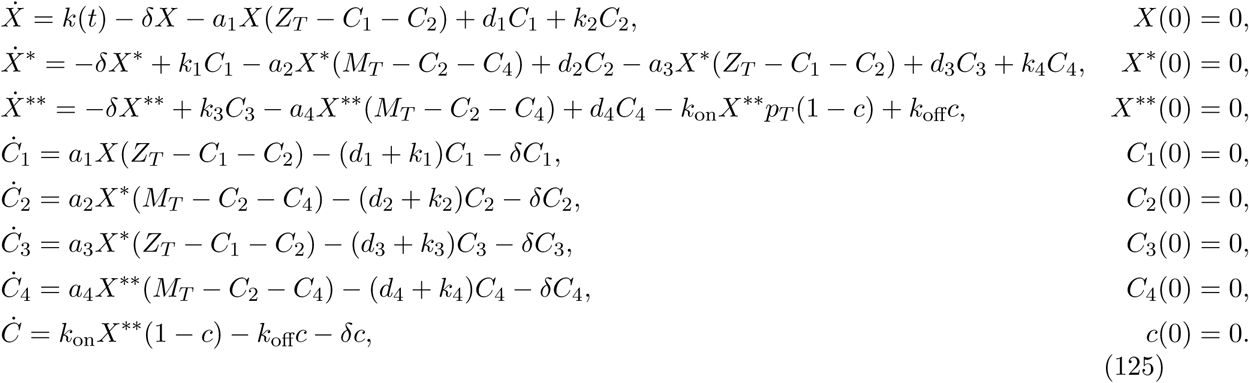

#### Steps 1 and 2

Based on the system of ODEs (125), we bring this system to form (1) as shown in Table 8. We now solve for the functions <u>Ψ</u> and *ϕ* as defined by Assumptions 5 and 6.

**Table 8.** System variables, functions and matrices for a double phosphorylation cycle with substrate as input brought to form (1).

| $U$ | $X$ | $v$ | $c$ |
| --- | --- | --- | --- |
| $\underline{X}$ | $[X^* \ X^{**} \ C_1 \ C_2 \ C_3 \ C_4]^T_{6 \times 1}$ | $Y, I$ | $X^{**}, [0 \ 1 \ 0 \ 0 \ 0 \ 0]_{1 \times 6}$ |
| $G_1$ | $\max \left\{ \frac{a_1 Z_T}{\delta}, \frac{d_1}{\delta}, \frac{k_1}{\delta}, \frac{a_2 M_T}{\delta}, \frac{d_2}{\delta}, \frac{k_2}{\delta}, \frac{a_3 Z_T}{\delta}, \frac{d_3}{\delta}, \frac{k_3}{\delta}, \frac{a_4 M_T}{\delta}, \frac{d_4}{\delta}, \frac{k_4}{\delta} \right\}$ | $G_2$ | $\max \left\{ \frac{k_{\text{on}} p_T}{\delta}, \frac{k_{\text{off}}}{\delta} \right\}$ |
| $f_0(U, R\underline{X}, S_1 v, t)$ | $k(t) - \delta(X + X^* + X^{**} + C_1 + C_2 + C_3 + C_4 + p_T c)$ | $s(\underline{X}, v)$ | $\frac{1}{G_2} (k_{\text{on}} X^{**} (1 - c) - k_{\text{off}} c - \delta c)$ |
| $\underline{r}(U, \underline{X}, S_2 v)$ | $\frac{1}{G_1} [\delta X^*, \delta(X^{**} + p_T c), -a_1 X(Z_T - C_1 - C_3) + d_1 C_1 + \delta C_1, k_2 C_2 + \delta C_2, \delta C_3, \delta C_4]^T_{6 \times 1}$ | | |
| $f_1(u, \underline{x}, S_3 v)$ | $\frac{1}{G_1} \begin{bmatrix} k_1 C_1 - a_2 X^*(M_T - C_2 - C_4) + d_2 C_2 - a_3 X^*(Z_T - C_1 - C_2) + d_3 C_3 + k_4 C_4, \\ k_3 C_3 - a_4 X^{**}(M_T - C_2 - C_4) + d_4 C_4, \\ -k_1 C_1, \\ a_2 X^*(M_T - C_2 - C_4) - d_2 C_2, \\ a_3 X^*(Z_T - C_1 - C_2) - (d_3 + k_3) C_3, \\ a_4 X^{**}(M_T - C_2 - C_4) - (d_4 + k_4) C_4 \end{bmatrix}_{6 \times 1}$ | | |
| $A$ | $[1 \ 1 \ 1 \ 1 \ 1 \ 1]_{1 \times 6}$ | $D$ | $1$ |
| $B$ | $\begin{bmatrix} -1 & 0 & 0 & 0 & 0 & 0 \\ 0 & -1 & 0 & 0 & 0 & 0 \\ 0 & 0 & -1 & 0 & 0 & 0 \\ 0 & 0 & 0 & -1 & 0 & 0 \\ 0 & 0 & 0 & 0 & -1 & 0 \\ 0 & 0 & 0 & 0 & 0 & -1 \end{bmatrix}_{6 \times 6}$ | $C$ | $\begin{bmatrix} 0 \\ -p_T \\ 0 \\ 0 \\ 0 \\ 0 \end{bmatrix}_{6 \times 1}$ |
| $R$ | $[1 \ 1 \ 1 \ 1 \ 1 \ 1]_{1 \times 6}$ | $S_1$ | $p_T$ |
| $S_2$ | $p_T$ | $S_3$ | $0$ |
| $T$ | $1$ | $M$ | $[1 \ 1 \ 1 \ 1 \ 1 \ 1]_{1 \times 6}$ |
| $Q$ | $\mathbb{I}_{6 \times 6}$ | $P$ | $[0 \ p_T \ 0 \ 0 \ 0 \ 0]^T_{6 \times 1}$ |

Solving for *<u>X</u>* = <u>Ψ</u> by setting (*Br* + *f*_1_)_6×1_ = 0, we have:

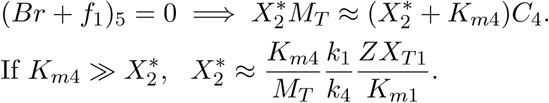

Simultaneously solving these two expressions, for *K*_*m*1_ ≫ *X* and *K*_*m*3_ ≫ *X*^*^:

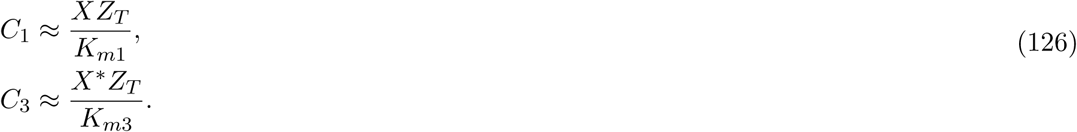

Simultaneously solving these two expressions, for *K*_*m*2_ ≫ *X*^*^ and *K*_*m*4_ ≫ *X*^**^:

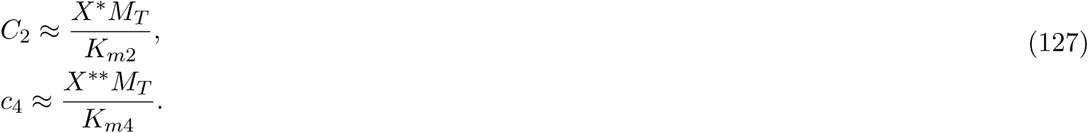

From (126), (127) and (128), 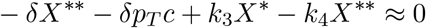

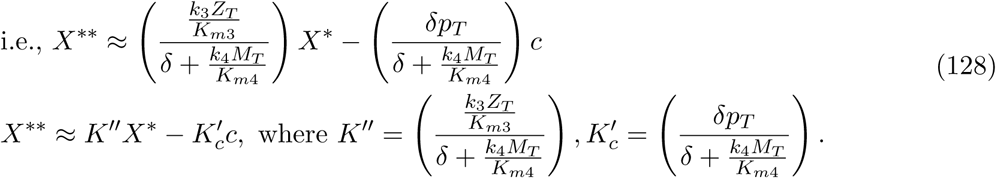

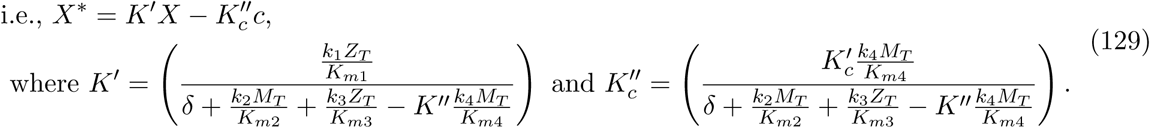

Thus, from equations (126)-(129), for *K*’, *K*”, *K*’_c_ and *K*”_c_ defined in (128) and (129), we have the function <u>Ψ</u>(*U*; *υ*):

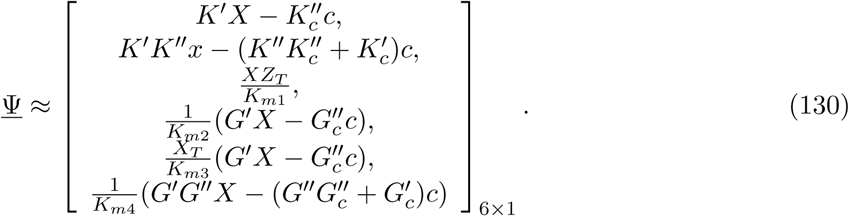

Solving for *ϕ* by setting *s*(*<u>X</u>*; *υ*) = 0, we have:

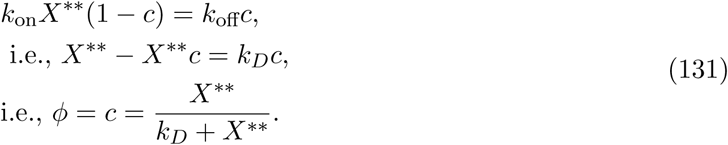

Here again, we find <u>Γ</u> from (130) and (131) under Remark 1, and find that it satisfies Assumption 7. We then state without proof the following claims for this system:

##### Claim 7

*For the matrix B and functions r, f*_1_ *and s defined in Table 8, Assumption 3 is satisfied for this system.*

##### Claim 8

*For the functions f*_0_ *and <u>r</u> and matrices R*, *S*_1_ *and A defined in Table 8, and the functions <u>γ</u> and ϕ as found above, Assumption 9 is satisfied for this system.*

For matrices *T*; *Q*, *M*, *P* defined in Table 8, we see that Assumption 4 is satisfied. Further, for <u>Ψ</u> and *ϕ* defined by (130) and (131), Assumption 5 and 6 are satis_ed. Thus, Theorems 1, 2 and 3 can be applied to this system.

#### Results: Step 3 and Test (i)

Retroactivity to the input: From Table 8, we see that *R* and *S*_1_ cannot be made small. Thus, Test (i) fails and retroactivity to the input cannot be made small.

#### Step 4 and Test (ii)

Retroactivity to the output: From Table 8, *S*_1_ and *S*_2_ cannot be made small. Thus, Test (ii) fails and retroactivity to the output cannot be made small.

#### Step 5 and Test (iii)

Input-output relationship: From (130),

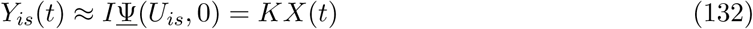

for *t ∈* [*t_b_*, *t_f_*]. Thus the input-output relationship has *m* = 1 and *K* = KK as defined in (128), (129), which can be tuned by tuning the total kinase and phosphatase concentrations *Z_T_* and *M_T_*.

**Test (iv)** is not tested for since Tests (i) and (ii) failed

